# Does the strength of women’s attraction to male vocal masculinity track changes in steroid hormones?

**DOI:** 10.1101/403949

**Authors:** Benedict C Jones, Amanda C Hahn, Katarzyna Pisanski, Hongyi Wang, Michal Kandrik, Anthony J Lee, Iris J Holzleitner, David R Feinberg, Lisa M DeBruine

## Abstract

Recent studies that either used luteinizing hormone tests to confirm the timing of ovulation or measured steroid hormones from saliva have found little evidence that women’s preferences for facial or body masculinity track within-subject changes in women’s fertility or hormonal status. Fewer studies using these methods have examined women’s preferences for vocal masculinity, however, and those that did report mixed results. Consequently, we used a longitudinal design and measured steroid hormones from saliva to test for evidence of hormonal regulation of women’s (N=351) preferences for two aspects of male vocal masculinity (low pitch and low formants). Analyses suggested that preferences for masculine pitch, but not masculine formants, may track within-woman changes in estradiol. Although these results present some evidence for the hypothesis that within-subject hormones regulate women’s attraction to masculine men, we do not discount the possibility that the effect of estradiol on pitch preferences in the current study is a false positive.

## Introduction

The Dual Mating Strategy hypothesis of ovulatory shifts in women’s mate preferences proposes that women show stronger preferences for masculine men during high-fertility ovulatory phase of their menstrual cycle (Gangestad & Simpson, 2000; Little et al., 2011; Penton-Voak et al., 1999). Early tests of this hypothesis suggested women do indeed show stronger preferences for masculine characteristics in men’s faces (Little & Jones, 2012; Penton-Voak et al., 1999), voices (Puts, 2005, 2006; Feinberg et al., 2006), bodies (Little et al., 2007), behavioral displays (Gangestad et al., 2004), and body odors (Havlicek et al., 2005). However, these studies have recently been criticized for employing small sample sizes, using weak methods to assess women’s position in the menstrual cycle and/or hormonal status, and being overly-reliant on between-subject designs (Blake et al., 2016; Gangestad et al., 2016; Jones et al., 2018a).

These criticisms have triggered a concerted effort from many research groups to test for fertility- and-or hormone-linked changes in women’s masculinity preferences using methods that address the methodological limitations of earlier work. Recent large-scale studies of women’s preferences for masculinity in men’s faces (Jones et al., 2018a; Marcinkowska et al., 2018a; Marcinkowska et al., 2018b) and bodies (Jünger et al., 2018a; Marcinkowska et al., 2018a; Marcinkowska et al., 2018b) that used luteinizing hormone tests and/or measured women’s hormone levels found no clear evidence that face or body preferences tracked within-subject changes in women’s fertility or hormonal status. One exception to this pattern was Dixson et al. (2018), who found that women’s preferences for facial masculinity and femininity tracked changes in estradiol and progesterone, respectively. Dixson et al. (2018) did not find significant correlations between face preferences and fertility, however.

Recent studies of women’s preferences for vocal masculinity using similar designs to those described above have reported more mixed results, however. One study found that women’s preferences for vocal masculinity were slightly stronger in test sessions where women had higher salivary estradiol, although this effect was not significant^1^ (Pisanski et al., 2014). By contrast, two other recent studies found no significant effects of fertility on women’s vocal masculinity preferences and found no evidence that women’s vocal masculinity preferences were reliably with steroid hormone levels (Jünger et al., 2018b).

In light of the mixed results for vocal masculinity described above, here we report new data on the hormonal correlates of within-women changes in preferences for vocal masculinity. Following Pisanski et al. (2014), we used two different types of masculinity preference test. One test assessed women’s preferences for masculinized versus feminized voice pitch. The other test assessed women’s preferences for voices that had been masculinized and feminized in formant frequencies. Both pitch and format frequencies are reliably sexually dimorphic in young adults (Pisanski et al., 2016).

## Methods

### Participants

We tested 371 heterosexual women (mean age=21.6 years, SD=3.31 years) who reported that they were not using any form of hormonal contraceptive (i.e., all women reported having natural menstrual cycles). Women participated as part of a larger study investigating possible effects of steroid hormones on different aspects of women’s behavior (Jones et al., 2018a, 2018b, 2018c). In this larger study, participants completed up to three blocks of test sessions. Each of the three blocks of test sessions consisted of five weekly test sessions. The data reported here are all responses from blocks of test sessions in the larger study where women provided voice preference data in at least one test session. Following these restrictions, 328 women had completed five or more test sessions and 96 of these women completed ten test sessions. Forty-three women completed fewer than five test sessions.

Sixty-two of our participants’ first block of test sessions was previously reported in Pisanski et al. (2014). Analyses are reported with and without these data points.

### Stimuli

Voice stimuli were identical to those used in Kandrik et al. (2016) and Pisanski et al. (2014). Recordings of 6 men between the ages of 18 and 25 speaking the English monopthong vowels, “ah”/α/, “ee”/i/, “e”/ε/, “oh”/o/, and “oo”/u/, were made in an anechoic sound-controlled booth using a Sennheiser MKH 800 cardioid condenser microphone (MKH-P48), at an approximate distance of 5-10 cm. Voice recordings were digitally encoded using an M-Audio Fast Track Ultra interface at a sampling rate of 96 kHz and 32-bit amplitude quantization, and transferred to a computer as PCM WAV files using Adobe Soundbooth CS5 version 3.0.

Following other recent work on perceptions of sexually dimorphic vocal characteristics (e.g., Kandrik et al., 2016; Pisanski et al., 2014), we created two feminized and two masculinized versions of each original voice recording by independently manipulating voice pitch or formants using the Pitch-Synchronous Overlap Add (PSOLA) algorithm in Praat version 5.2.15 (Boersma & Weenink, 2013). Pitch was lowered (masculinized) or raised (feminized) by 10% from baseline while holding formants constant. Likewise, formants were lowered (masculinized) or raised (feminized) by 10% from baseline while holding pitch constant. This process created 6 pairs of voices that differed in pitch and 6 pairs of voices that differed in formants. Following these manipulations, we amplitude-normalized the sound pressure level of all voices to 70 decibels using the root mean squared method. Voice pitch and formant measures for the feminized and masculinized voice stimuli are given in Kandrik et al. (2016). We have previously shown that these voice manipulations reliably alter masculinity and dominance perceptions in the predicted way (i.e., lowered pitch or formants increase relative judgments of masculinity and dominance; Kandrik et al., 2016; Pisanski et al., 2014).

### Vocal masculinity preference test

In each test session, participants listened to 12 pairs of voices (each pair consisting of a masculinized and a feminized version of the same voice) through headphones (Sennheiser HD202). Women were instructed to select the more attractive voice in each pair and to indicate the strength of that preference by choosing from the options “slightly more attractive”, “somewhat more attractive”, more attractive”, and “much more attractive”. Trial order and the order in which participants listened to the masculinized and feminized versions in each pair were fully randomized.

Responses on the masculinity preference test were coded using the following scale (higher scores indicate stronger masculinity preferences and the scale is centered on chance, i.e., zero):

0.5 to 3.5: masculinized voice rated ‘slightly more attractive’ (=0.5), ‘somewhat more attractive’ (=1.5), ‘more attractive’ (=2.5) or ‘much more attractive’ (=3.5) than feminized voice.

−0.5 to −3.5: feminized voice rated ‘slightly more attractive’ (=−0.5), ‘somewhat more attractive’ (=−1.5), ‘more attractive’ (=−2.5) or ‘much more attractive’ (= −3.5) than masculinized voice.

Each woman’s average masculinity preference score was calculated separately for the pitch-manipulated and formant-manipulated trials for each test session. This method for assessing and coding masculinity preferences has been used in many previous studies (e.g., Jones et al., in press a; Zietsch et al., 2015). Higher scores indicate stronger masculinity preferences.

### Saliva samples

Participants provided a saliva sample via passive drool (Papacosta & Nassis, 2011) in each test session. Participants were instructed to avoid consuming alcohol and coffee in the 12 hours prior to participation and avoid eating, smoking, drinking, chewing gum, or brushing their teeth in the 60 minutes prior to participation. Each woman’s test sessions took place at approximately the same time of day to minimize effects of diurnal changes in hormone levels (Veldhuis et al., 1988; Bao et al., 2003).

Saliva samples were frozen immediately and stored at −32°C until being shipped, on dry ice, to the Salimetrics Lab (Suffolk, UK) for analysis, where they were assayed using the Salivary 17β-Estradiol Enzyme Immunoassay Kit 1-3702 (M=3.30 pg/mL, SD=1.27 pg/mL, sensitivity=0.1 pg/mL, intra-assay CV=7.13%, inter-assay CV=7.45%), Salivary Progesterone Enzyme Immunoassay Kit 1-1502 (M=148.45 pg/mL, SD=95.95 pg/mL, sensitivity=5 pg/mL, intra-assay CV=6.20%, inter-assay CV=7.55%), Salivary Testosterone Enzyme Immunoassay Kit 1-2402 (M=87.58 pg/mL, SD=27.18 pg/mL, sensitivity<1.0 pg/mL, intra-assay CV=4.60%, inter-assay CV=9.83%), and Salivary Cortisol Enzyme Immunoassay Kit 1-3002 (M=0.23 µg/dL, SD=0.16 µg/dL, sensitivity<0.003 µg/dL, intra-assay CV=3.50%, inter-assay CV=5.08%).

Hormone levels more than three standard deviations from the sample mean for that hormone, or where Salimetrics indicated levels were outside the sensitivity range of their relevant ELISA, were excluded from the dataset (∼1% of hormone measures were excluded for these reasons). The descriptive statistics given above do not include these excluded values. Values for each hormone were centered on their subject-specific means to isolate effects of within-subject changes in hormones. They were then scaled so the majority of the distribution for each hormone varied from −.5 to .5 to facilitate calculations in the linear mixed models. As hormone levels were centered on their subject-specific means, women with only one value for a hormone could not be included in the analyses.

### Analyses

Linear mixed models were used to test for possible effects of hormonal status on masculinity preferences. Analyses were conducted using R version 3.3.2 (R Core Team, 2016), with lme4 version 1.1-13 (Bates et al., 2014) and lmerTest version 2.0-33 (Kuznetsova et al., 2013). The dependent variable was the masculinity preference score for each test session (separate models were run for the pitch and formant manipulations). Predictors were the scaled and centered hormone levels. Random slopes were specified maximally following Barr et al. (2013) and Barr (2013). Full model specifications and full results for each analysis are given in our Supplemental Materials. Data files and analysis scripts are publicly available at https://osf.io/byu3h/. 95% confidence intervals are given in our Supplemental Materials.

## Results

Masculinity preferences for pitch- or formant-manipulated voice stimuli were analyzed separately. For each type of masculinity manipulation (pitch manipulation, formant manipulation) we ran three models. Our first model (Model 1) included estradiol, progesterone, and their interaction as predictors. Our second model (Model 2) included estradiol, progesterone, and estradiol-to-progesterone ratio as predictors. Our third model (Model 3) included testosterone and cortisol as predictors, but did not consider possible effects of estradiol or progesterone. This analysis strategy is identical to that used in Jones et al. (2018a), Jones et al. (2018b), and Jones et al. (2018c) to investigate the hormonal correlates of women’s facial masculinity preferences, sexual desire, and disgust sensitivity, respectively.

### Preferences for masculinized pitch

The intercept was significant in all three models (all estimates>0.70, all ts>26.60, all ps<.001, M=0.70, SEM=0.03), indicating that women generally judged pitch-masculinized versions of male voices to be more attractive than pitch-feminized versions. Model 1 revealed a significant positive effect of estradiol (estimate=0.19, t=2.38, p=.017), but neither the effect of progesterone (estimate=-0.04, t=-0.60, p=.55) nor the interaction between estradiol and progesterone (estimate=-0.04, t=-0.10, p=.92) were significant. Model 2 also revealed a significant positive effect of estradiol (estimate=0.21, t=2.58, p=.010), but neither the effects of progesterone (estimate=-0.09, t= −1.15, p=.25) nor estradiol-to-progesterone ratio (estimate=-0.05, t=-1.23, p=.22) were significant. In Model 3, neither testosterone (estimate=0.15, t=1.77, p=.08) nor cortisol (estimate=-0.06, t=-0.86, p=.39) had a significant effect on women’s pitch preferences.

### Preferences for masculinized formants

The intercept was significant in all three models (all estimates>0.44, all ts>13.08, all ps<.001, M=0.45, SEM=0.03), indicating that women generally judged formant-masculinized versions of male voices to be more attractive than formant-feminized versions. None of the three models testing for effects of hormones on formant-masculinized voices revealed significant effects of any hormones (all absolute estimates<0.18, all absolute ts<0.46, all ps>.50).

Repeating the analyses of pitch and formant preferences described above including women’s partnership status (partnered versus unpartnered) as an additional effect-coded factor revealed no significant interactions involving partnership status. The significant effects of estradiol on pitch preferences described above were also significant in these analyses. These analyses are described in full in our Supplemental Materials.

### Additional within-subject analyses

An analysis of voice preference data from 62 of our participants’ first block of test sessions was previously reported in Pisanski et al. (2014). Excluding these data from our analyses did not alter the pattern of significant results. Full results of these additional analyses are reported in our Supplemental Materials.

### Between-subject analyses

Recent work on links between between-subject (i.e., average) progesterone levels and face (Marcinkowska et al., 2018a; DeBruine et al., 2018) and body (Marcinkowska et al., 2018a) preferences suggest that masculinity preferences are predicted by the combined effects of relationship status and average progesterone. The corresponding effects for pitch and formant preferences in this data set (all participants included) were not significant. These results are described in full in our Supplemental Materials.

## Discussion

Consistent with previous research (e.g., Feinberg et al., 2005; Pisanski et al., 2014), women in our study showed strong preferences for male voices with masculinized pitch and formants over those with feminized pitch and formants, respectively. We also found that women’s preferences for masculinized pitch, but not masculinized formants, were stronger in test sessions with higher estradiol. These results differ from those reported in other recent work on fertility- and hormone-regulated voice preferences in important ways.

First, and despite using identical stimuli to Pisanski et al. (2014), we found that only pitch preferences tracked changes in women’s estradiol. This contrasts with Pisanski et al. (2014), who reported that vocal masculinity preferences tended to track changes in estradiol regardless of whether they had been manipulated in pitch or formants. Since masculine formants are reliably associated with masculine body shapes (Pisanski et al., 2016), our null results for formant preferences are arguably consistent with recent work finding no within-subject effects of hormonal status on women’s preferences for masculine body characteristics (Jünger et al., 2018a).

Second, our results differ from those reported in Jünger et al. (2018b). Across two studies, Jünger et al. (2018b) found no evidence that women’s preferences for vocal masculinity tracked changes in either fertility or estradiol. Jünger et al’s (2018b) results suggest that the within-subject effect of estradiol observed in the current study is not robust.

In conclusion, the current study saw some evidence that women’s preferences for one component of vocal masculinity (pitch), but not another (formants), track changes in estradiol. This is surprising since there is no a priori reason to expect hormonal effects for only one component of vocal masculinity. Although these results partially replicate those of a previous study using identical methods (Pisanski et al., 2014), the evidence is not particularly strong and they conflict with other recent work finding no evidence for hormonal regulation of vocal masculinity preferences (Jünger et al., 2018b). Thus, we do not rule out the possibility that the effect of estradiol on pitch preferences is a false positive. Indeed, the effects of estradiol on pitch preferences would not be significant if corrected for multiple comparisons. Arguably, more work is needed to investigate the reliability and robustness of hormonal effects on women’s vocal masculinity preferences.

## Women’s preferences for masculine pitch in men’s voices may track changes in estradiol

*Jones et al.*

## Setup and data processing

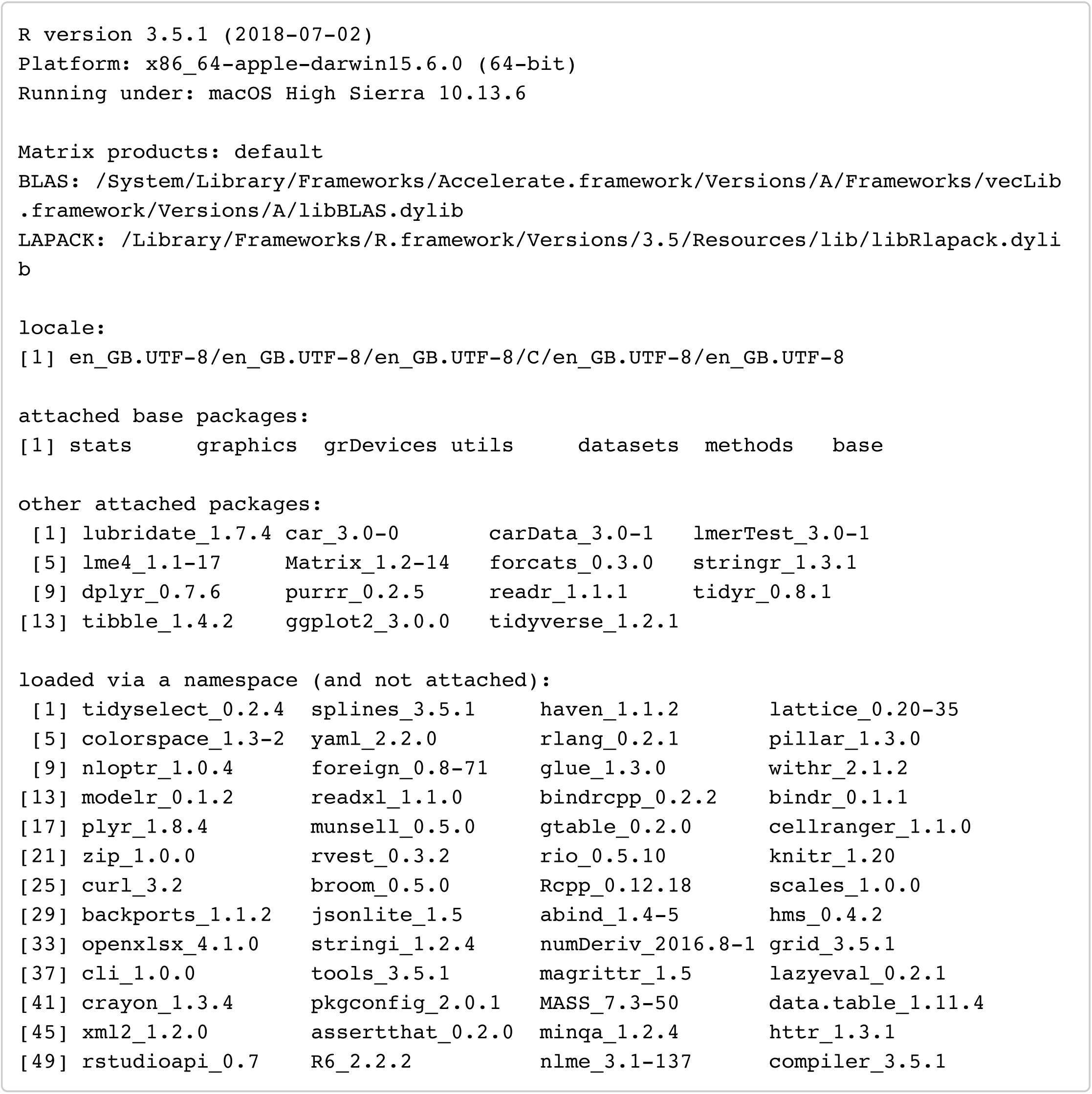

### Custom Functions

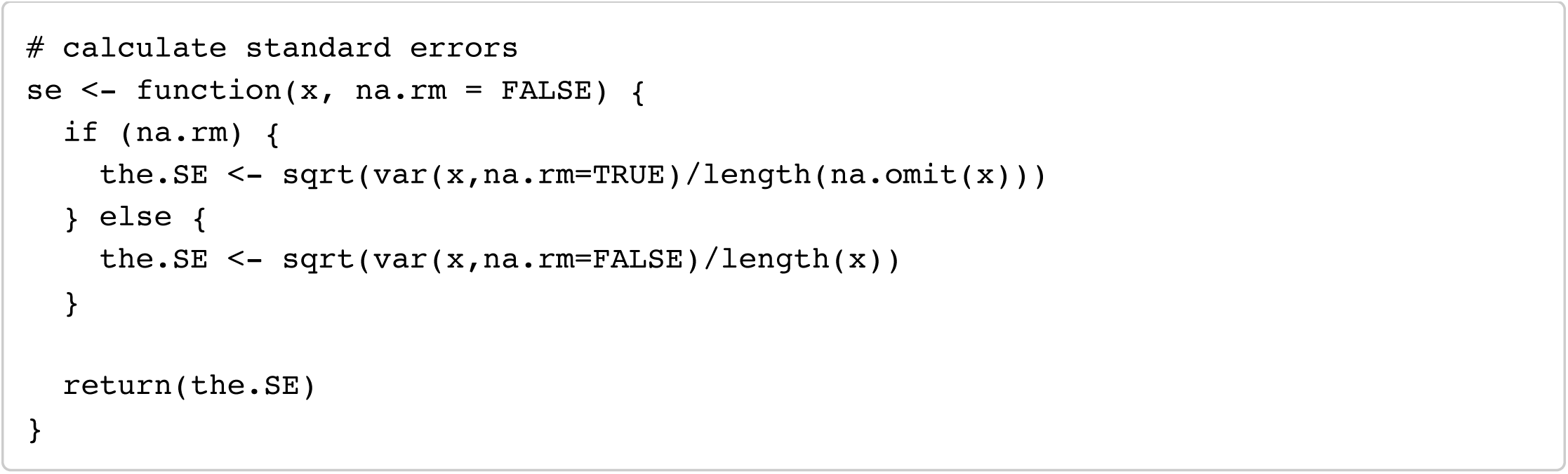

### Load full dataset

Data entered from all white, heterosexual women not using hormonal contraceptives.

Each row is all data from a single session (i.e. oc_id:date)

- “oc_id” = ID of the subject
- “age” = age (in years) of subject on day of testing
- “date” = date of testing session
- “block” = testing block (1, 2 or 3)
- “block_partner” = partnership status (0 = no partner, 1 = partner)
- “manip” = type of masculinity maipulation (formant or pitch)
- “rating.c” = mean voice rating (centered on chance)
- “prog” = salivary progesterone for that session
- “estr” = salivary estradiol for that session
- “test” = salivary testosterone for that session
- “cort” = salivary cortisol for that session
- “partner.e” = effect-coded partnership status (−0.5 = no partner, 0.5 = partner)

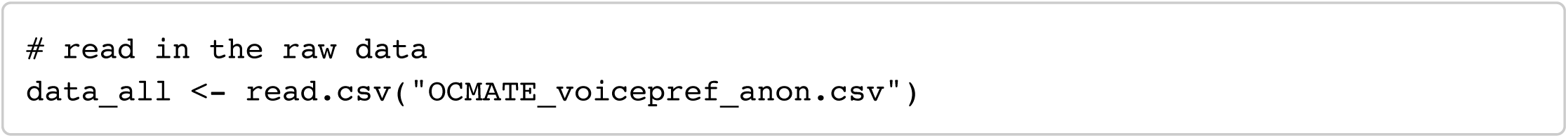

#### Age

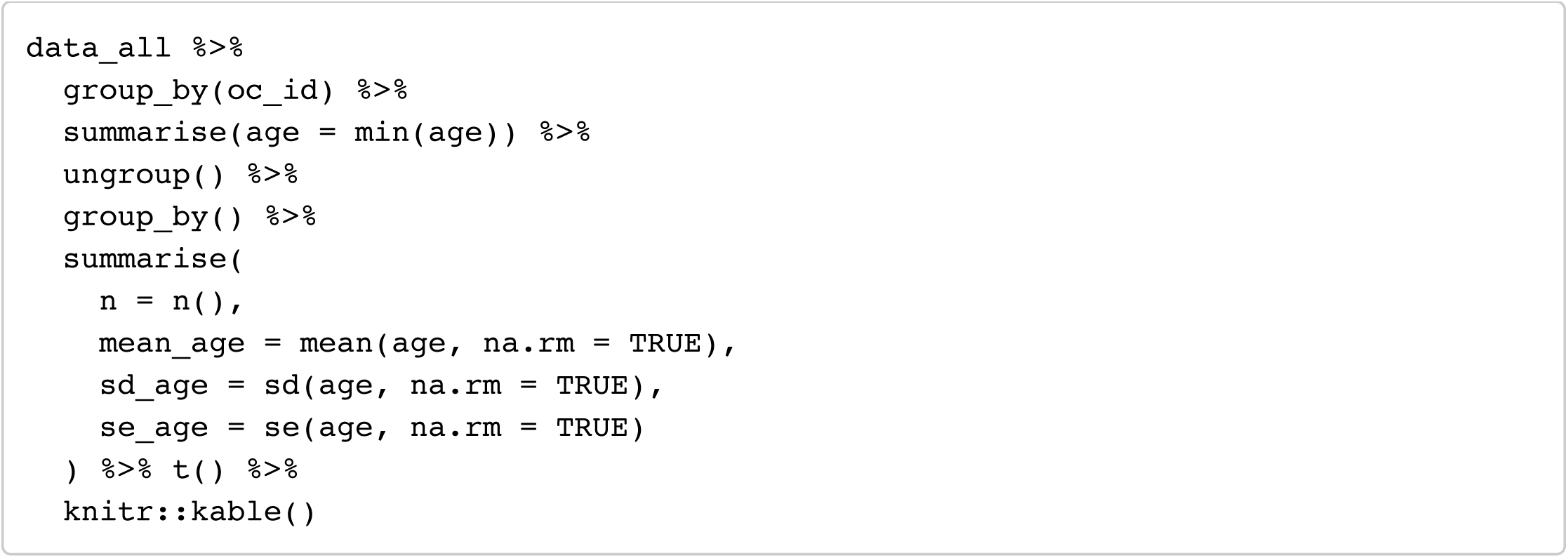

The table should have a header (column names)

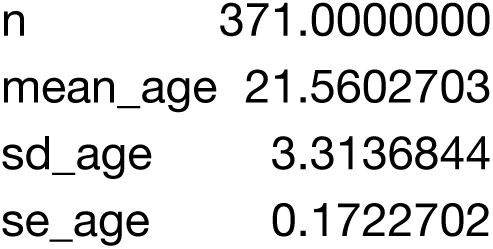

#### The number of sessions completed per woman

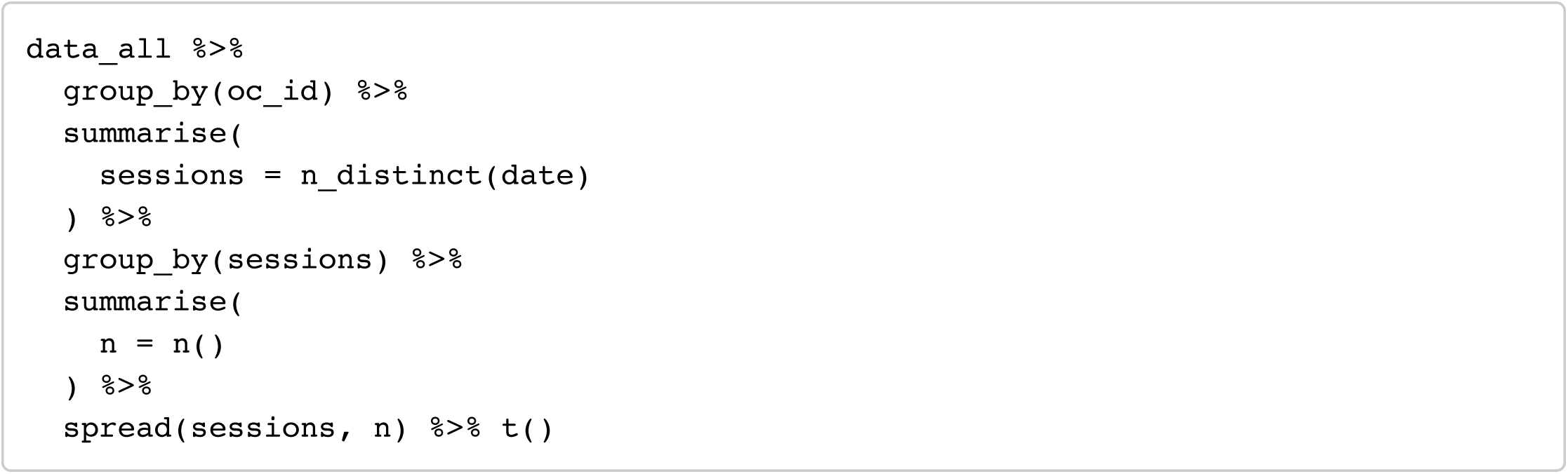

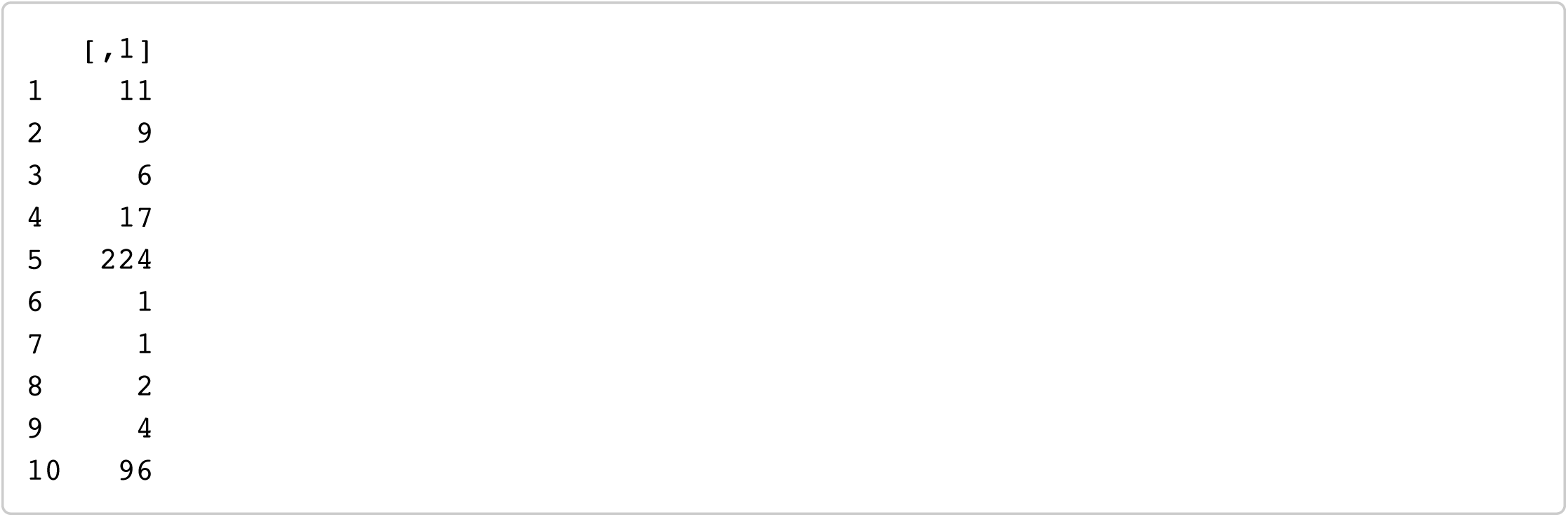

### Exclude observations with missing estradiol, progesterone or testosterone

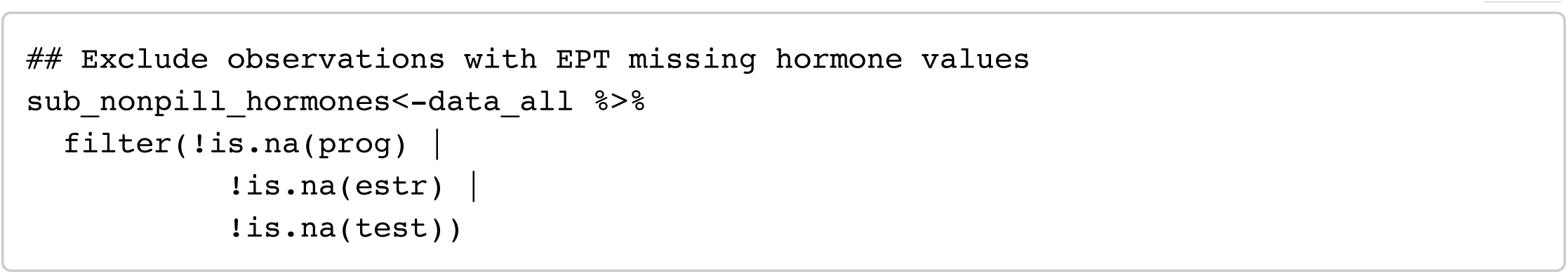

### Exclude subjects with only a single session in a block

This is necessary because you can’t calculate subject-centered means with only one data point.

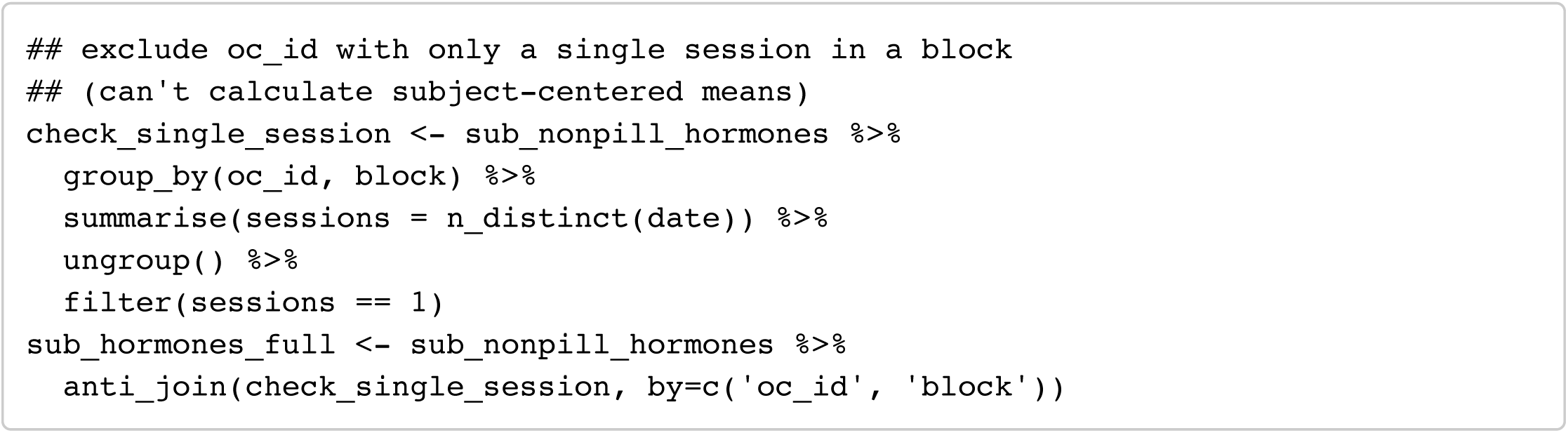

### Remove outlier hormone values

Remove below bottom sensitivity thresholds for assays (prog < 5, estr < 0.1) and remove outlier values (+/-3SD from the mean).

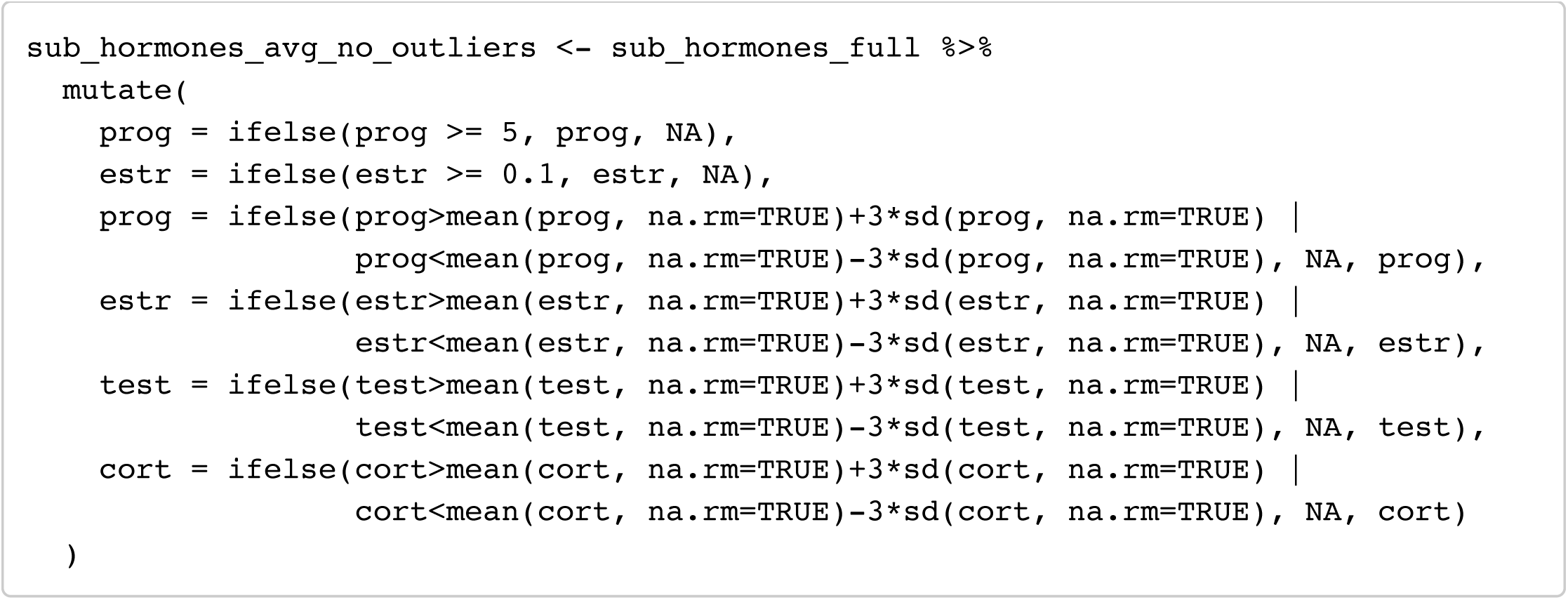

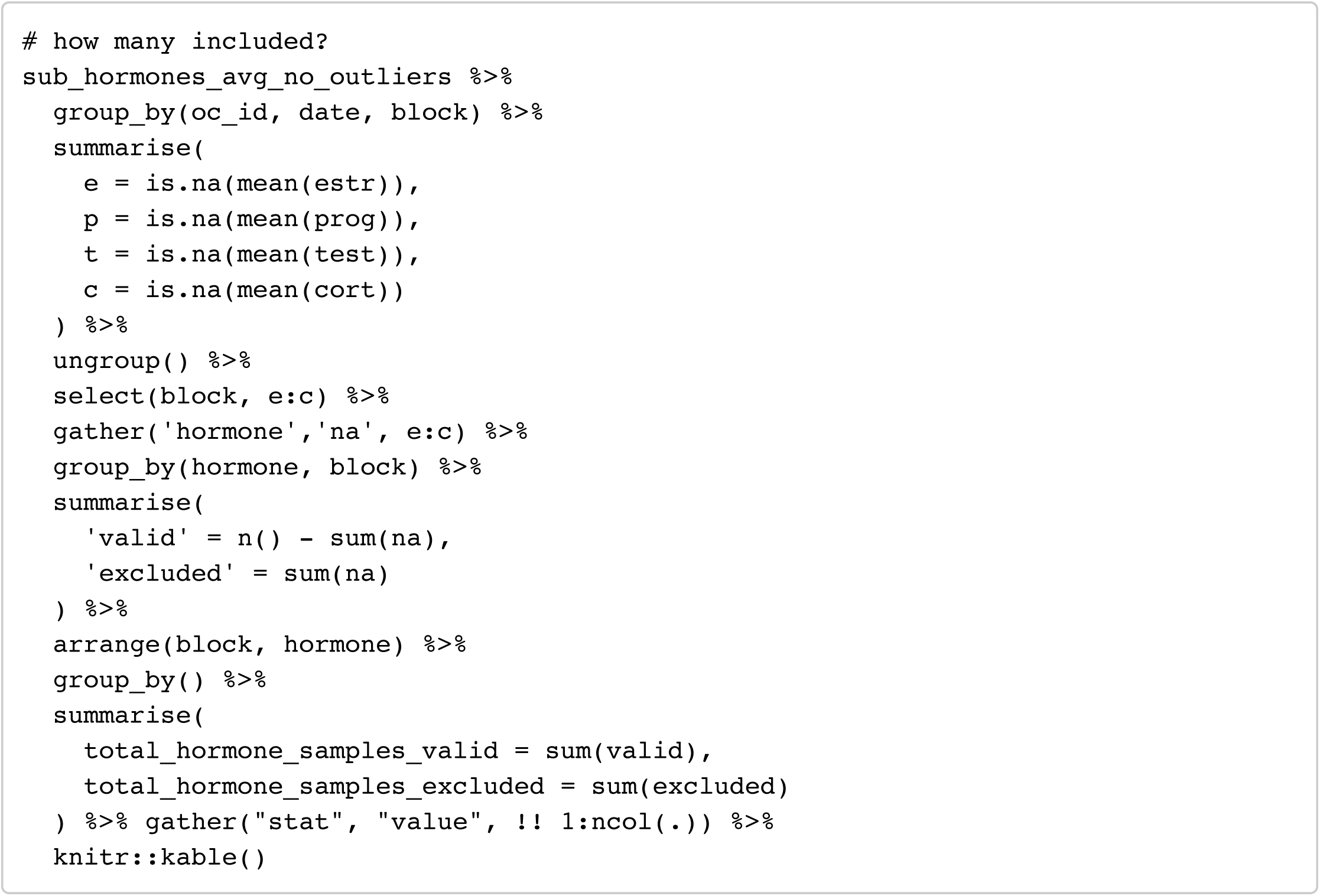

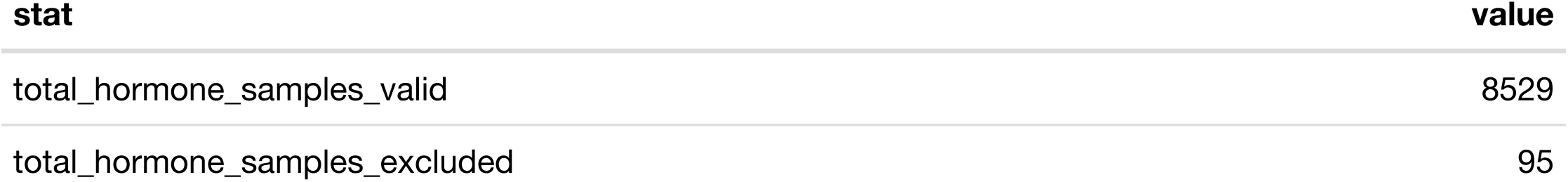

### Subject-mean-centre hormones

Divide results by a constant to put all hormones on ∼ −0.5 to +0.5 scale

Hide

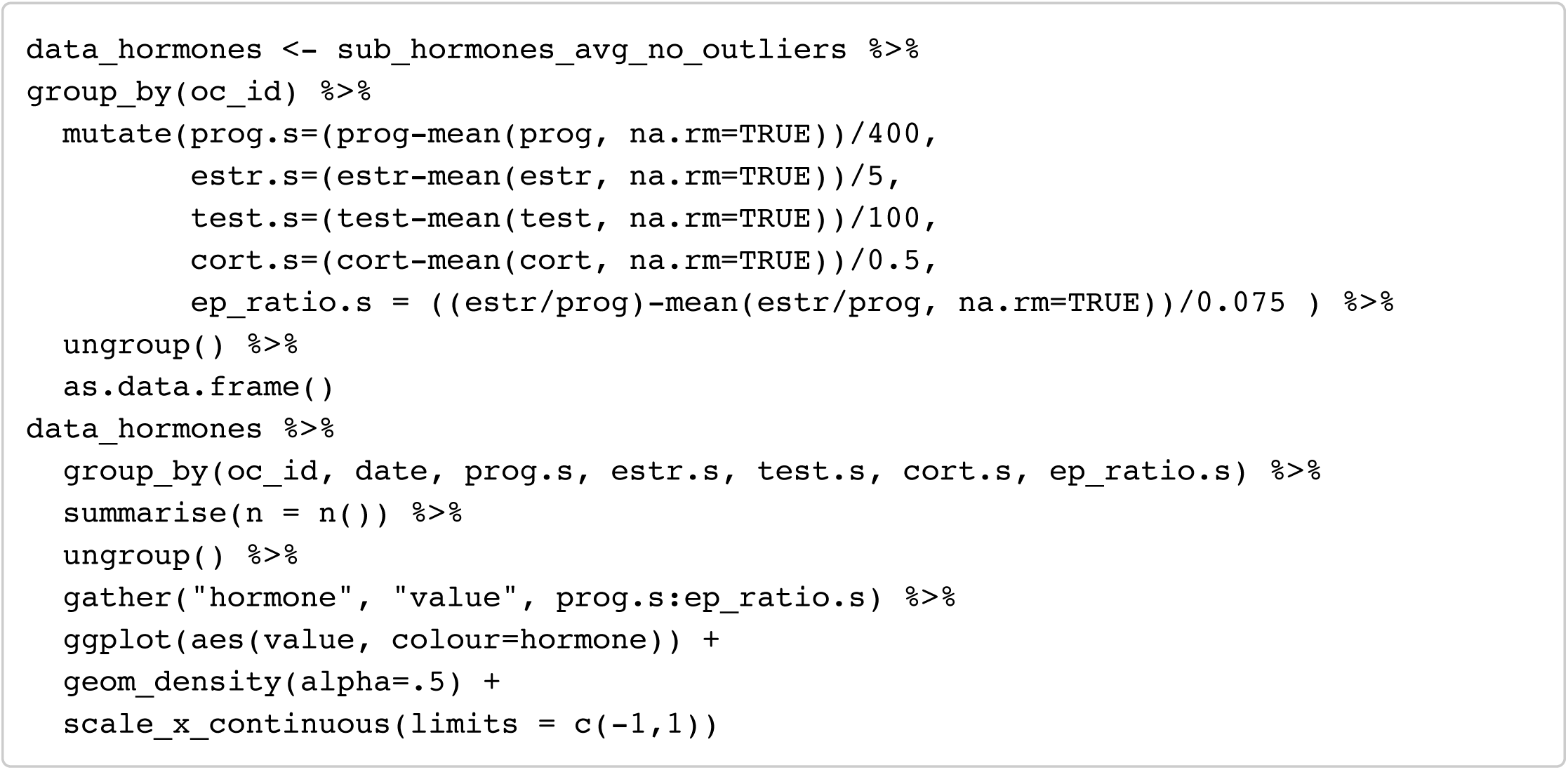

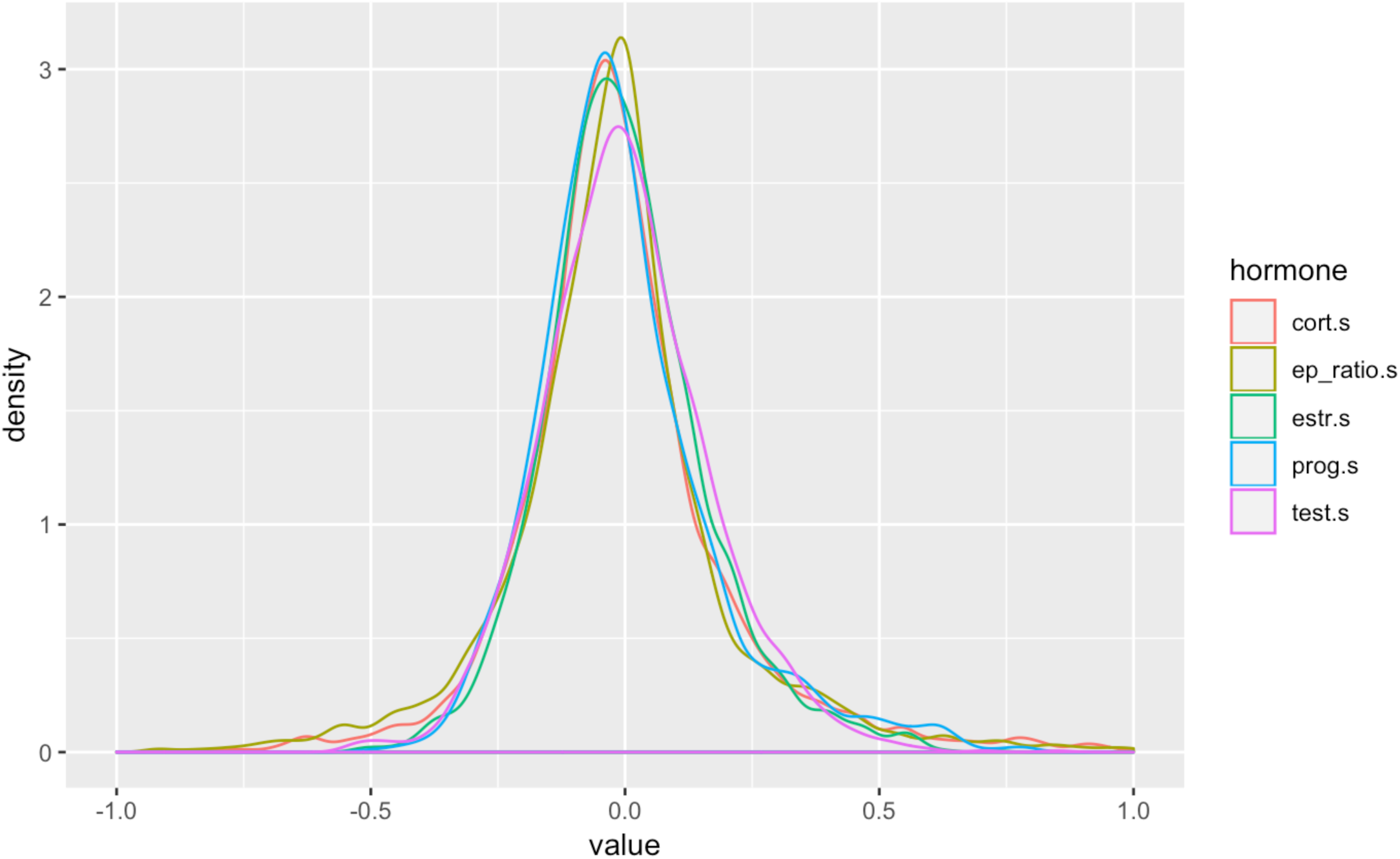

Hide

NA

## Descriptive stats

### Voice Masculinity preferences

Hide

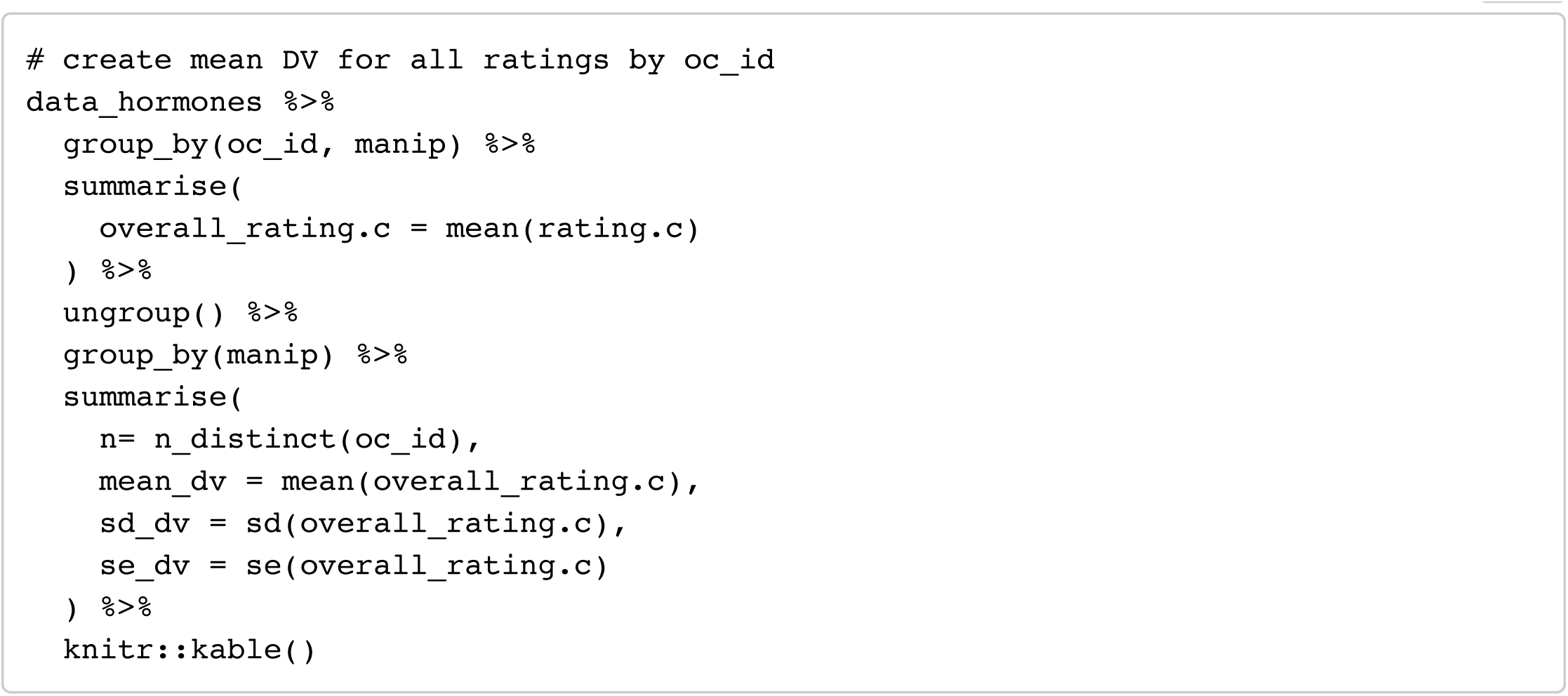

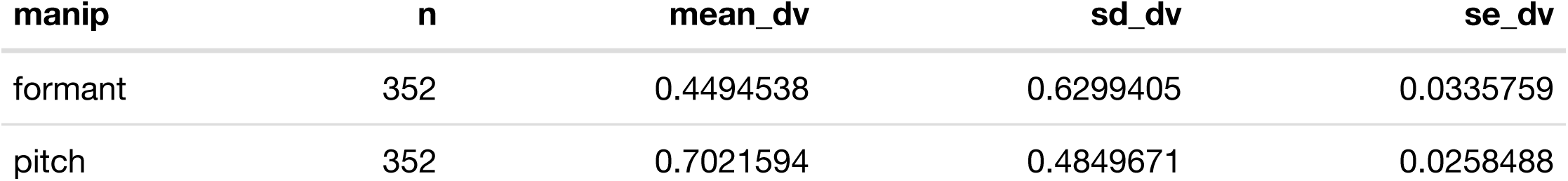

### Hormones

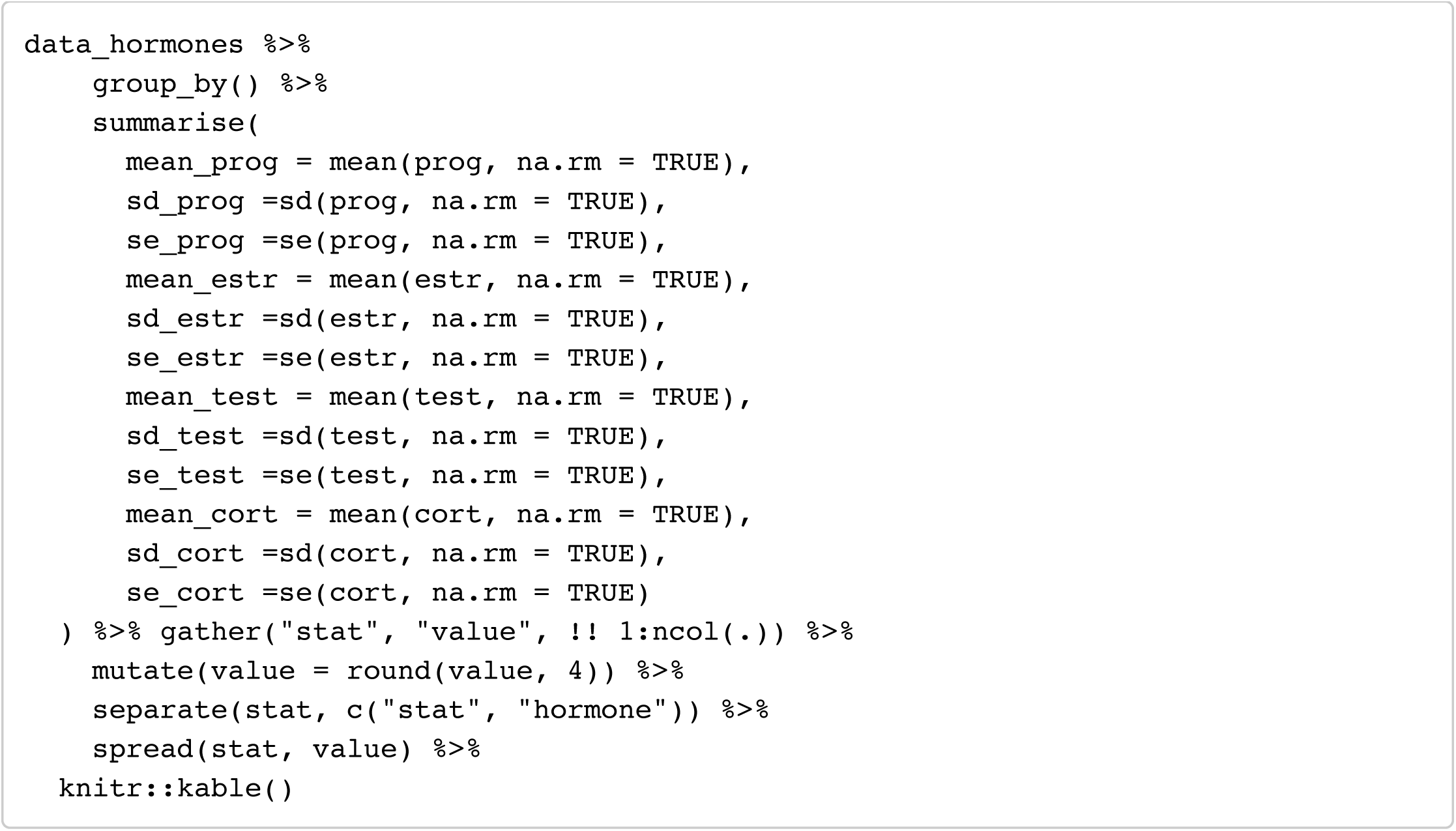

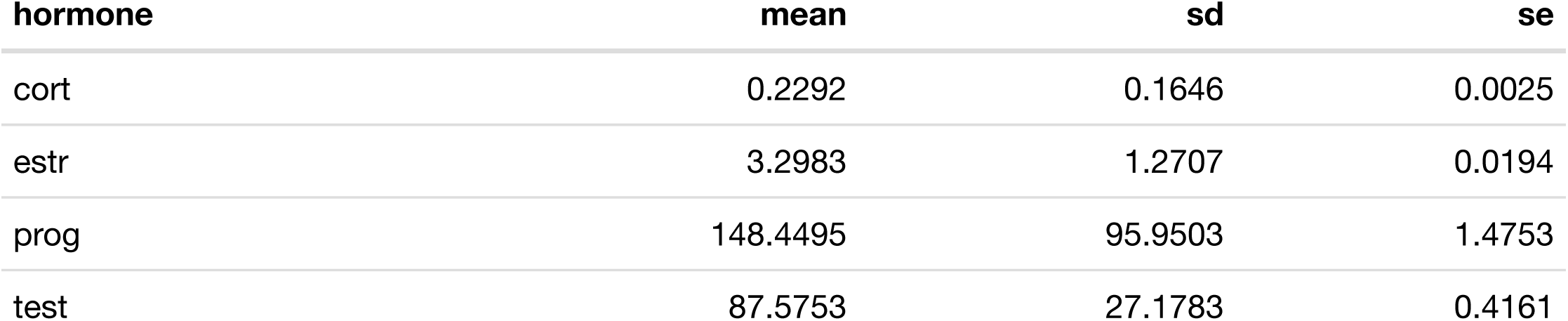

## Analyses of all data

### Masculinity preference (formants)

#### Model 1: formants ∼ E + P + E × P

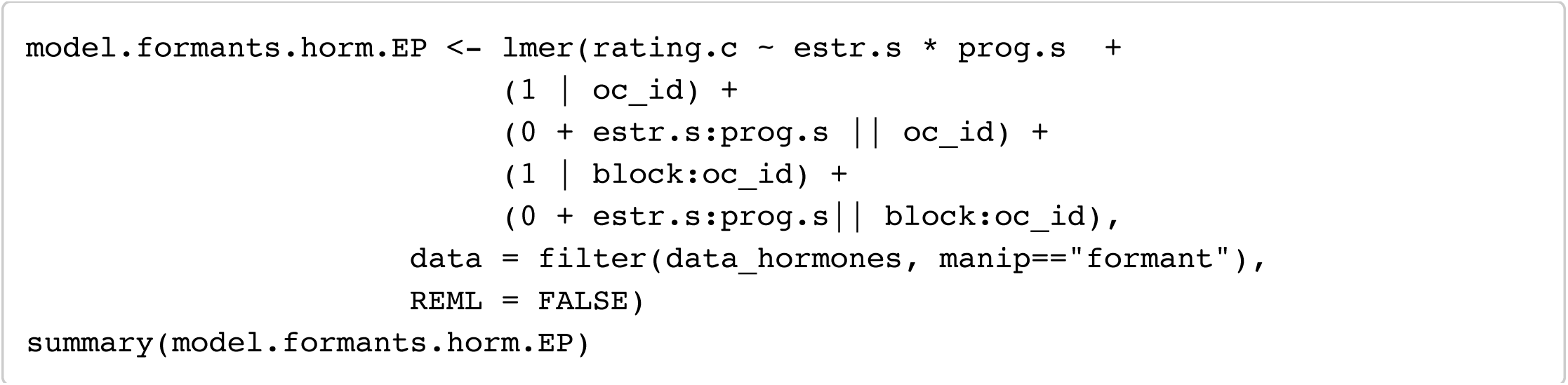

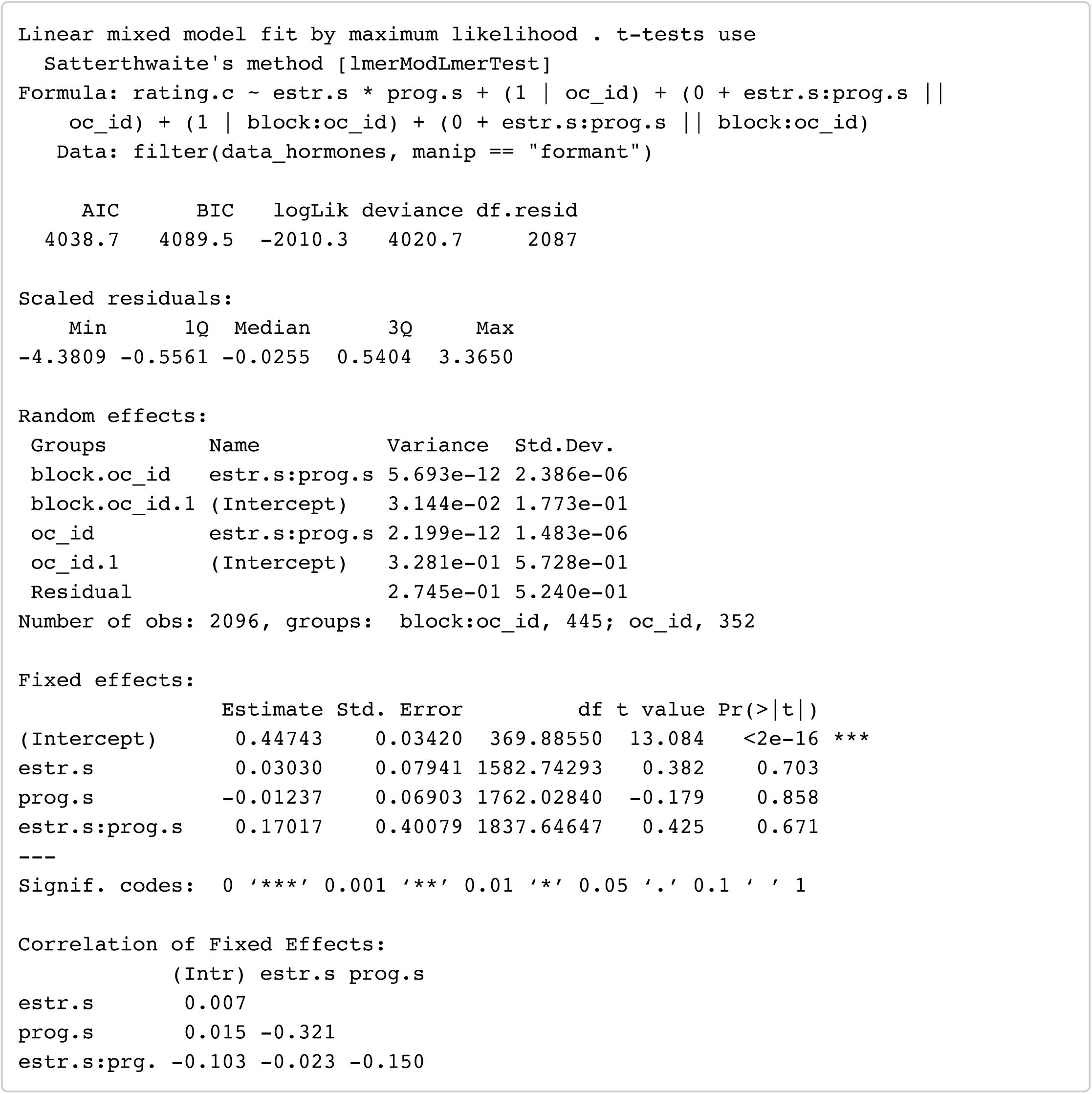

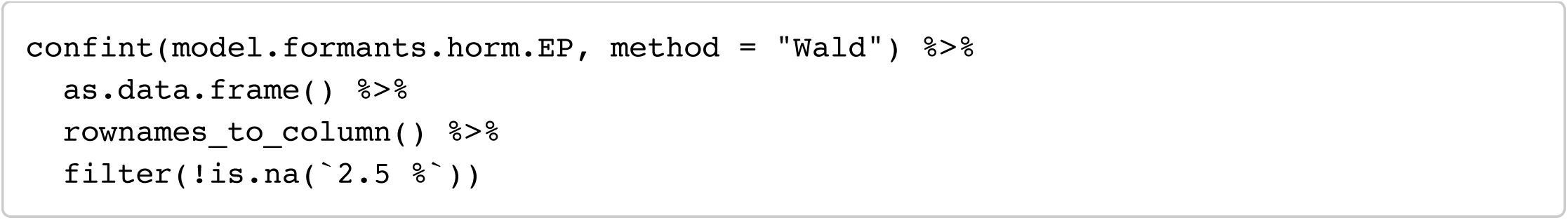

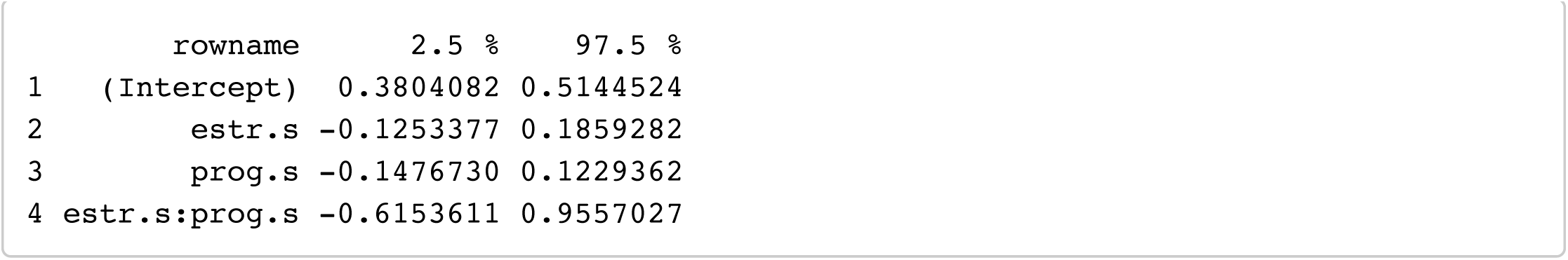

#### Model 2: formants ∼ E + P + EP_ratio

Hide

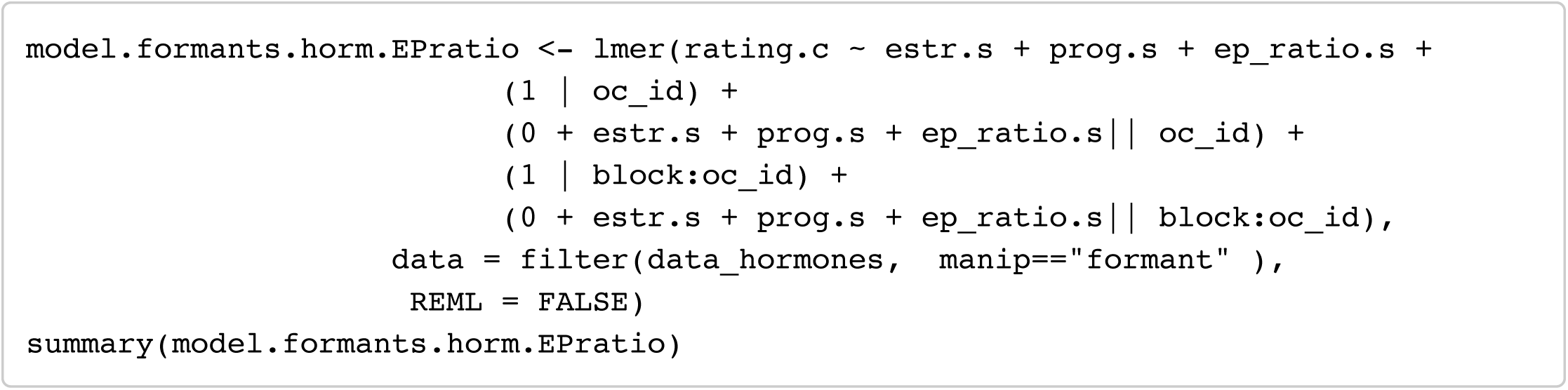

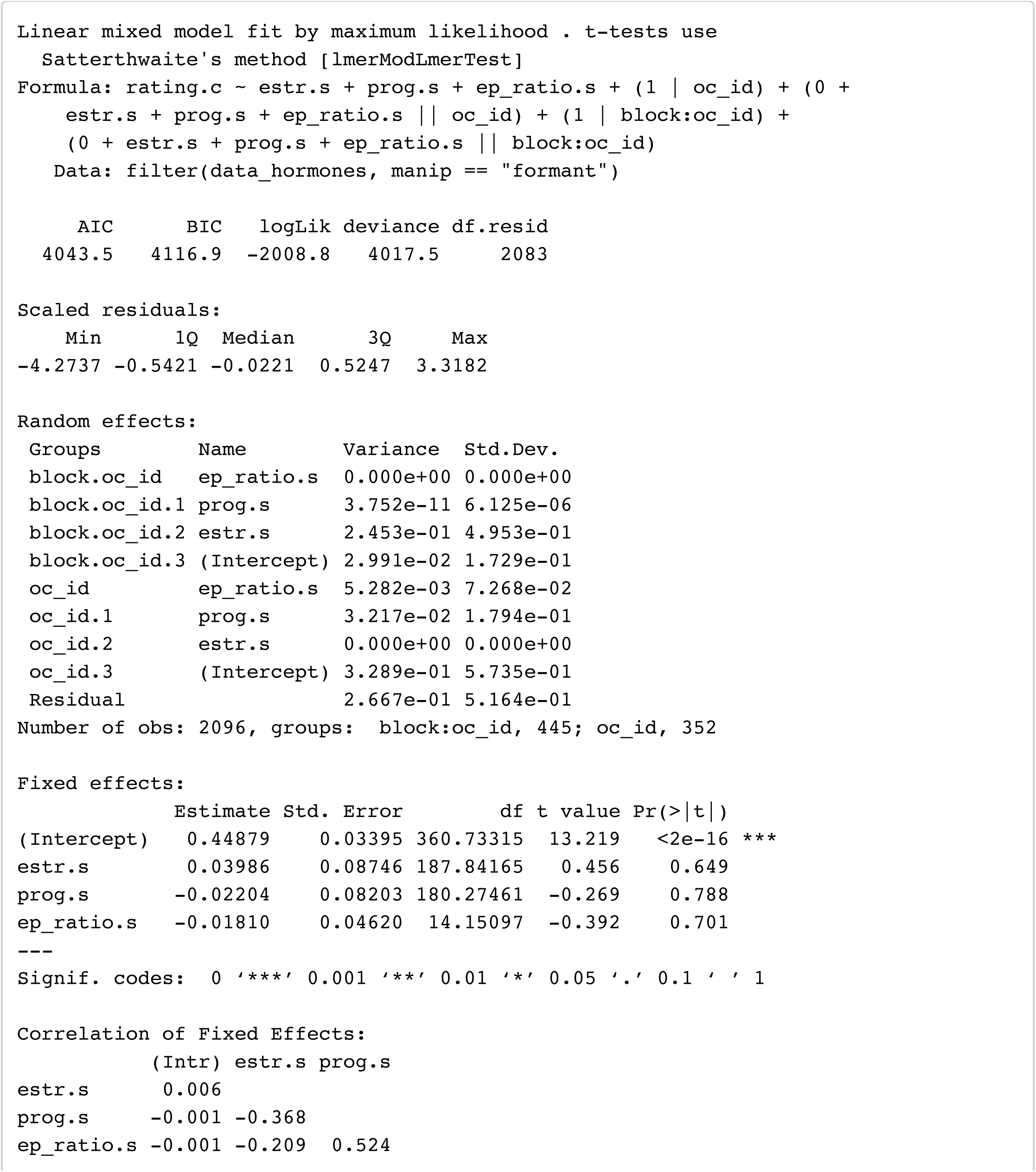

Hide

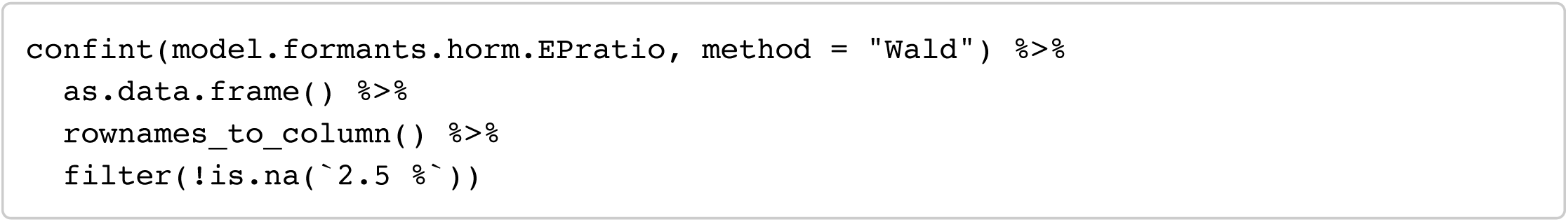

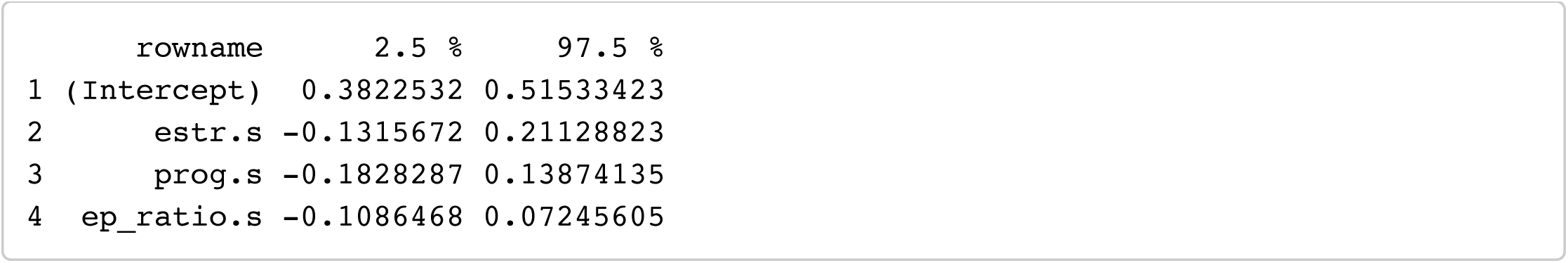

#### Model 3: formants ∼ T + C

Hide

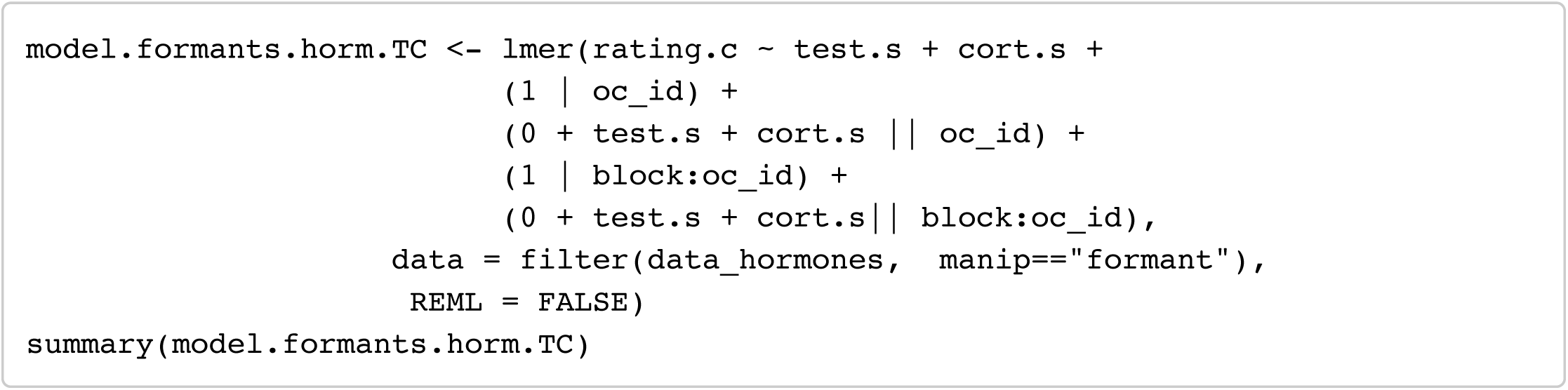

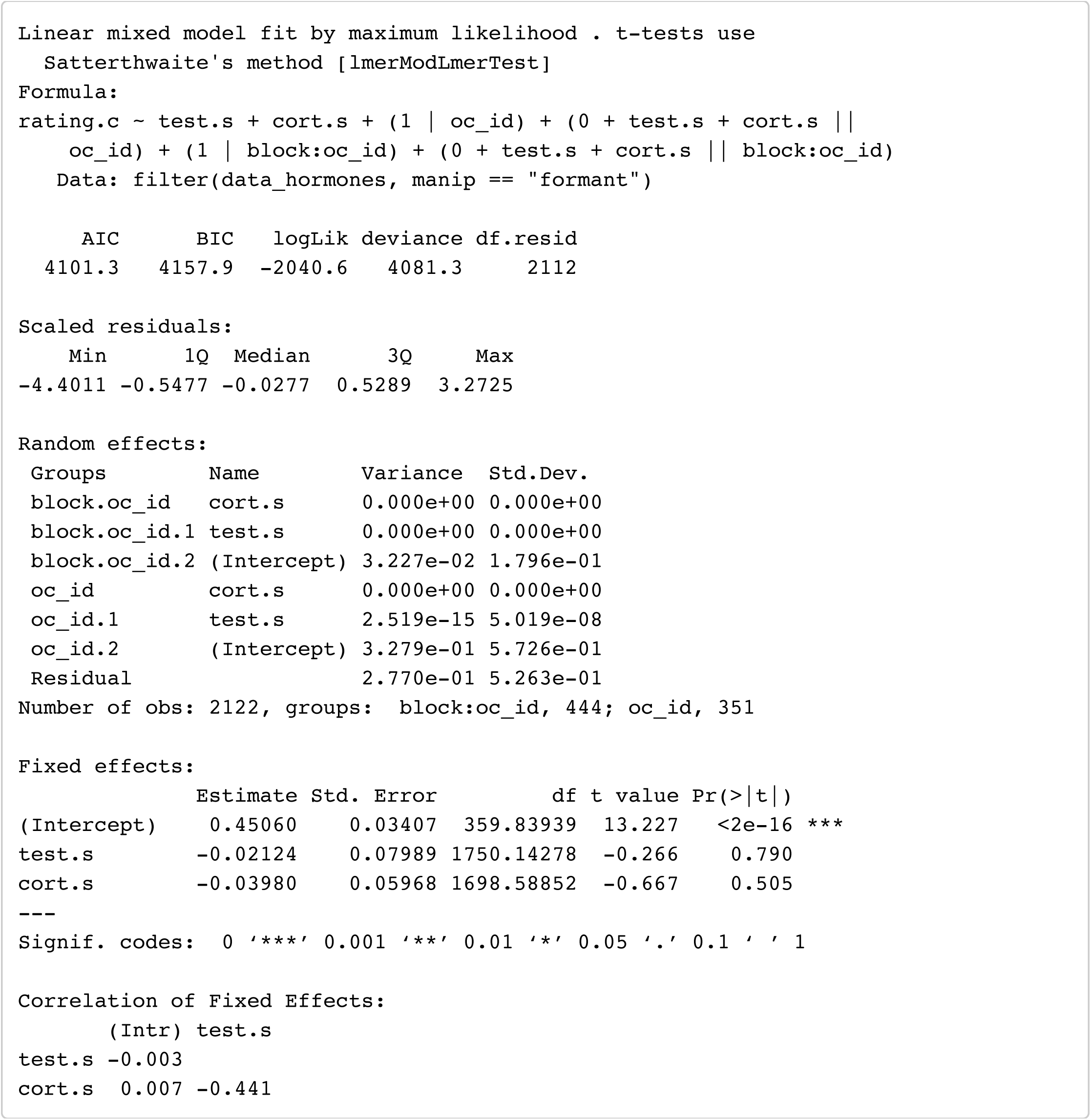

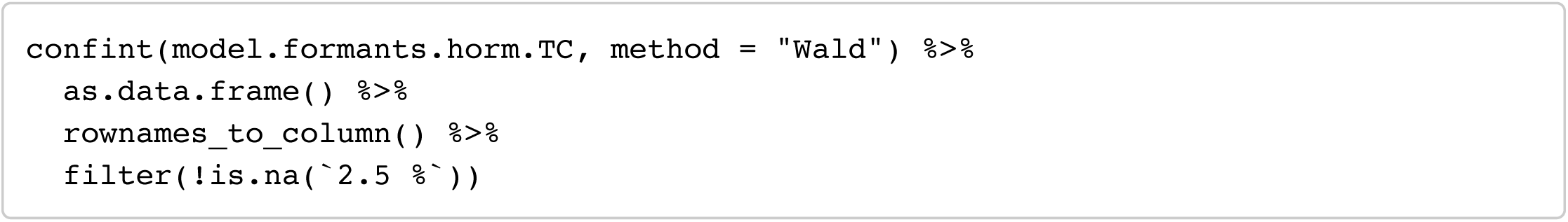

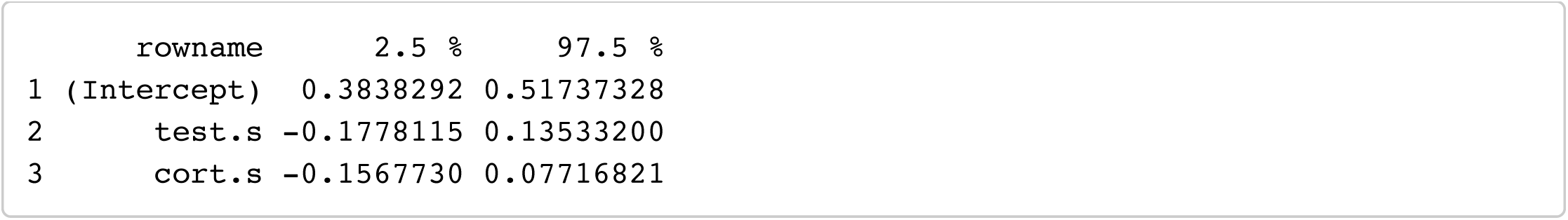

### Masculinity preference (pitch)

#### Model 1: Pitch ∼ E + P + E × P

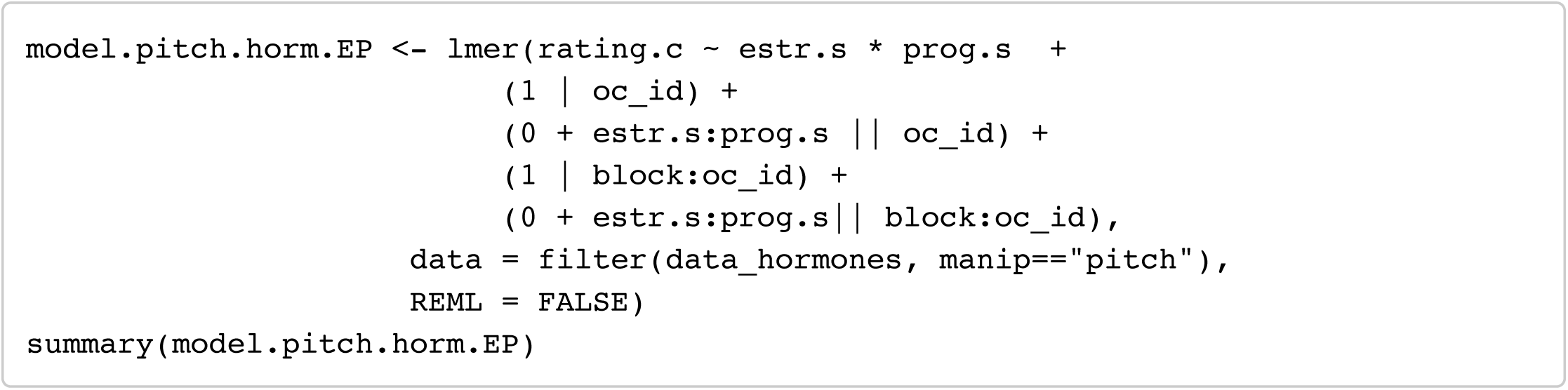

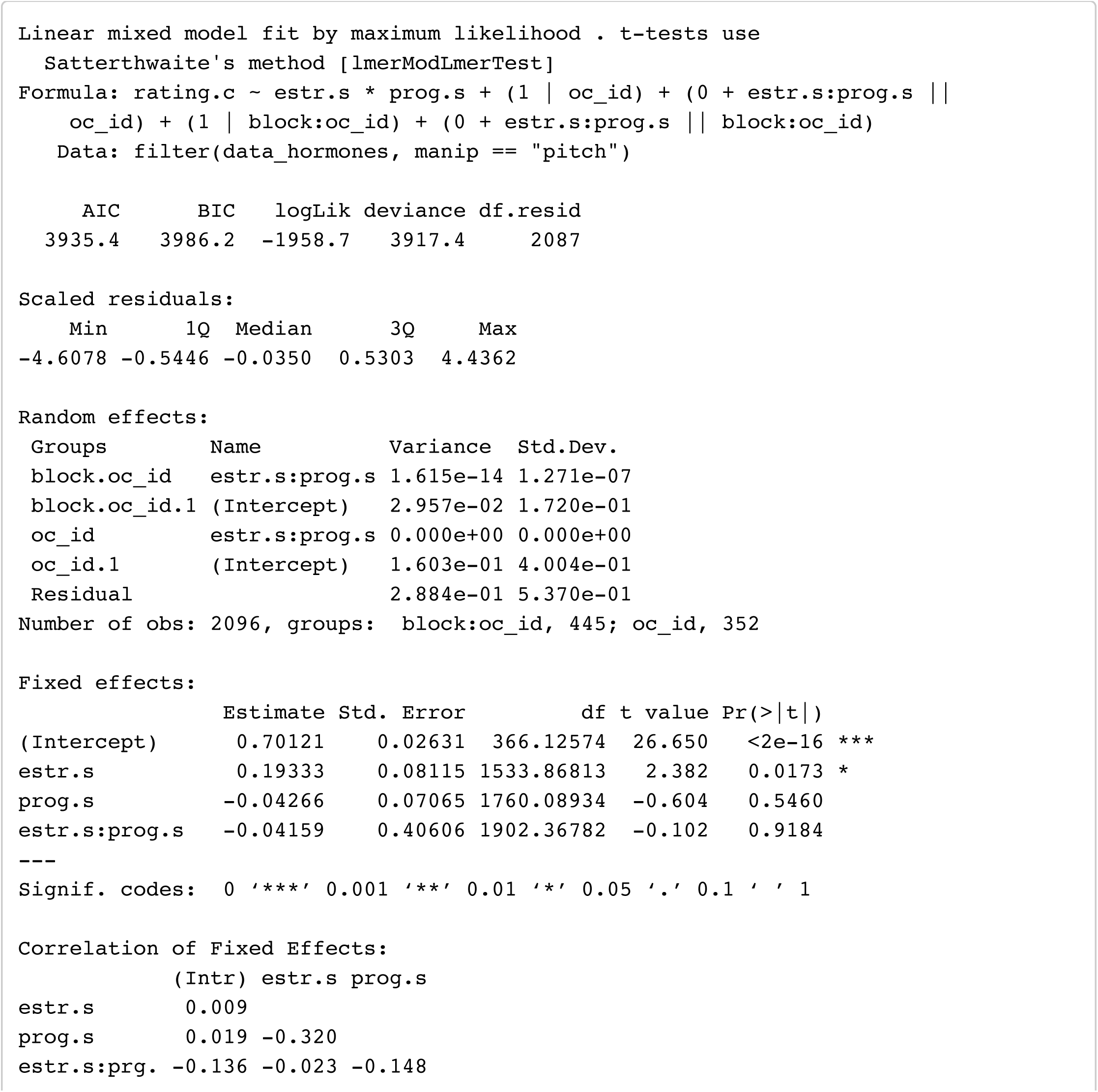

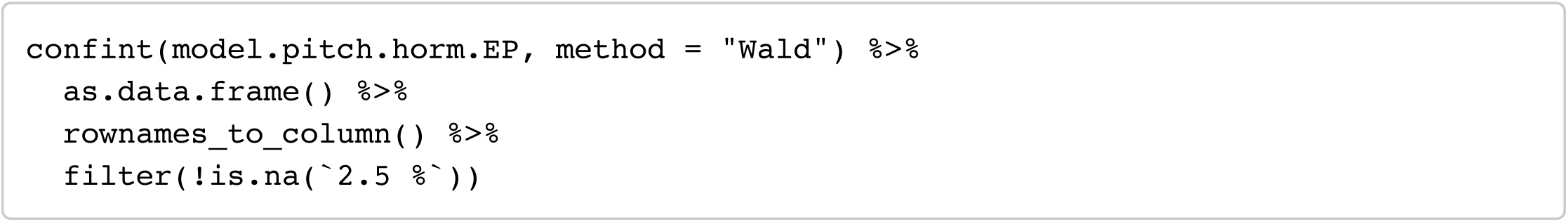

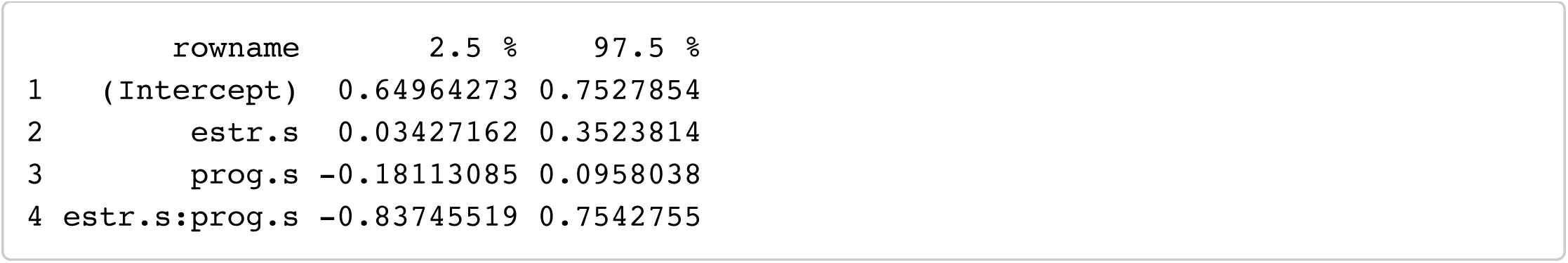

#### Model 2: Pitch ∼ E + P + EP_ratio

Hide

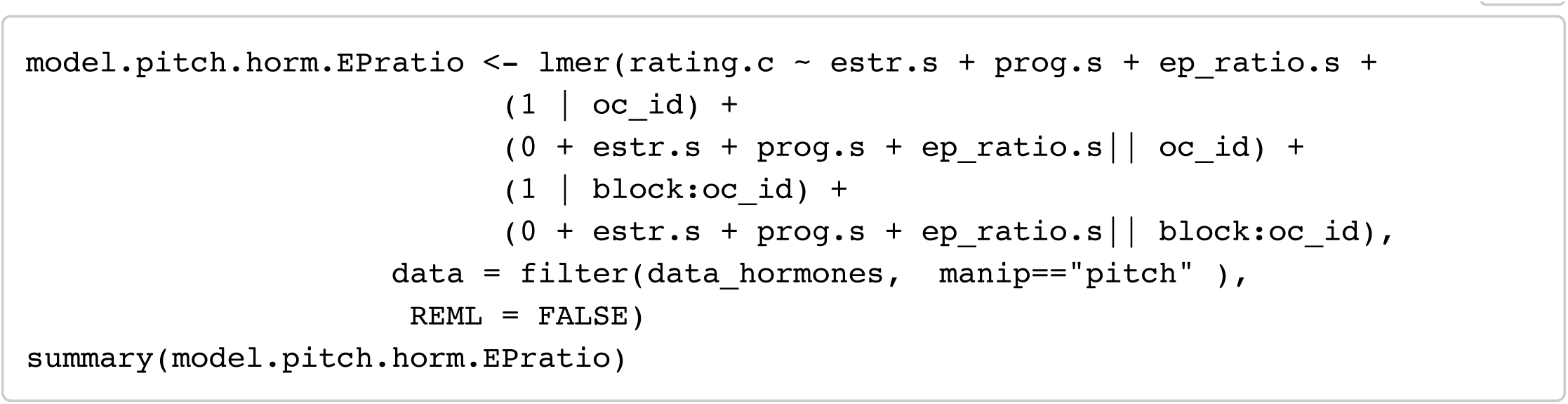

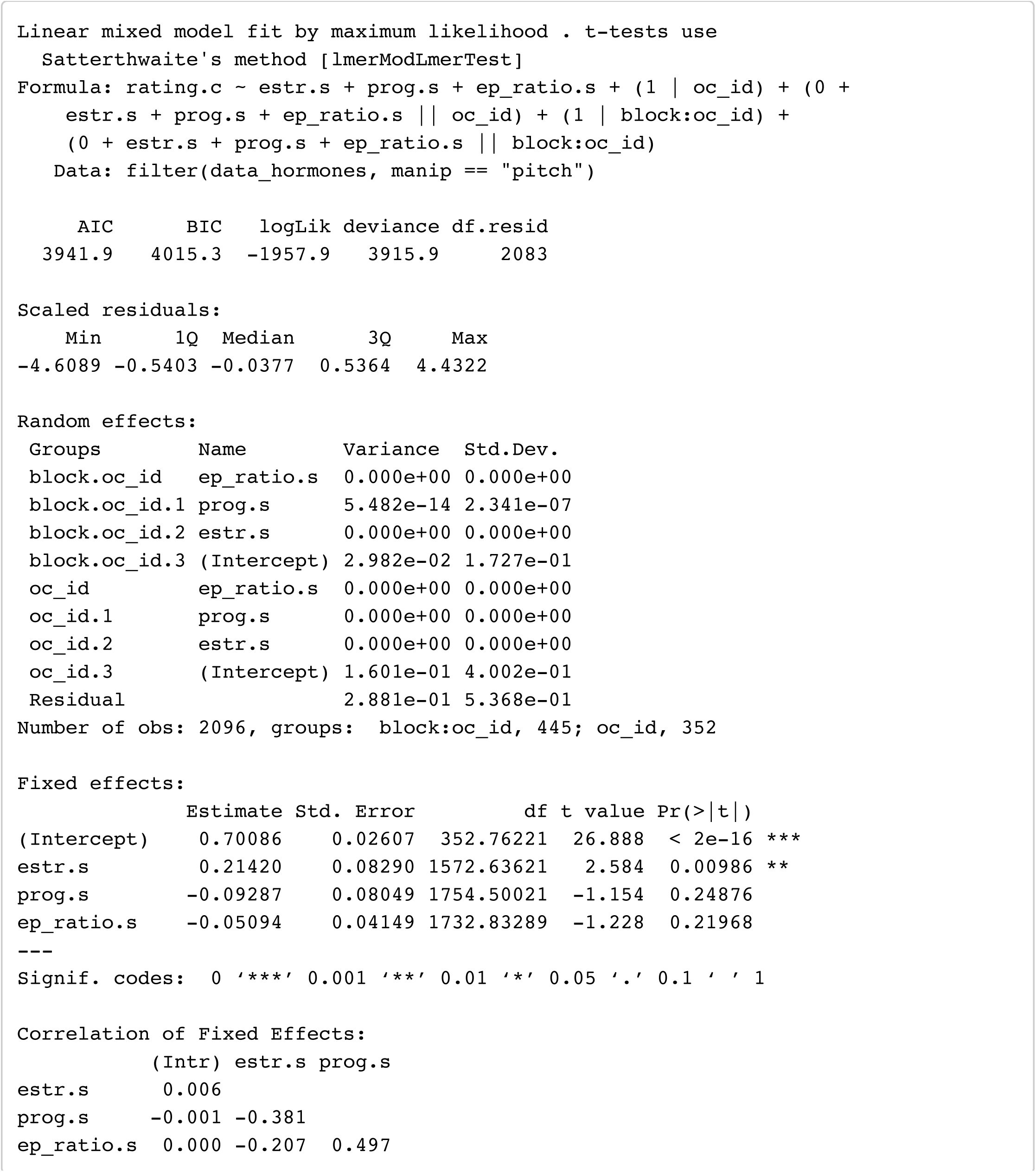

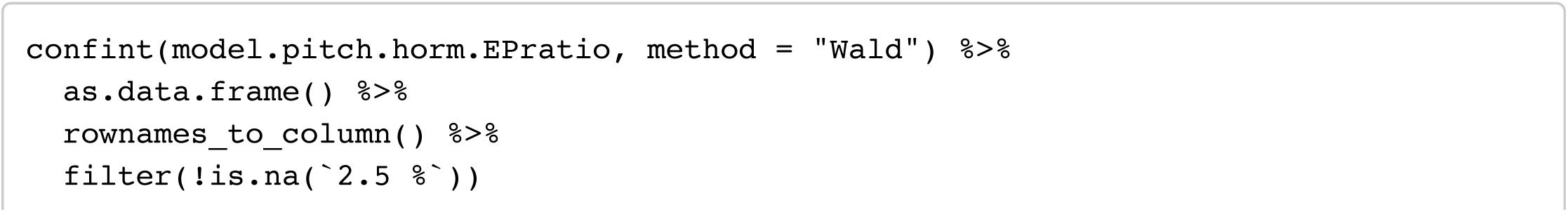

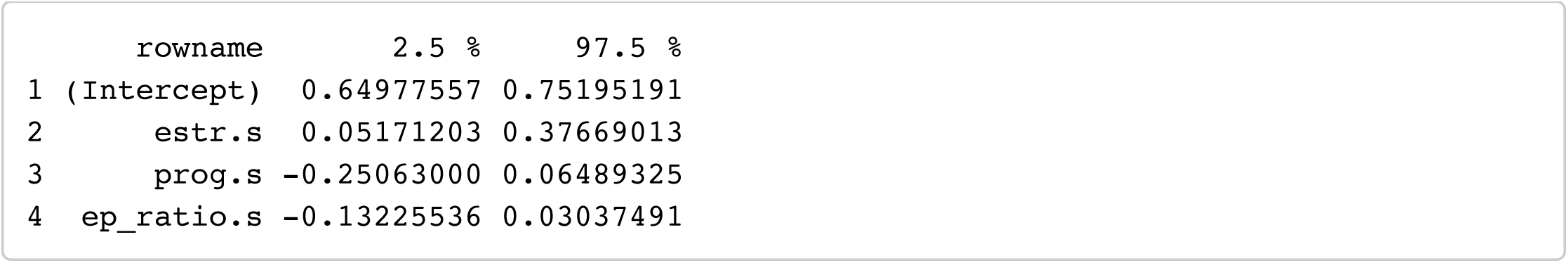

#### Model 3: Pitch ∼ T + C

Hide

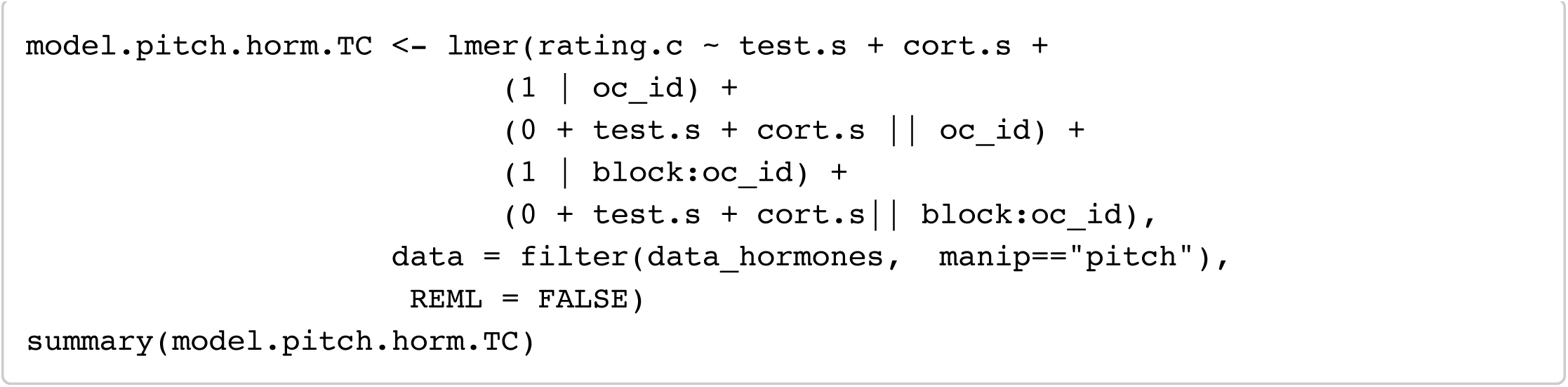

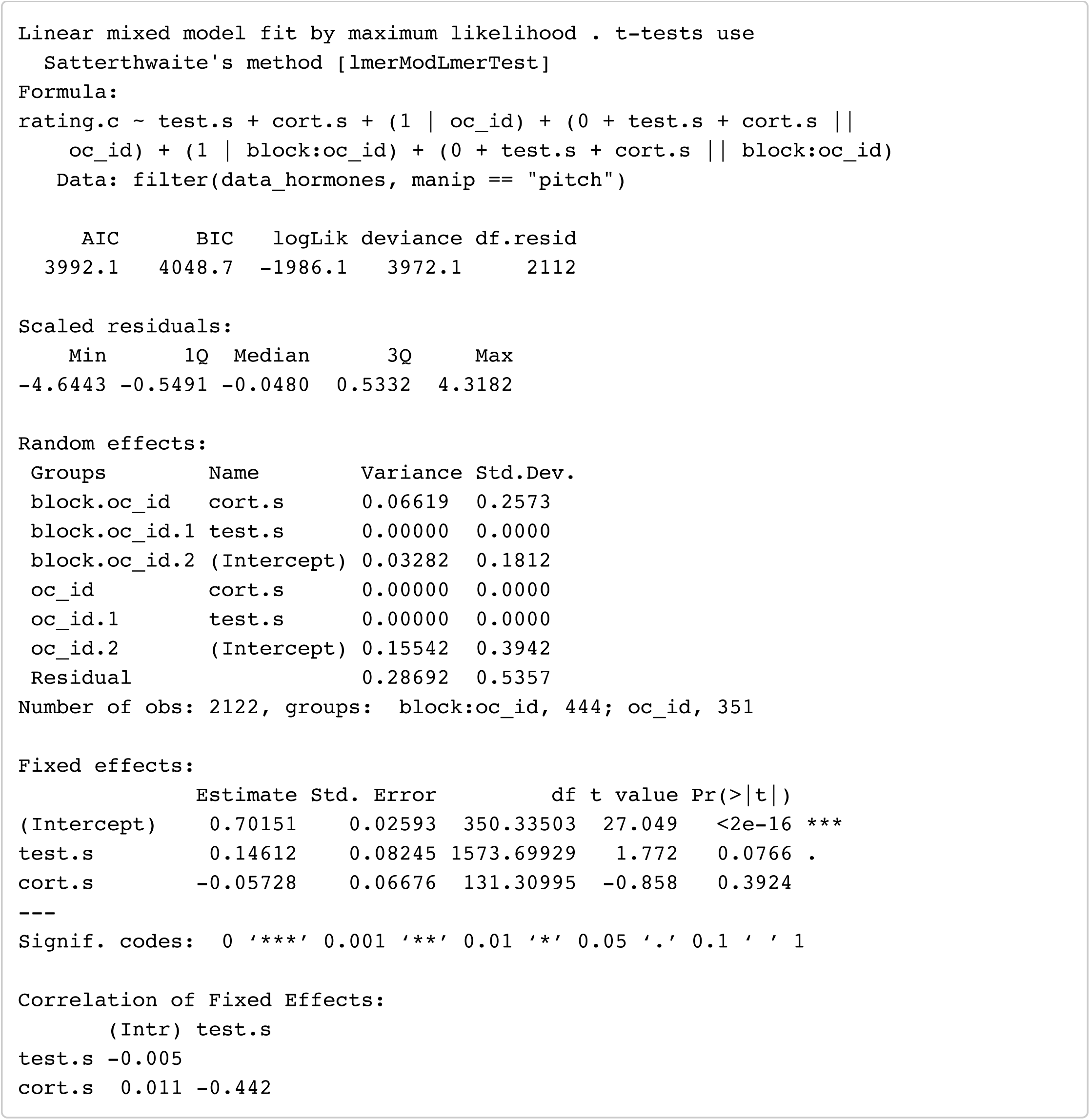

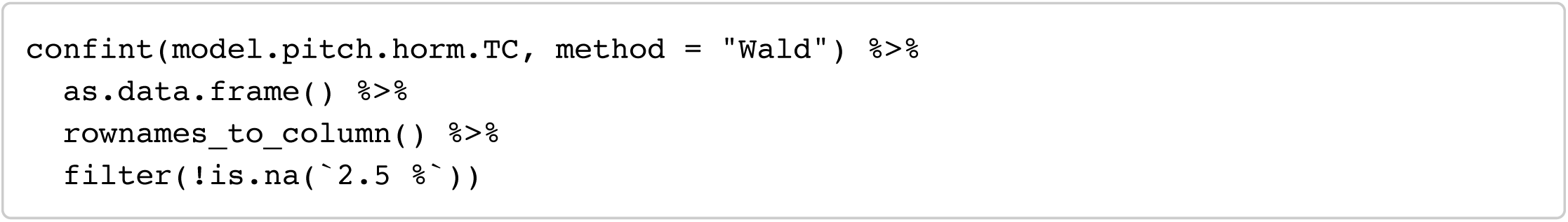

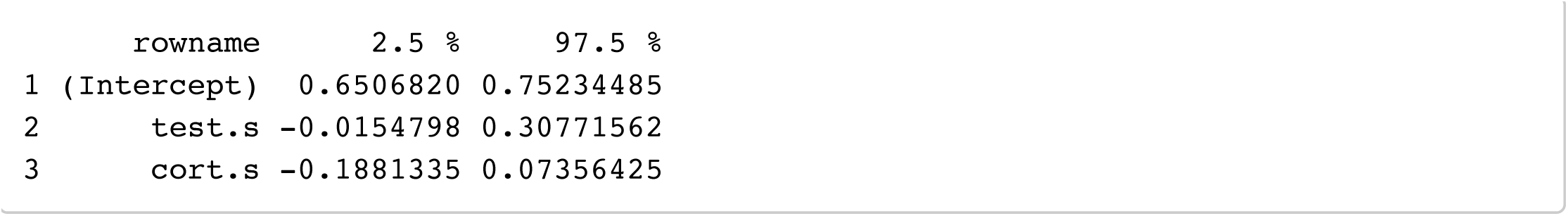

## Analyses excluding responses from Pisanski et al (2014)

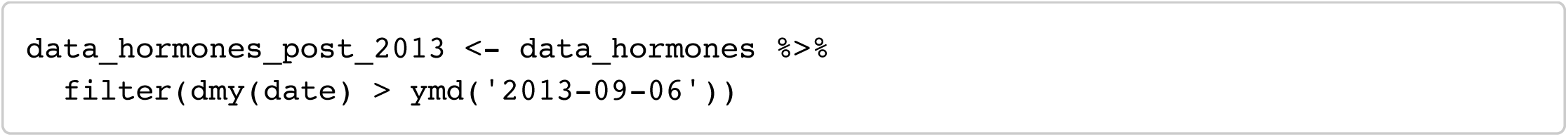

### Masculinity preference (formants)

Hide

#### Model 1: formants ∼ E + P + E x P

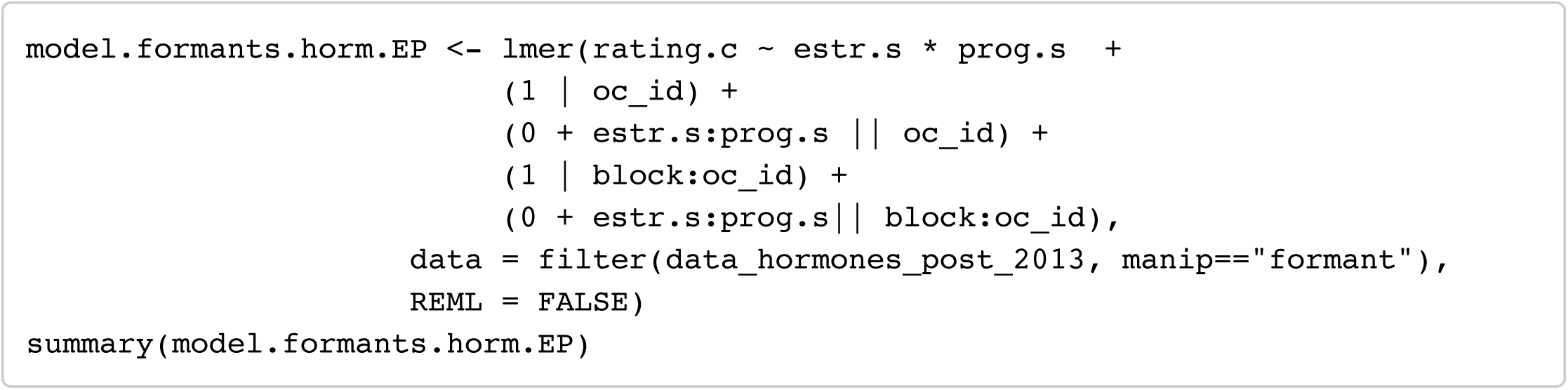

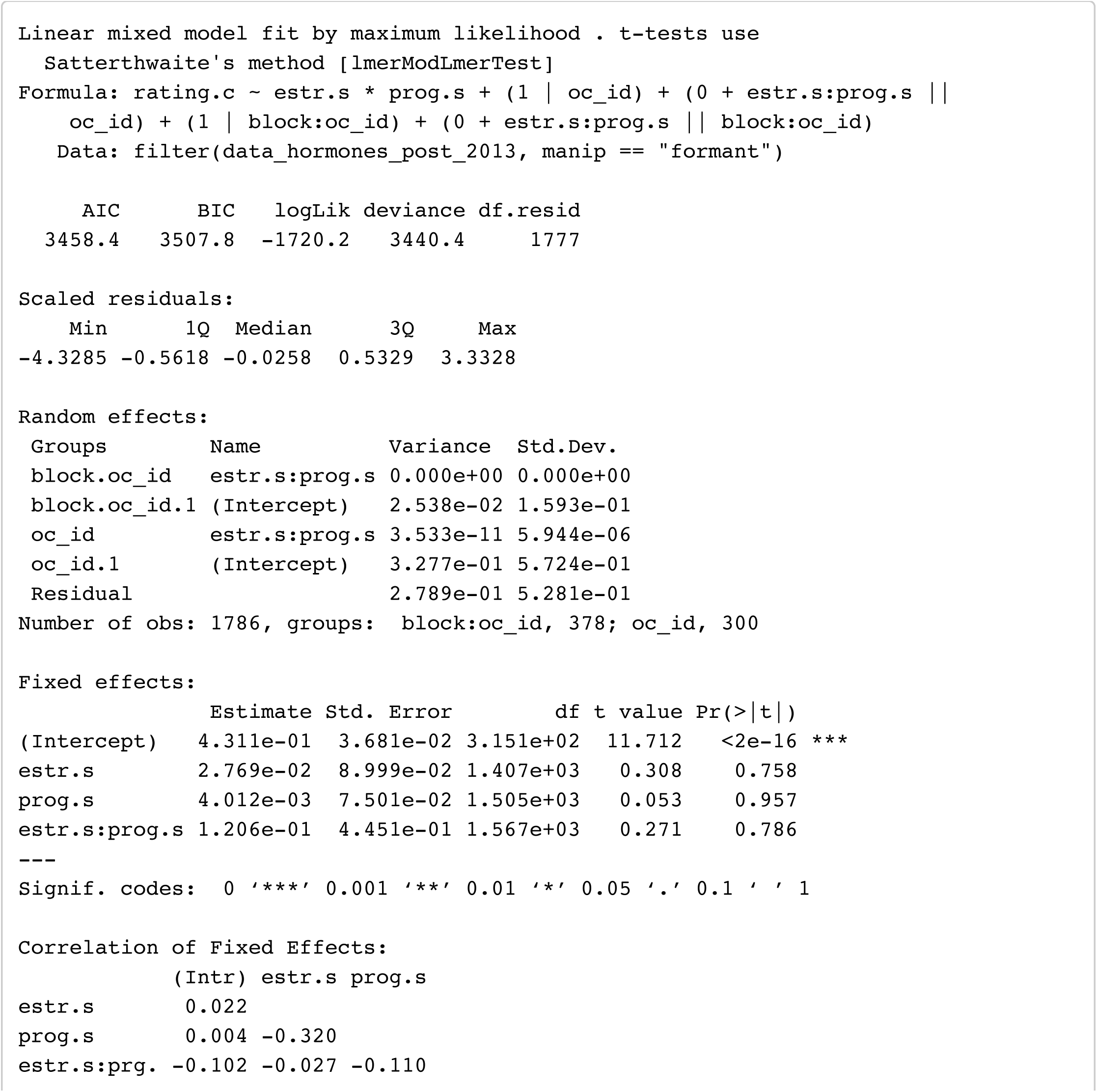

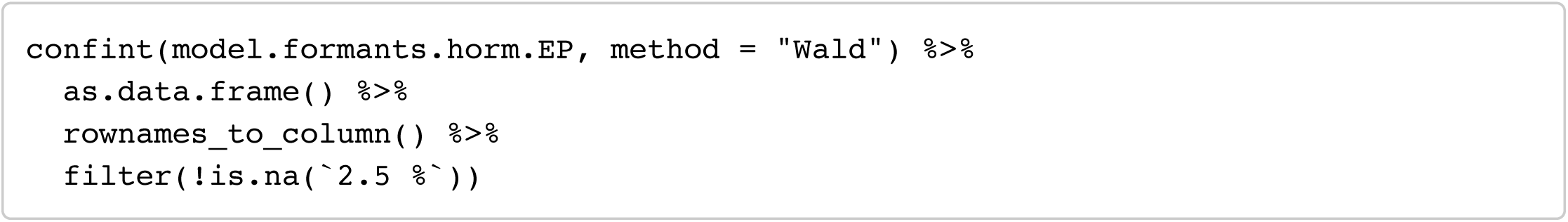

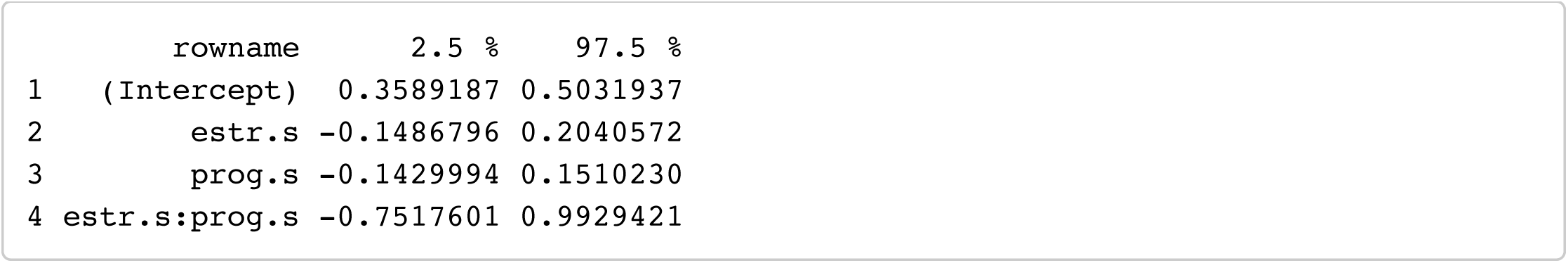

#### Model 2: formants ∼ E + P + EP_ratio

Hide

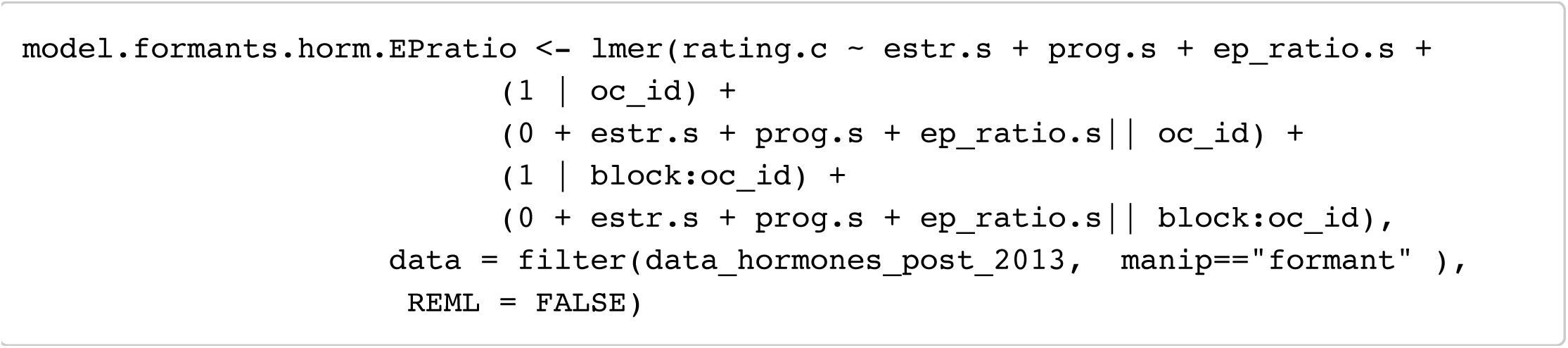

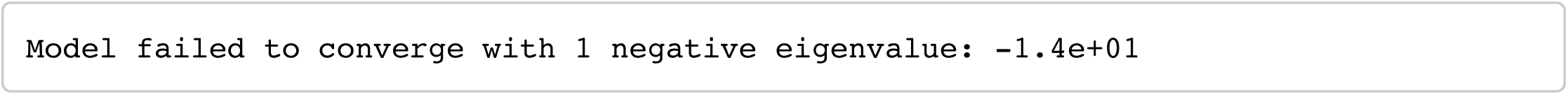

Hide

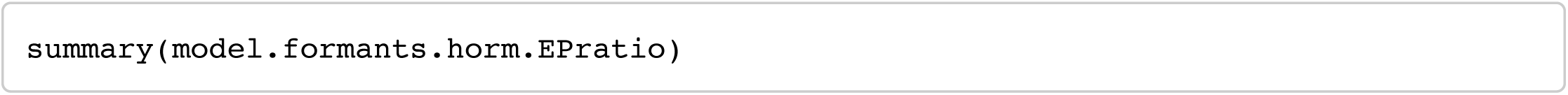

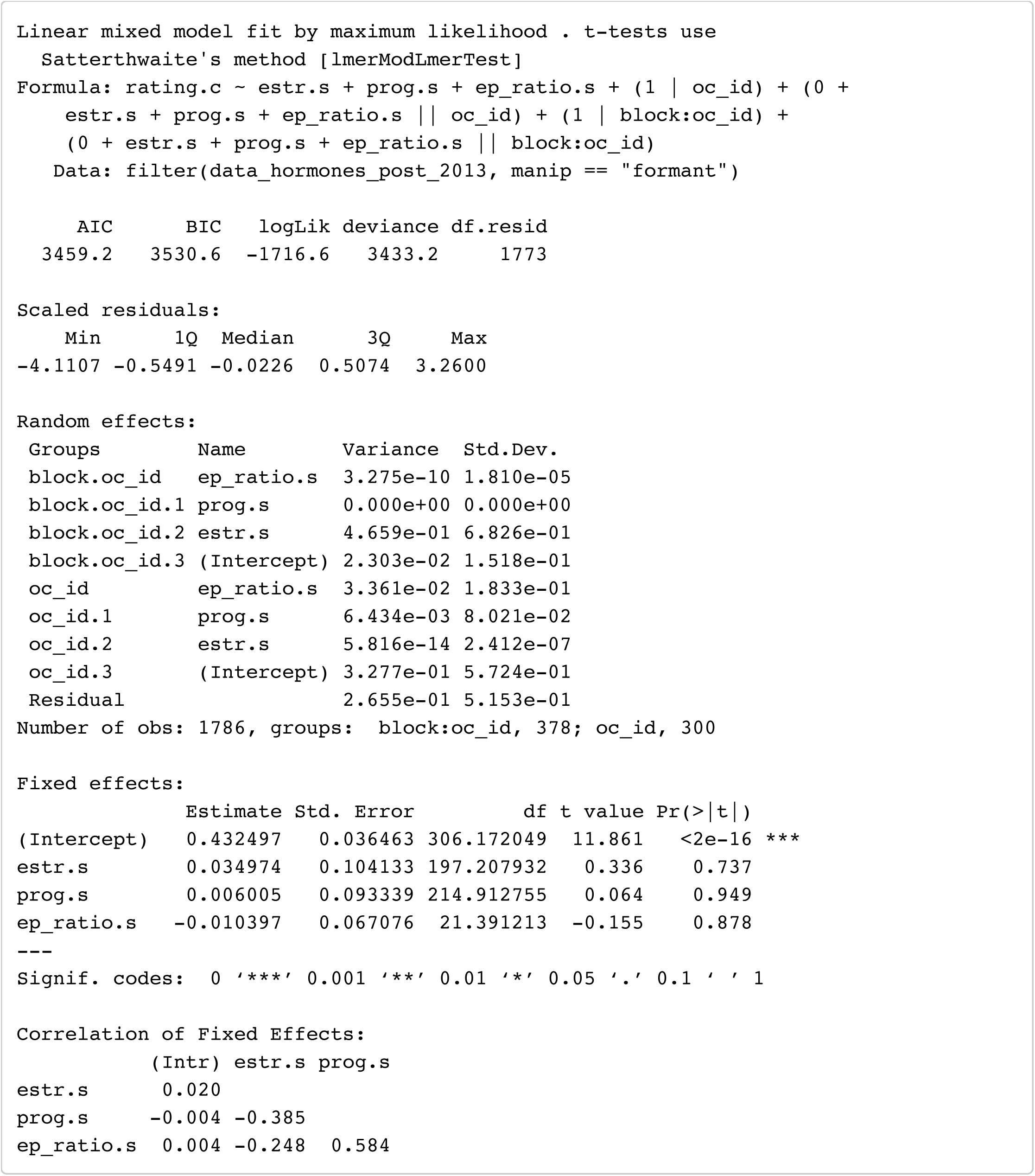

Hide

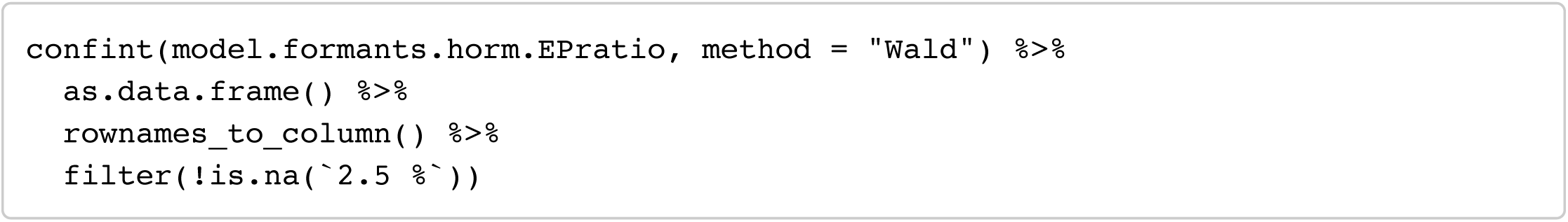

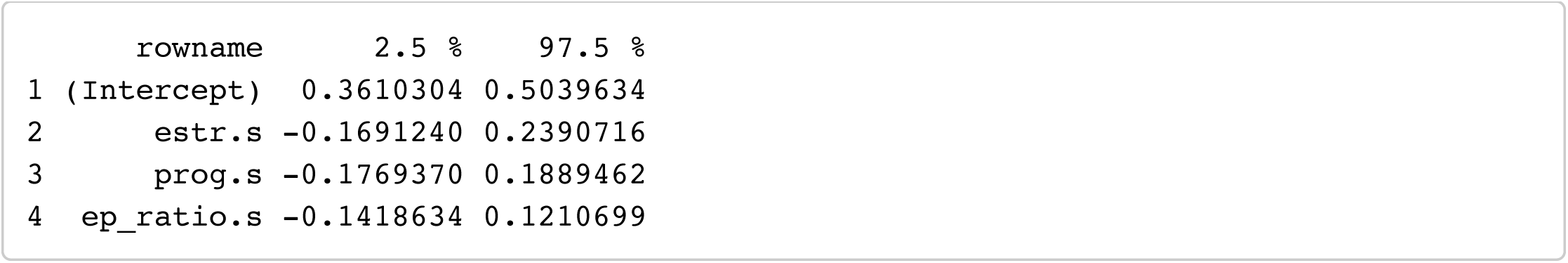

#### Model 3: formants ∼ T + C

Hide

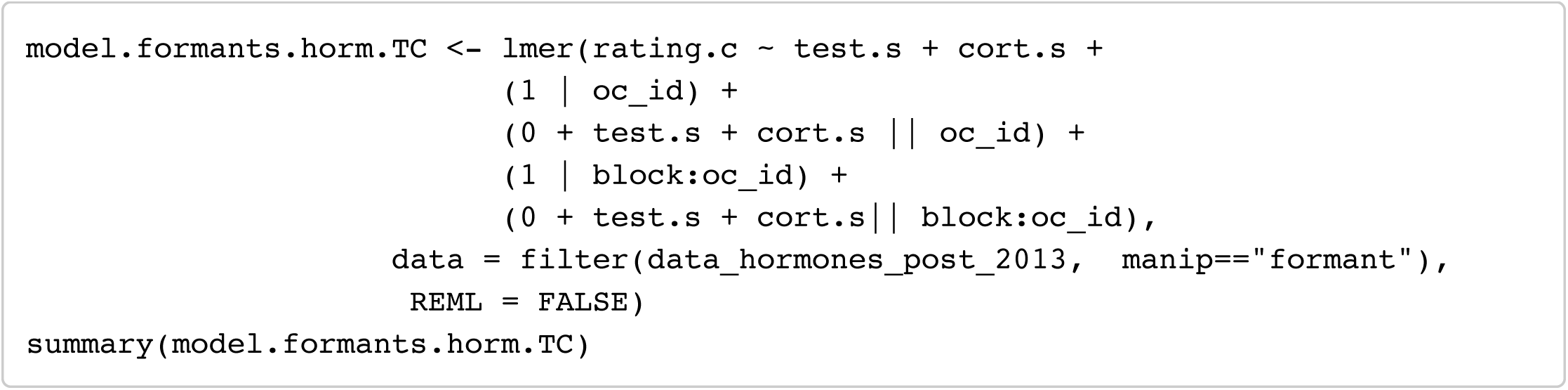

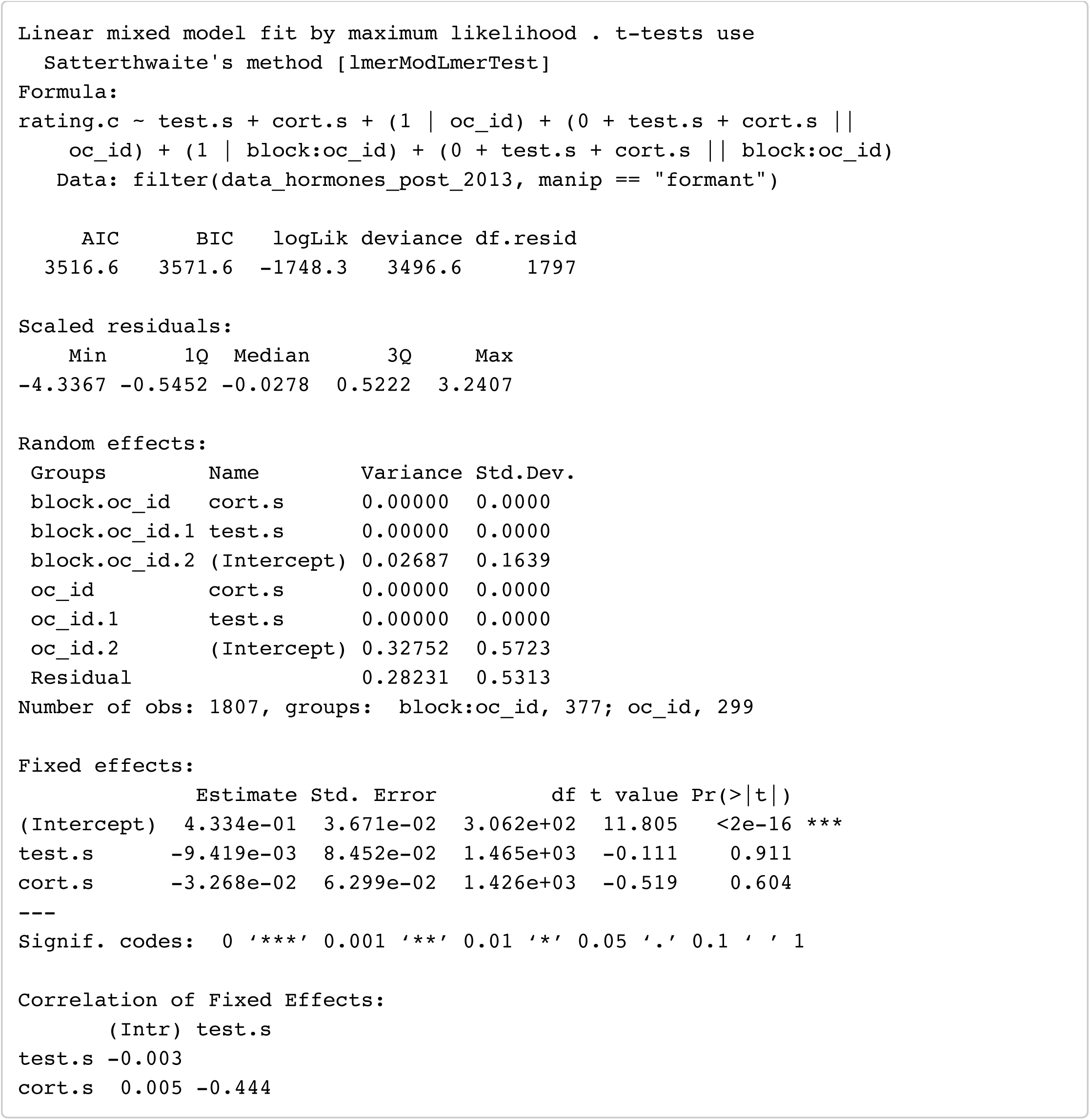

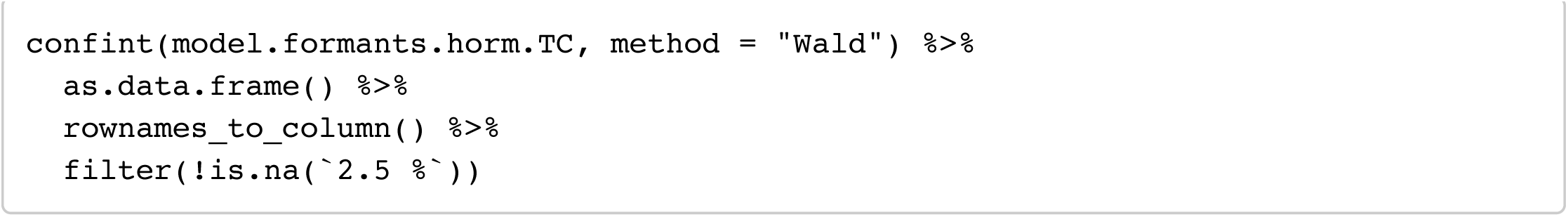

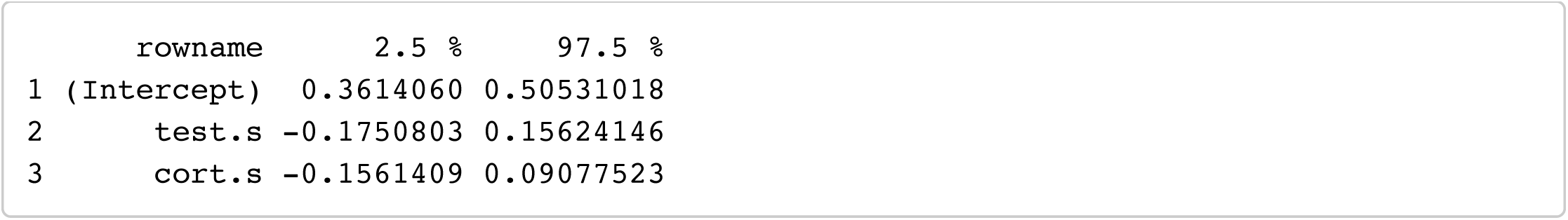

### Masculinity preference (pitch)

#### Model 1: Pitch ∼ E + P + E x P

Hide

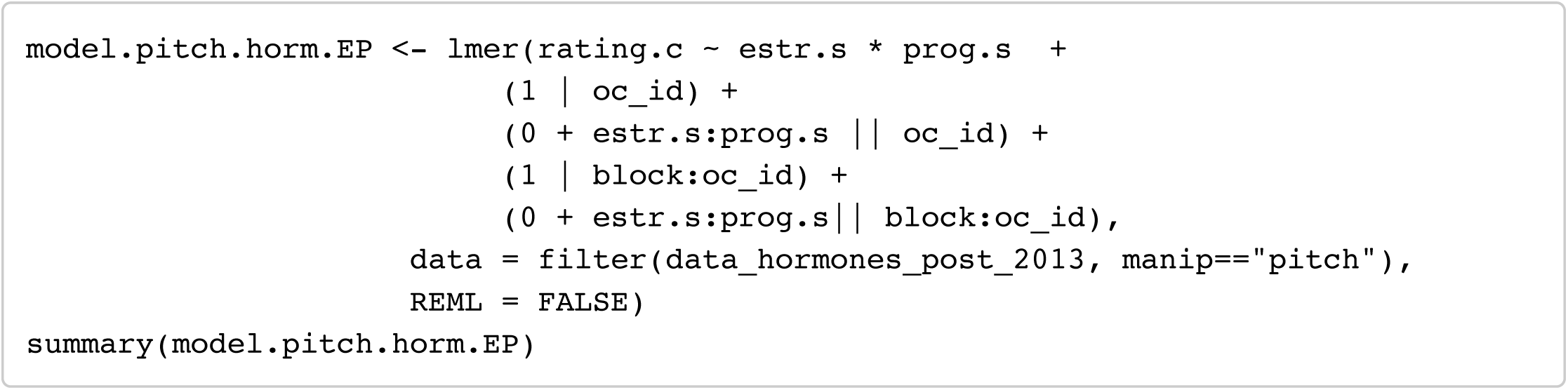

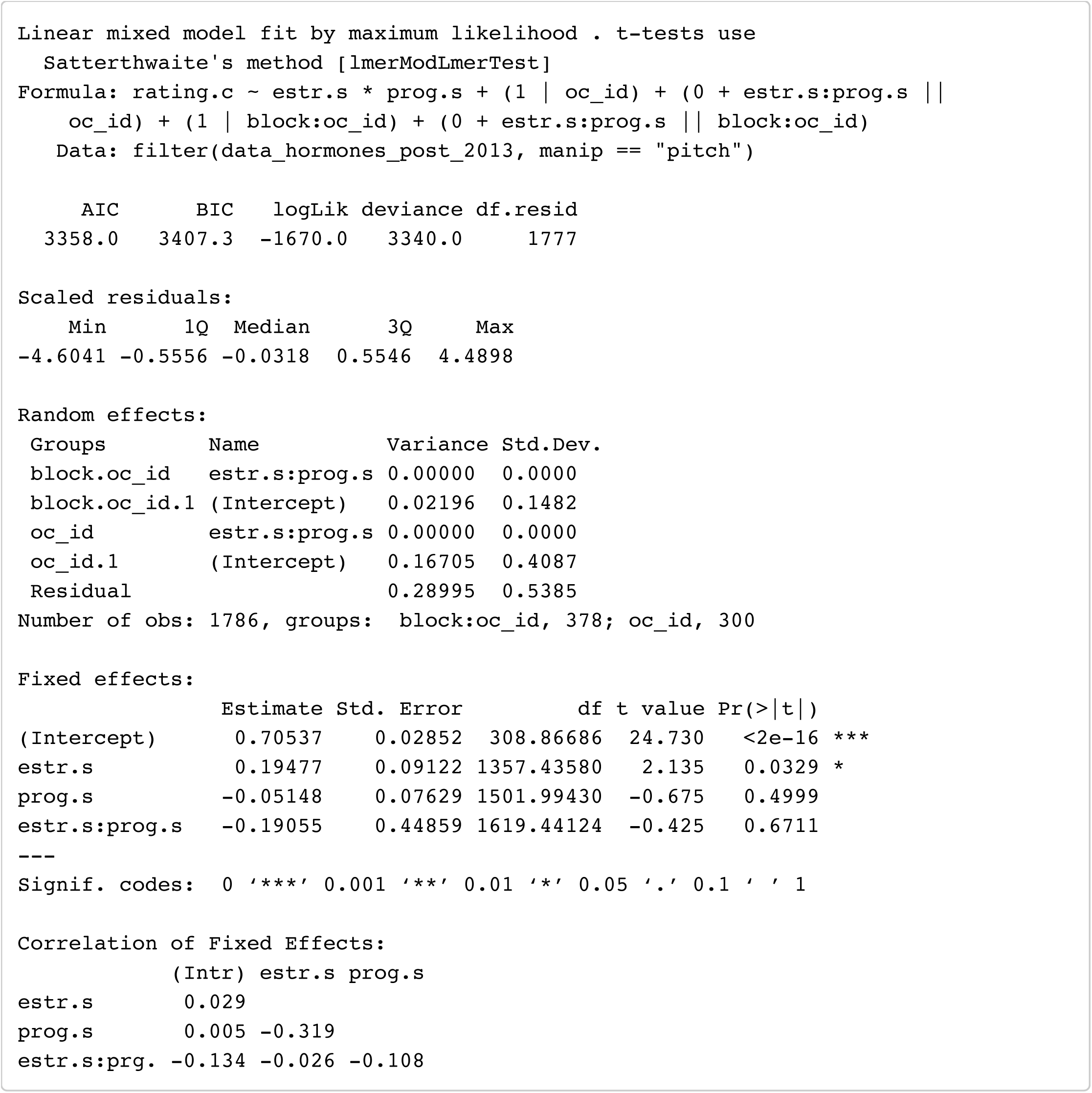

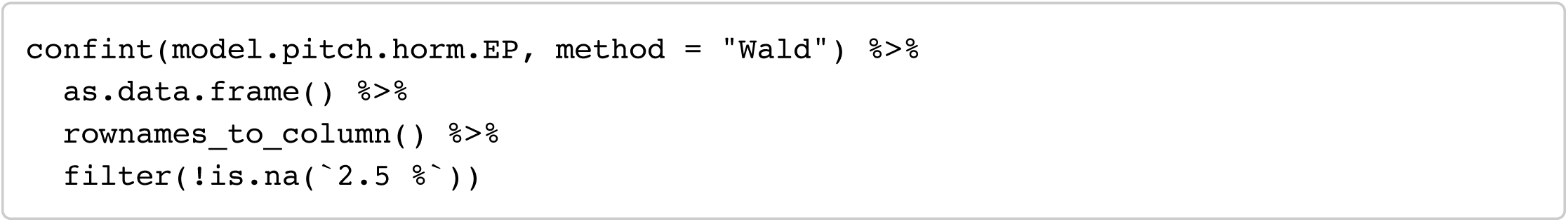

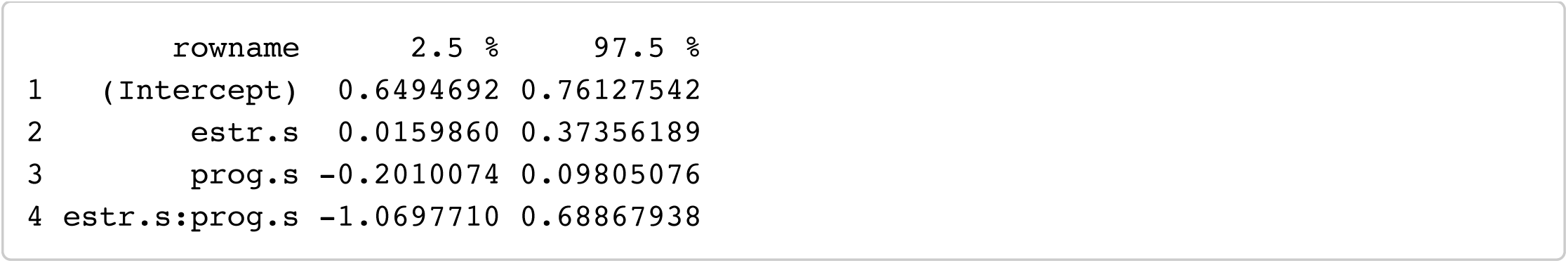

#### Model 2: Pitch ∼ E + P + EP_ratio

Hide

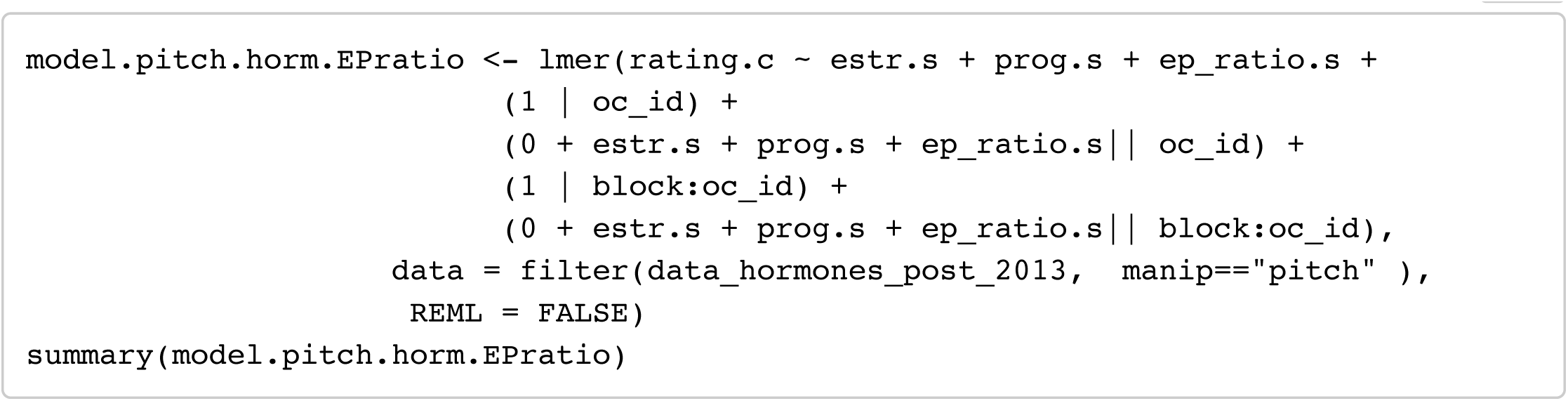

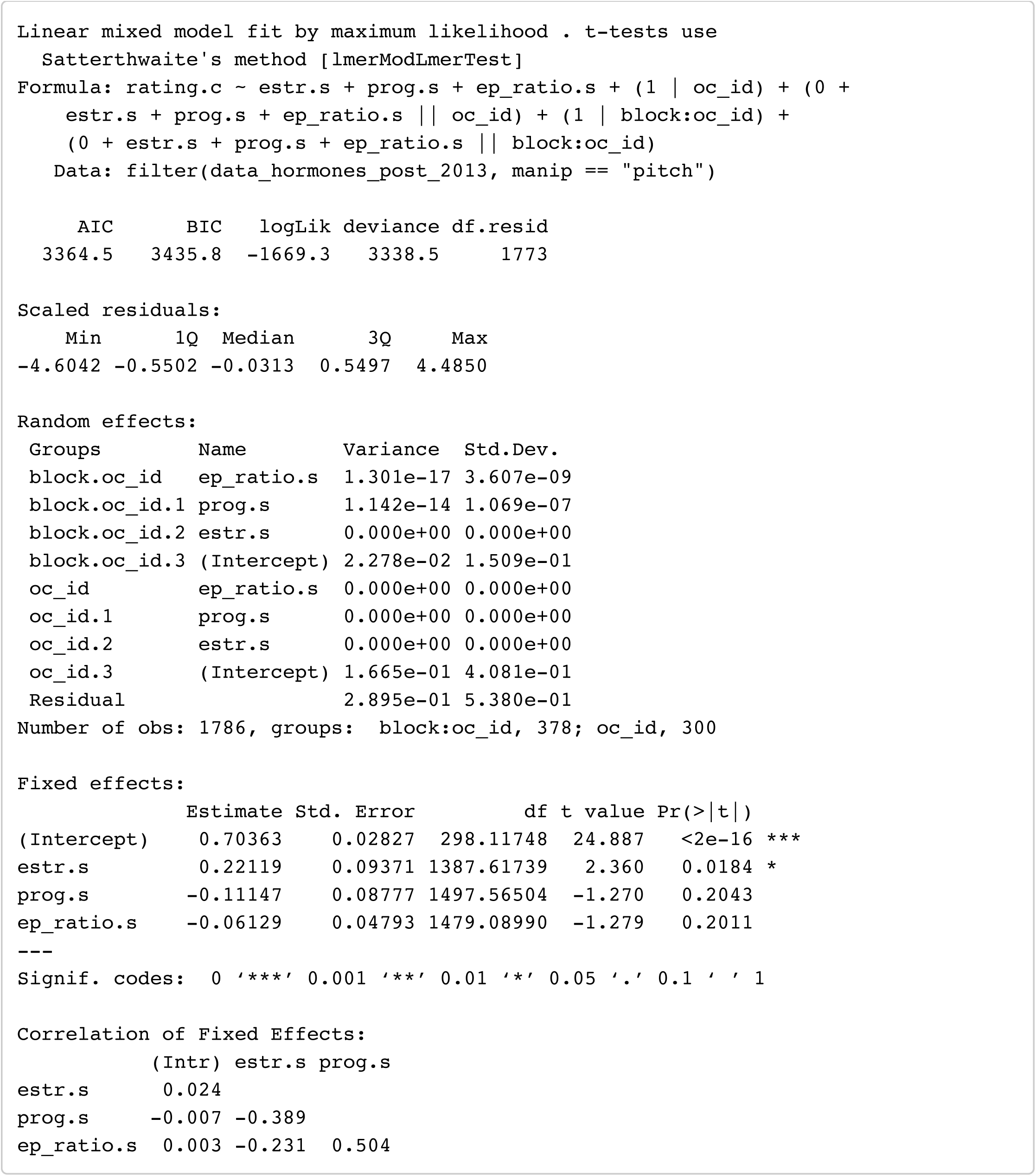

Hide

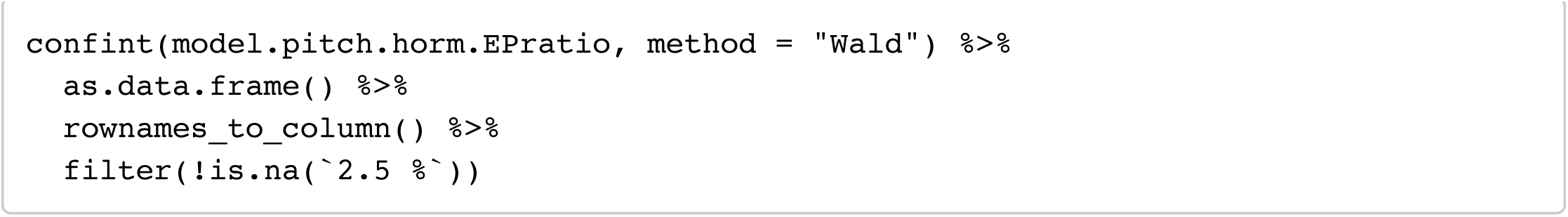

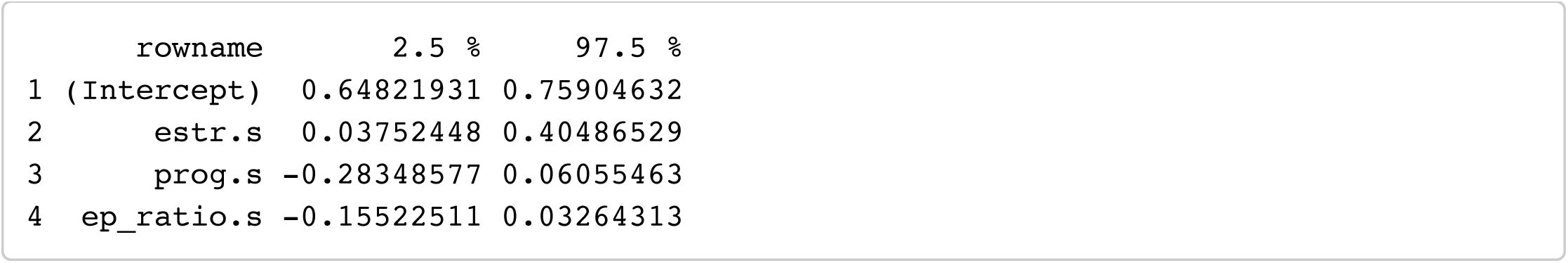

#### Model 3: Pitch ∼ T + C

Hide

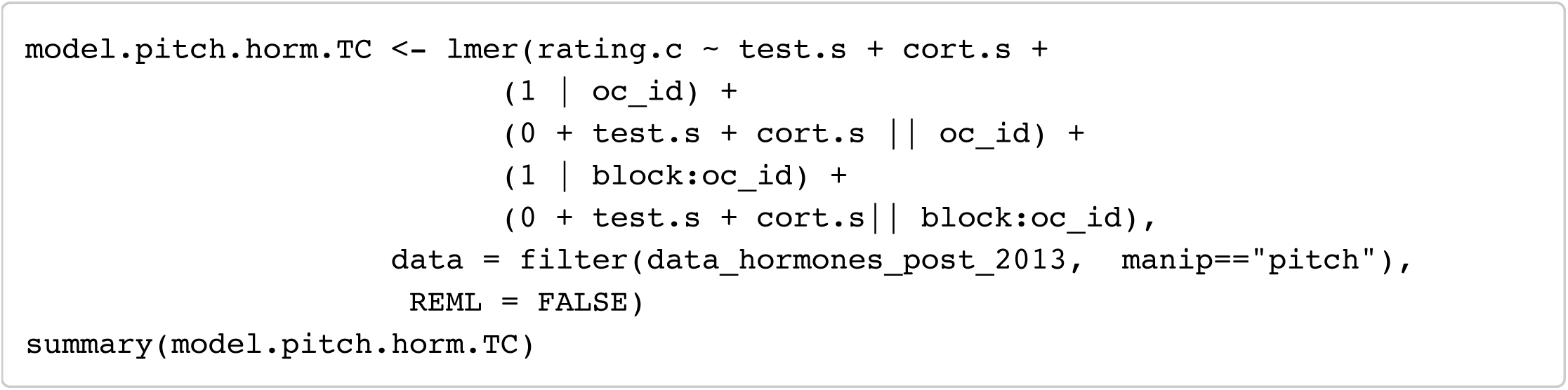

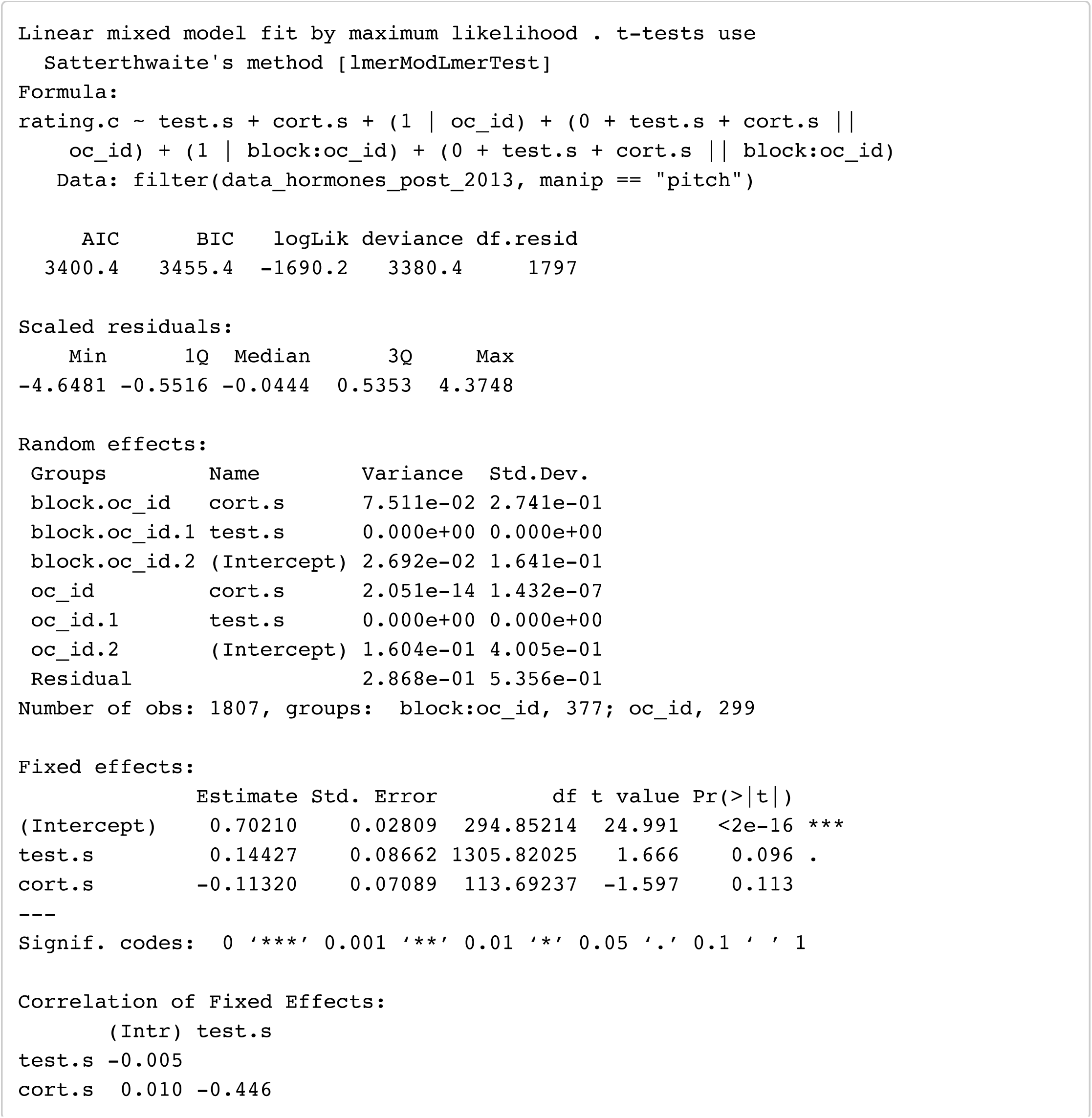

Hide

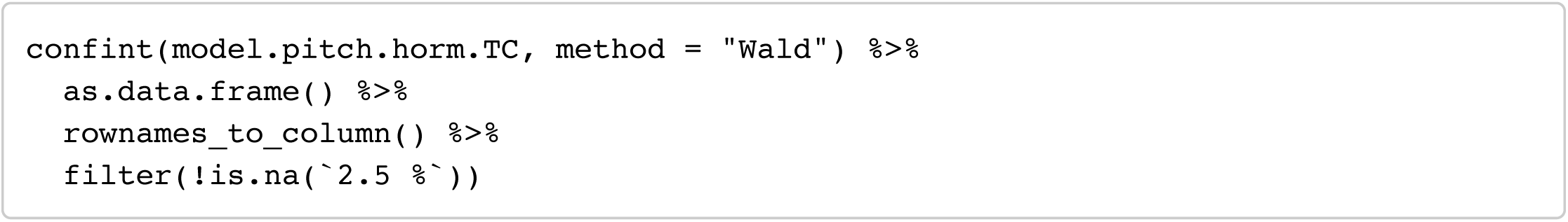

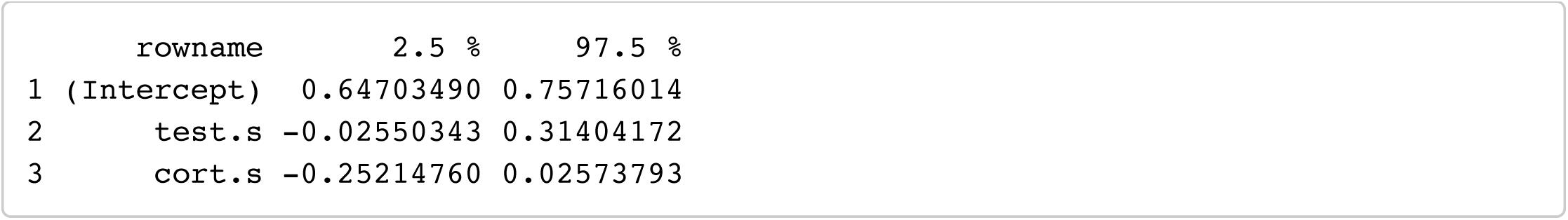

## Data subset: data_hormones_partner (with partnership status)

Only women with consistently reported partnership status for tests of effects of endogenous hormones considering effects of partnership status. Because women who reported their partnership status inconsistently are not included in partnership analyses, Ns for all partnership analyses will be slightly smaller

Hide

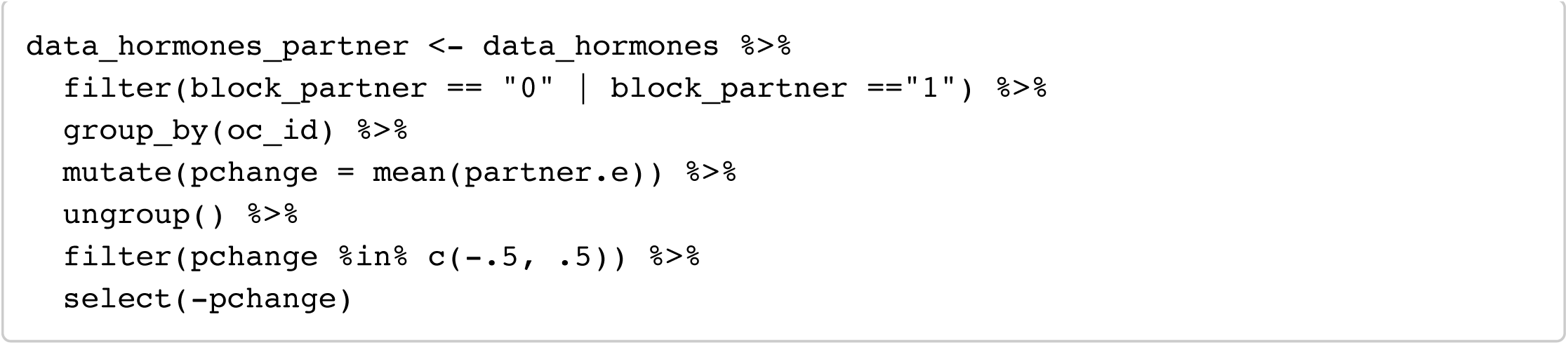

Hide

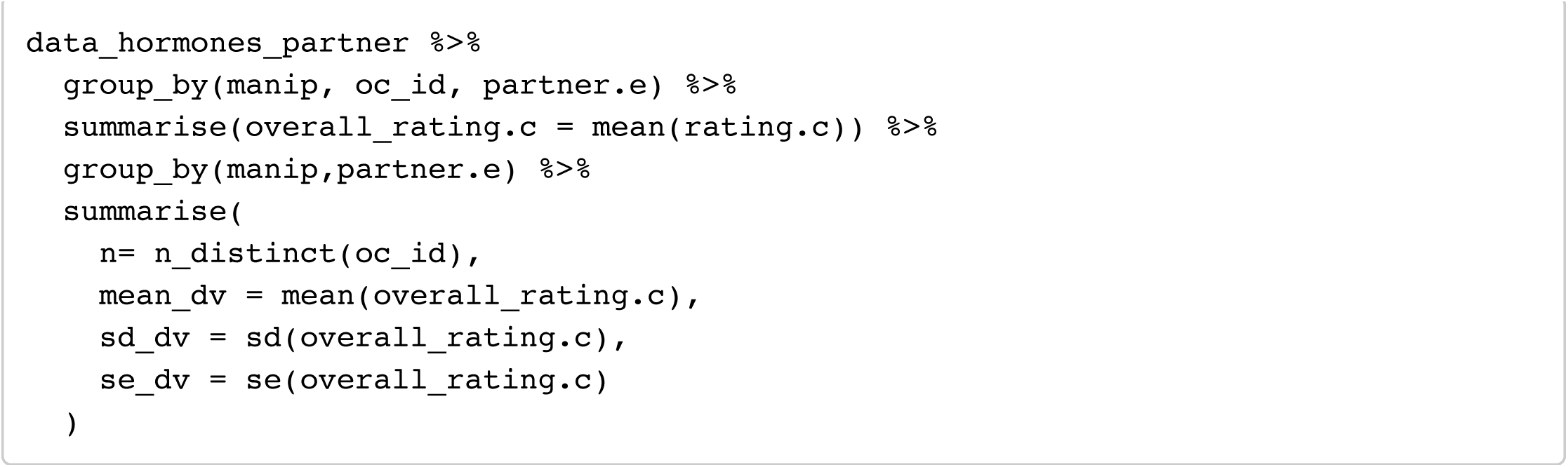

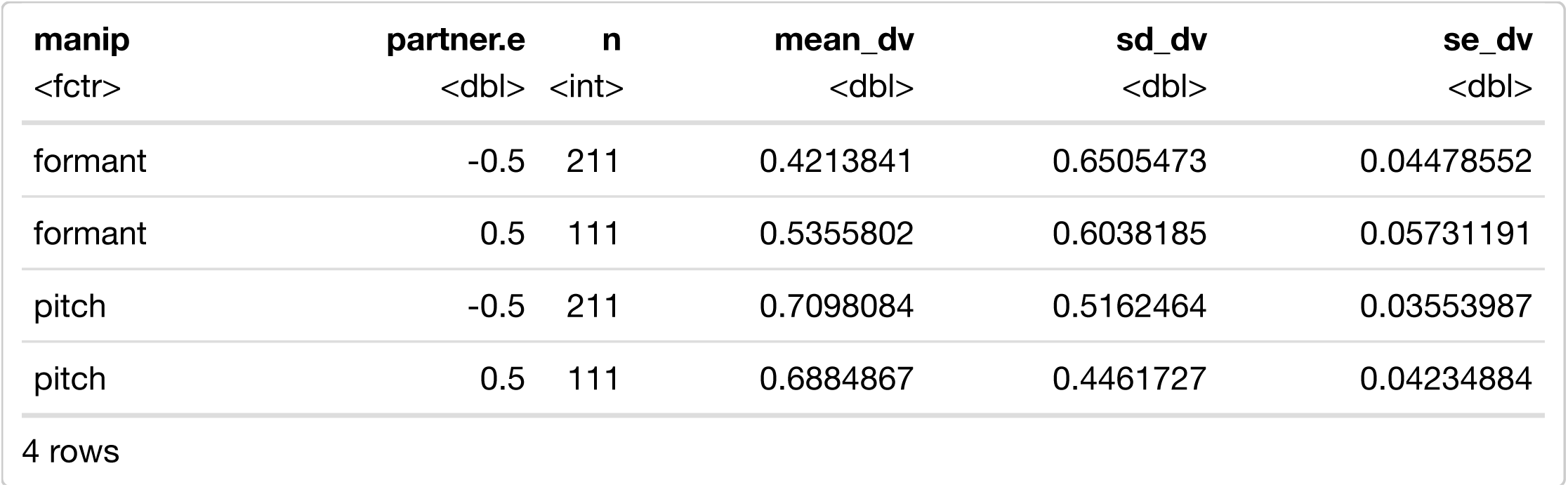

## Masculinity preference (formants)

### Model 1: formants ∼ E + P + E x P (with partnership status)

Hide

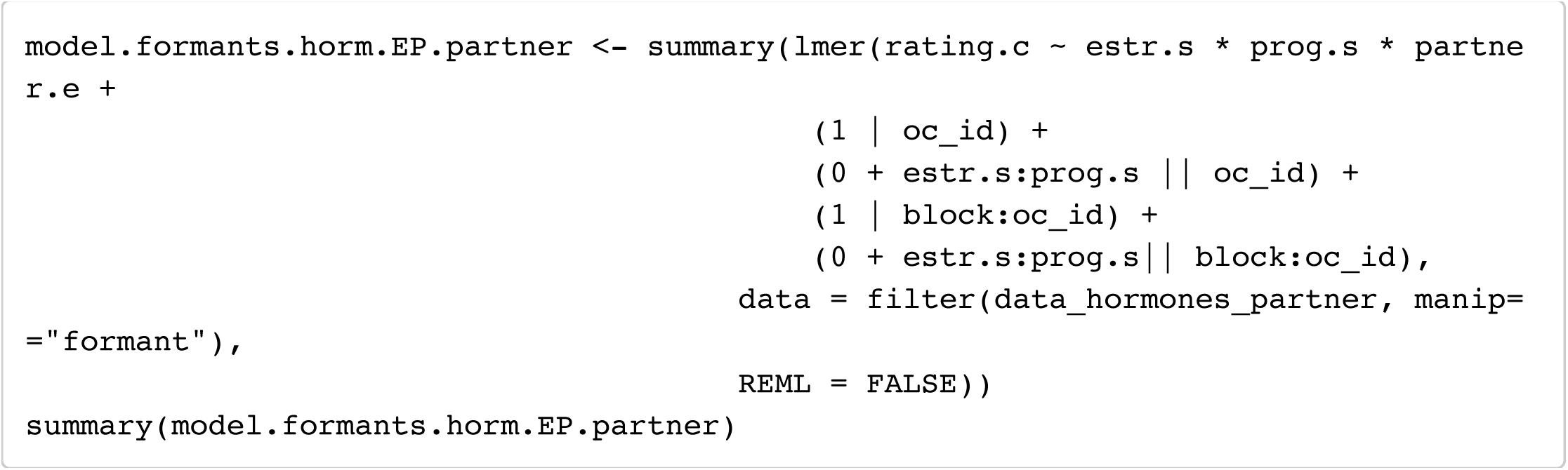

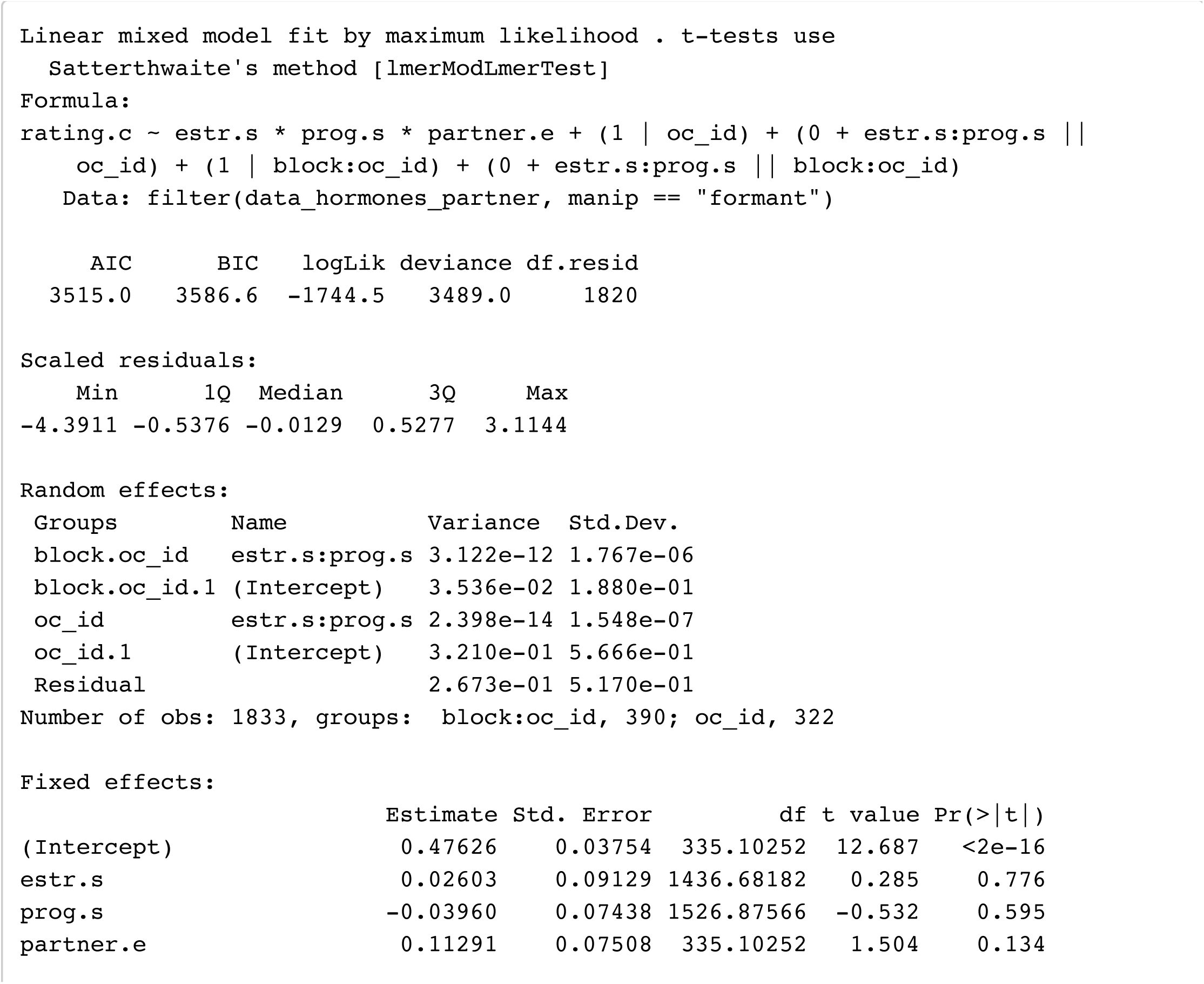

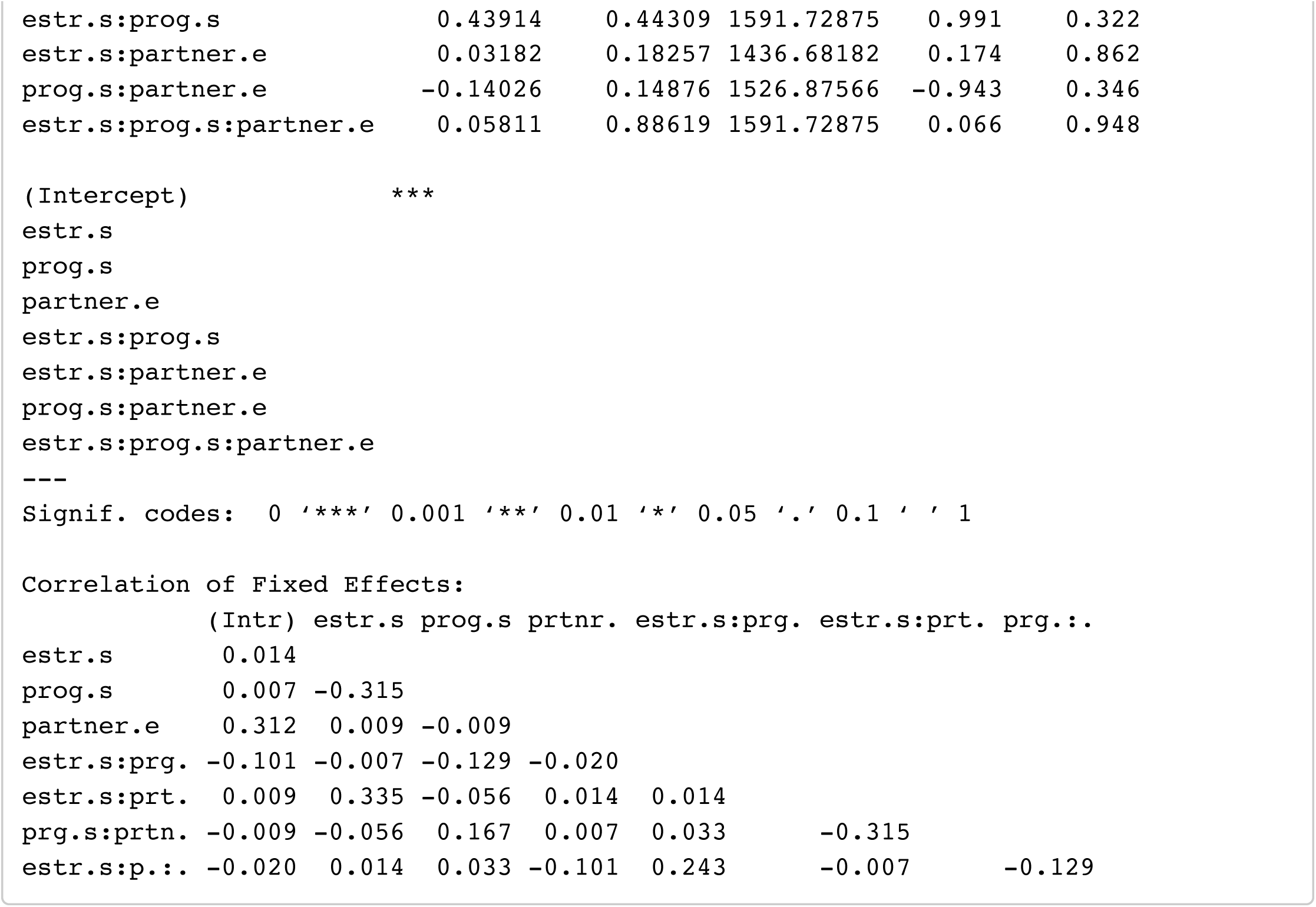

### Model 2: formants ∼ E + P + EP_ratio (with partnership

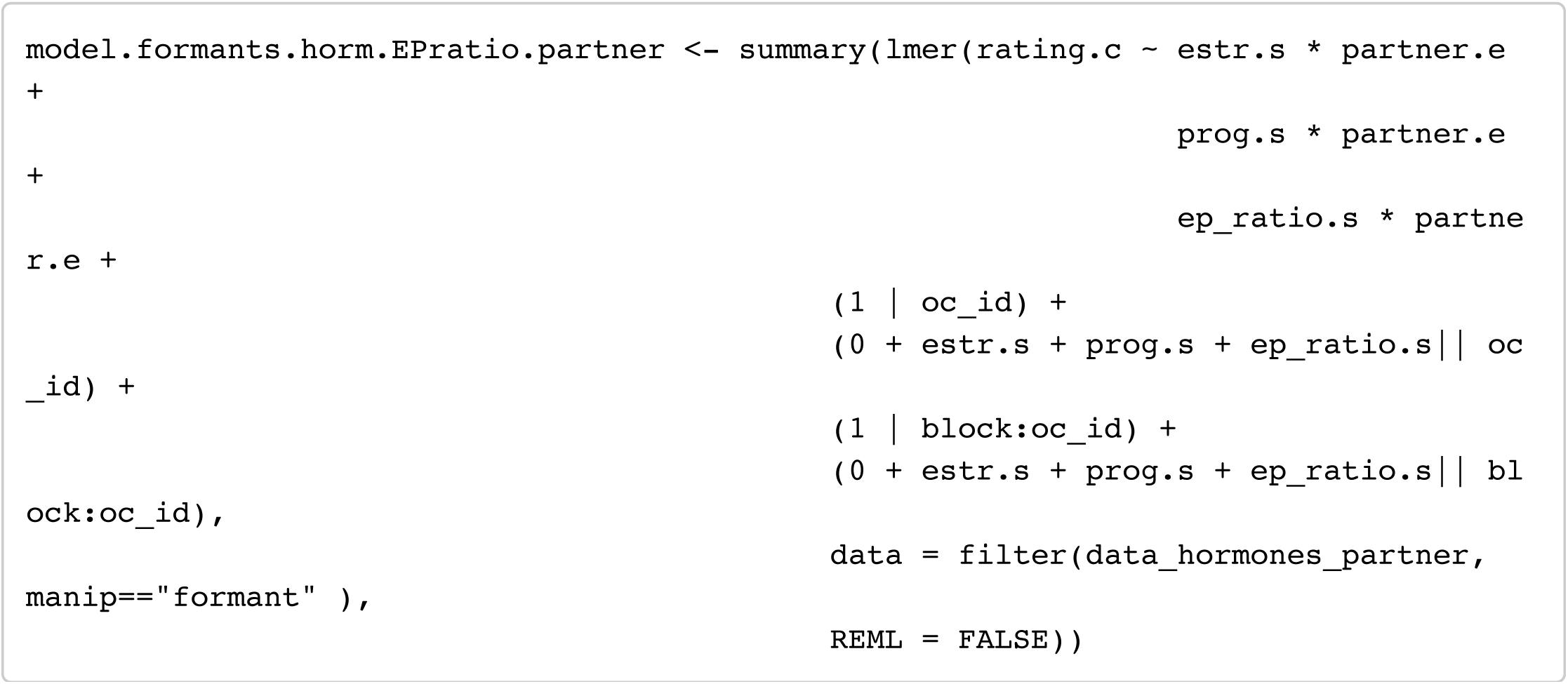

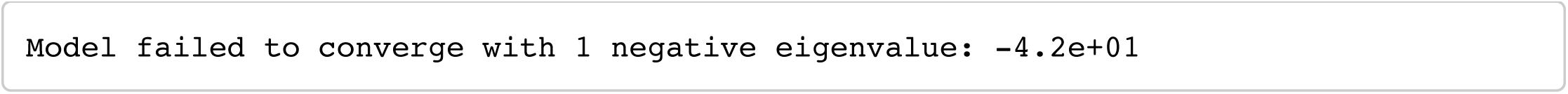

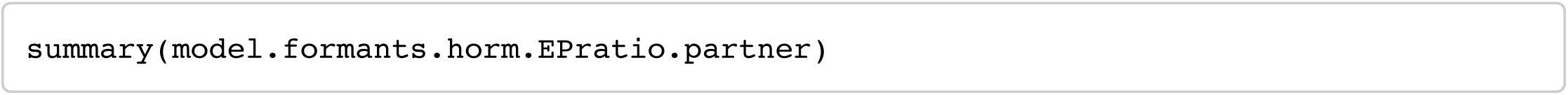

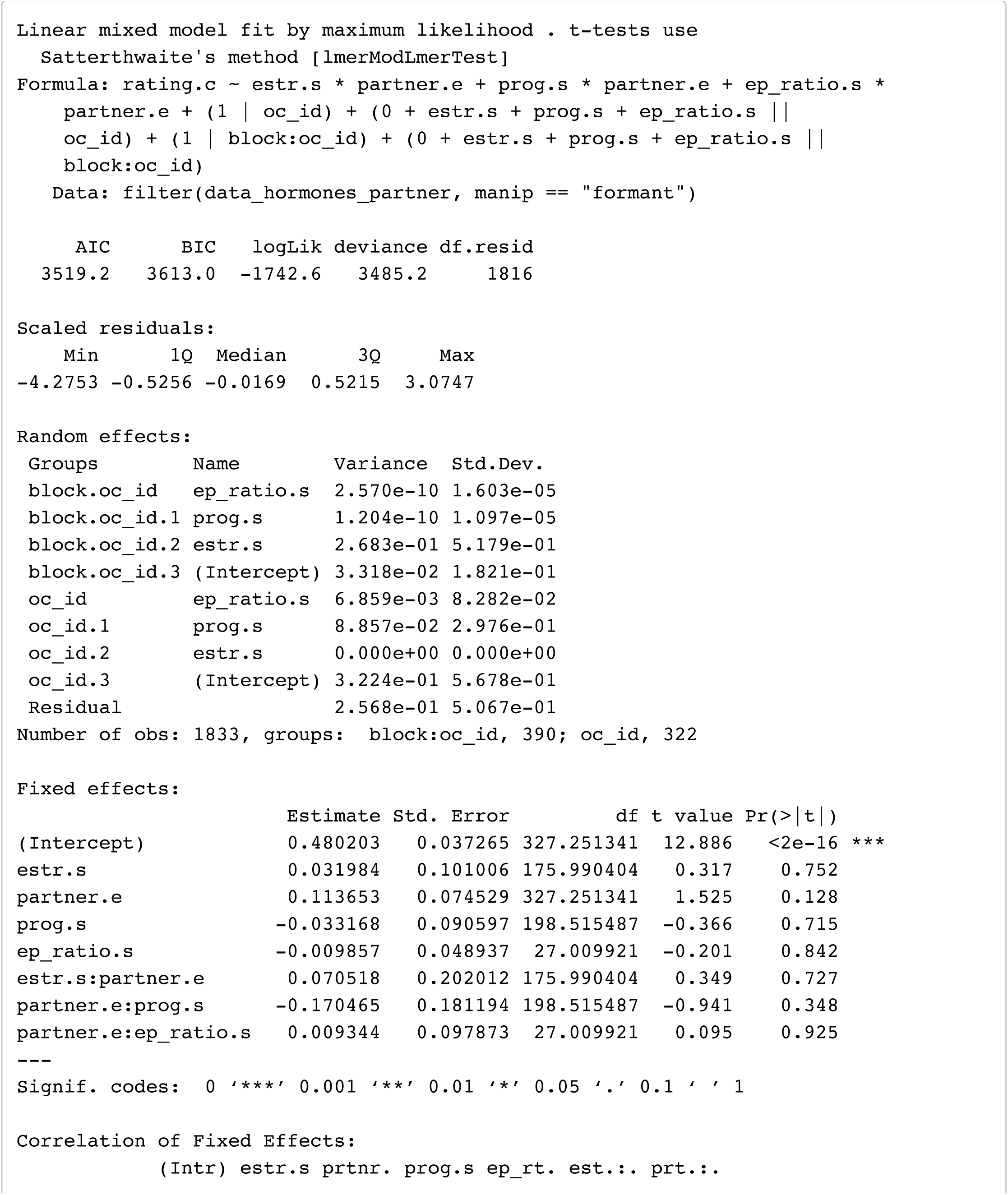

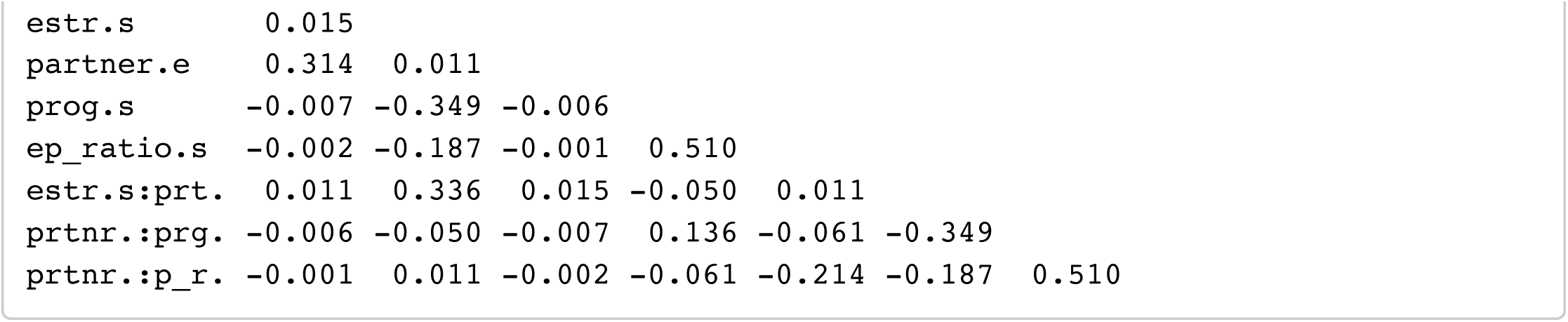

### Model 3: formants ∼ T + C (with partnership status)

Hide

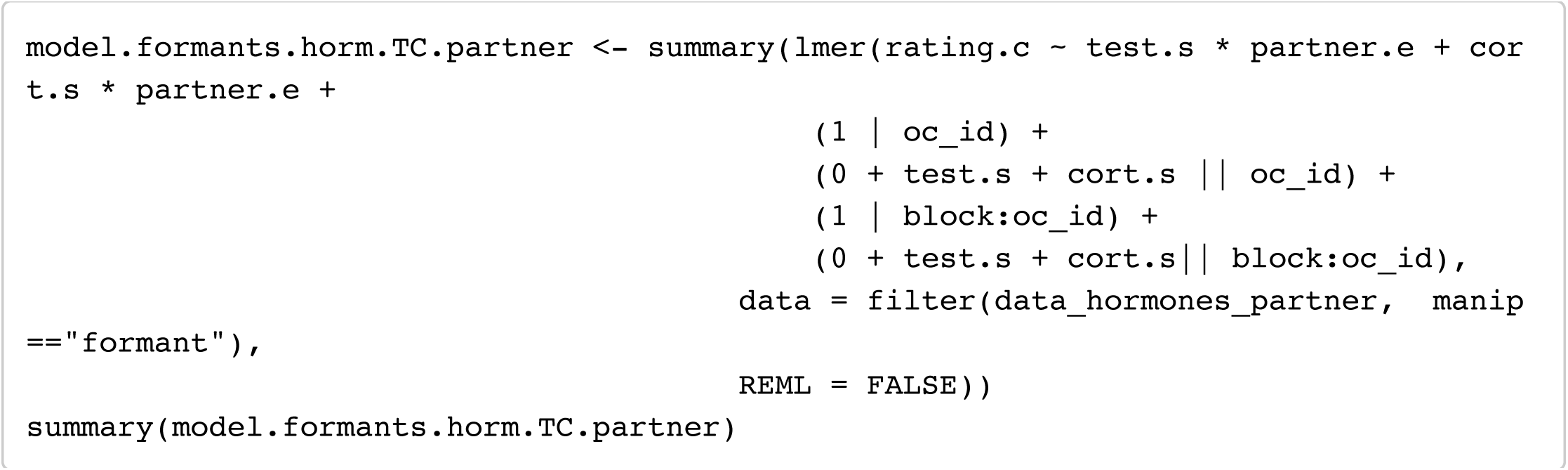

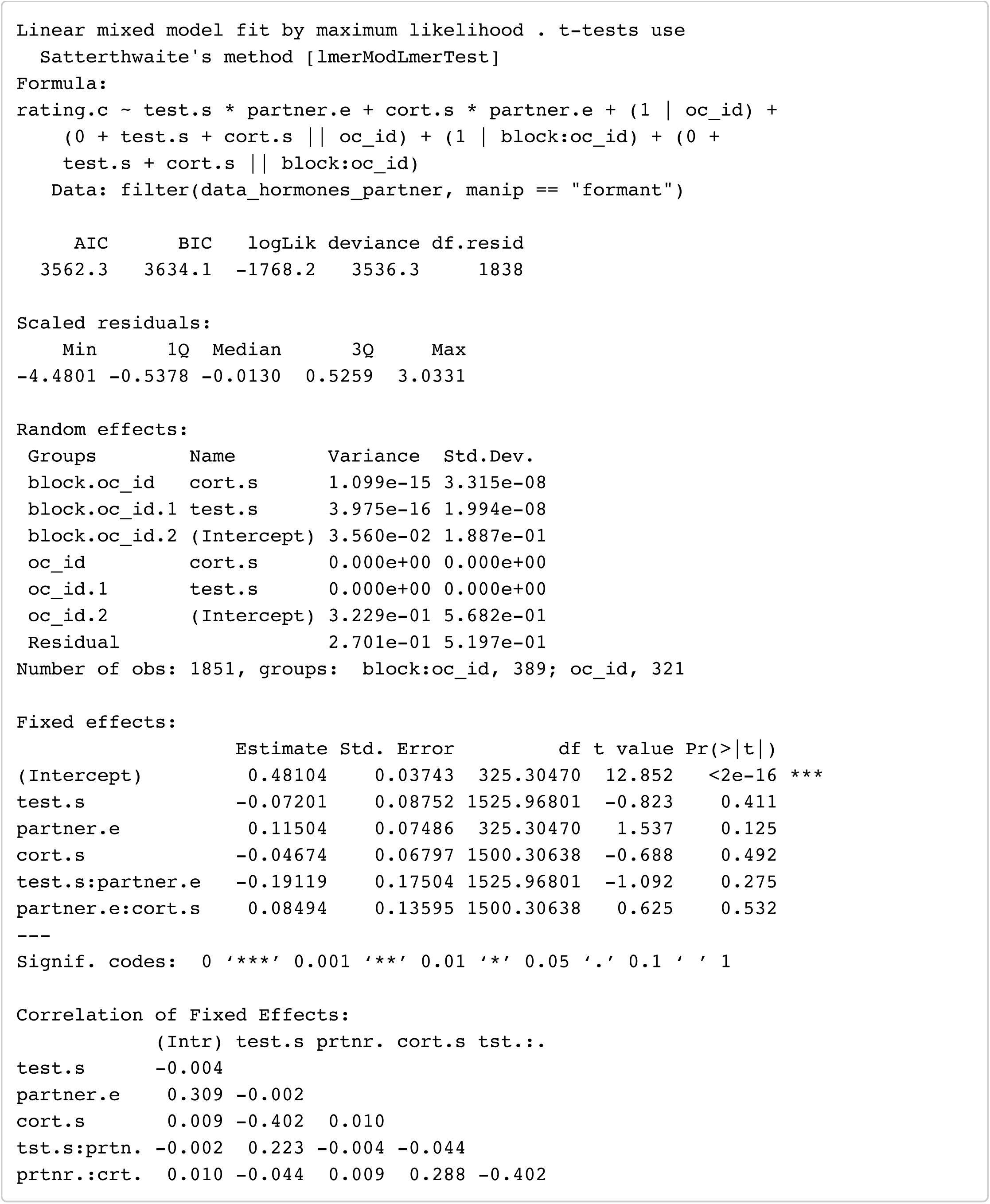

## Masculinity preference (Pitch)

### Model 1: Pitch ∼ E + P + E x P (with partnership status)

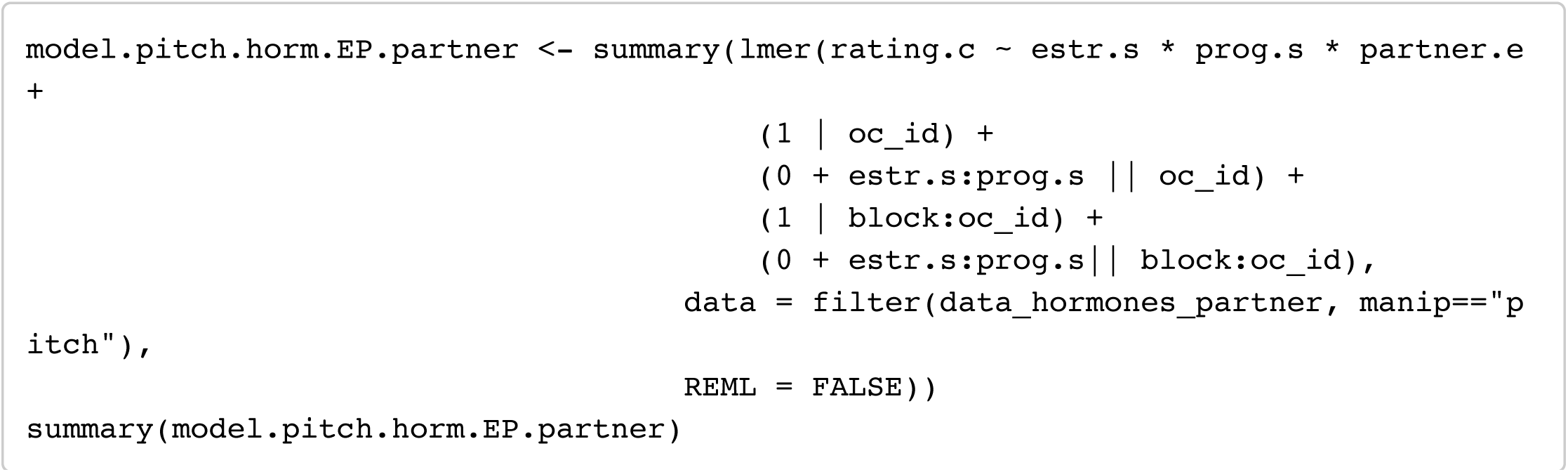

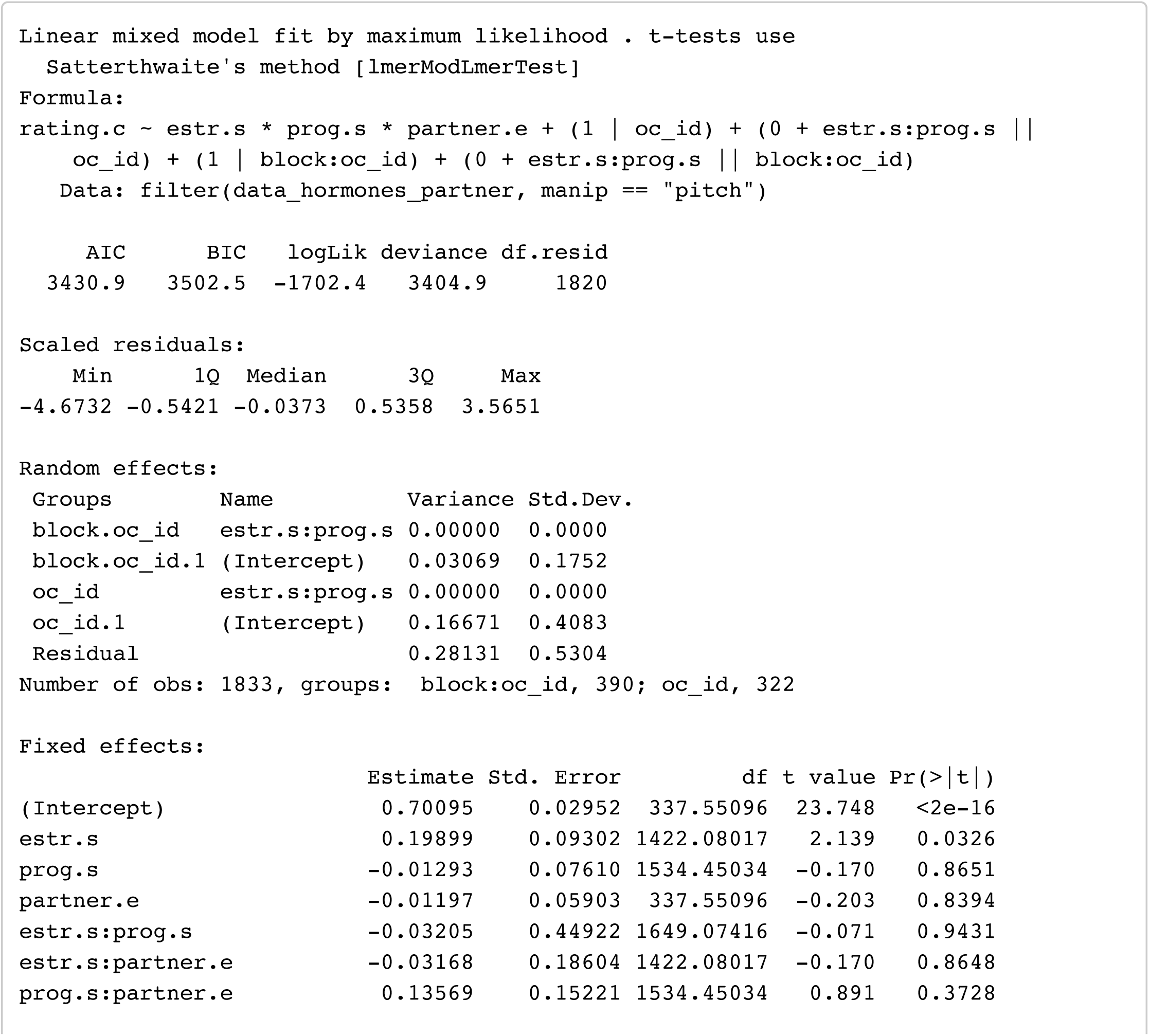

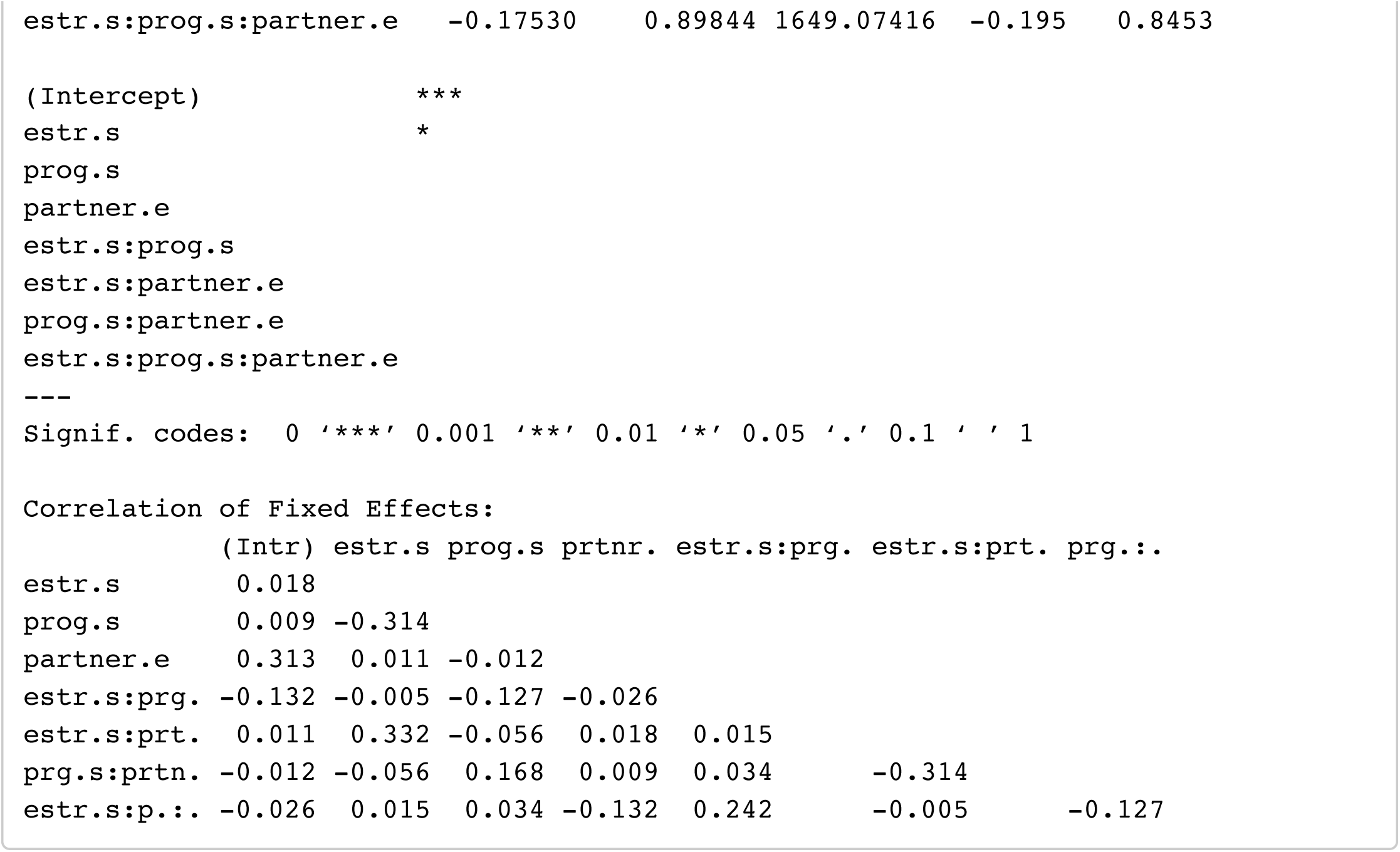

### Model 2: Pitch ∼ E + P + EP_ratio (with partnership status)

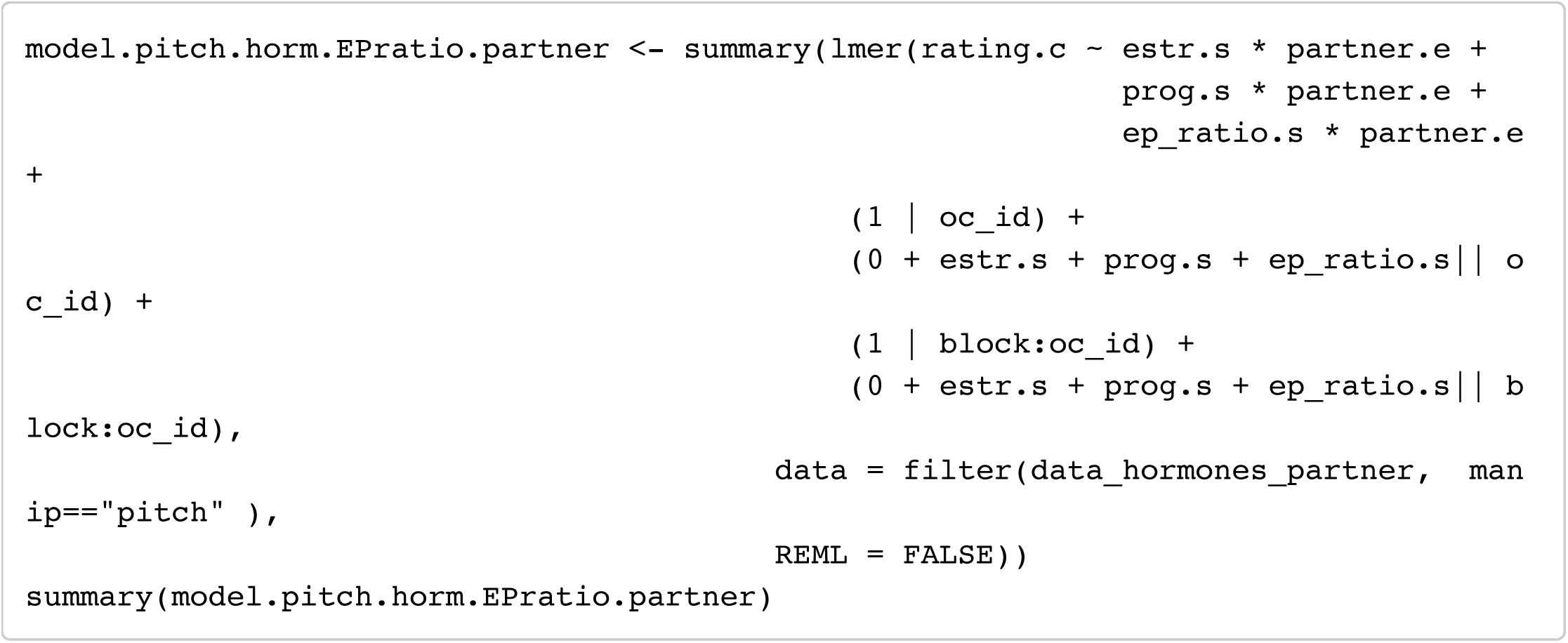

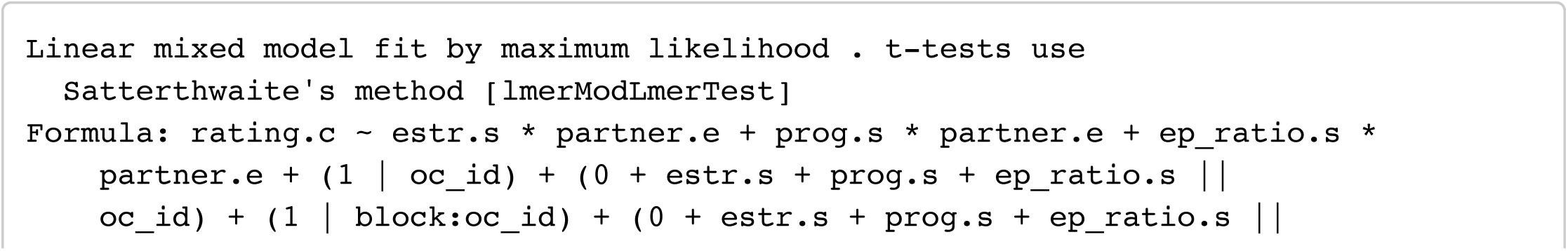

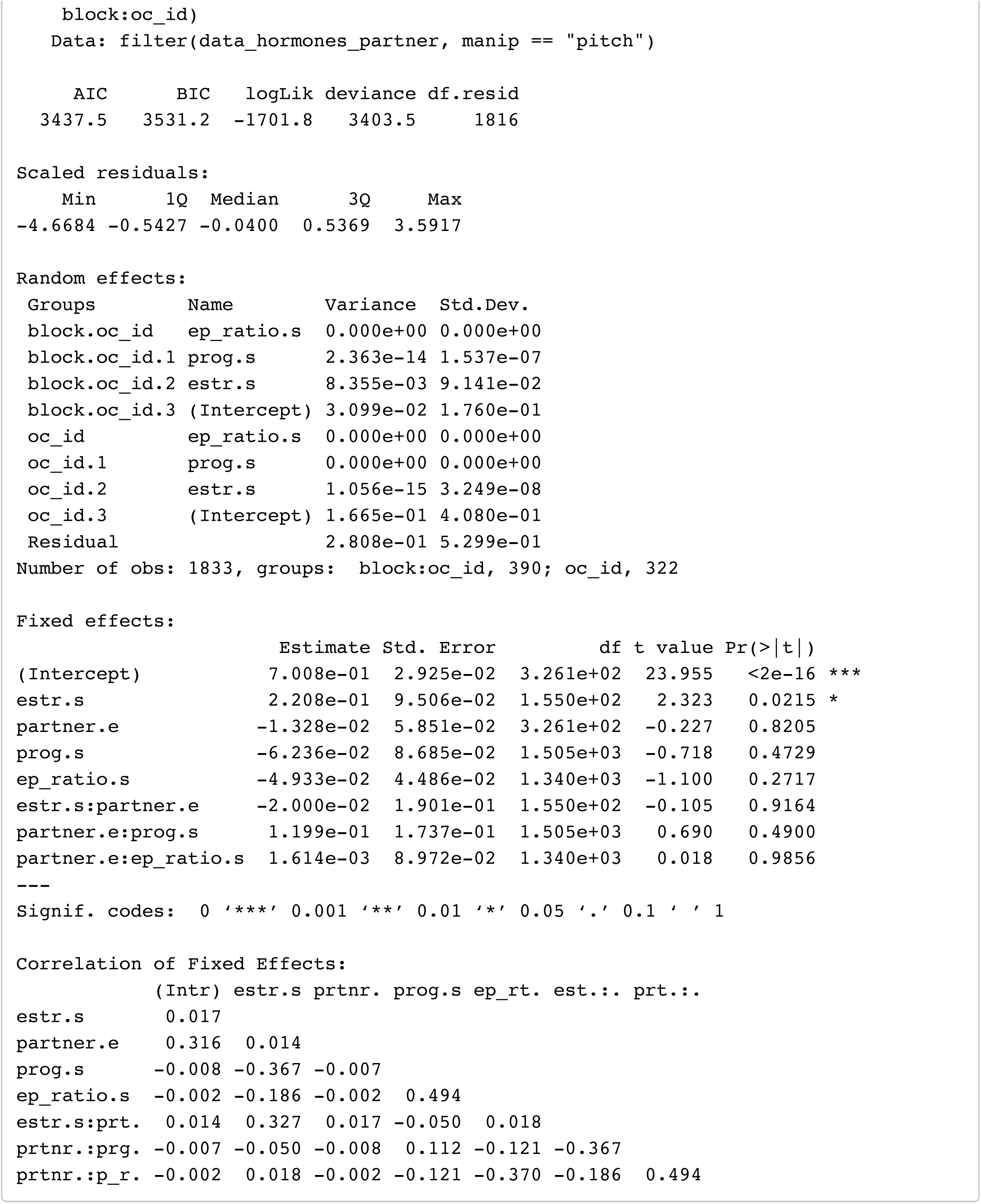

### Model 3: Pitch ∼ T + C (with partnership status)

Hide

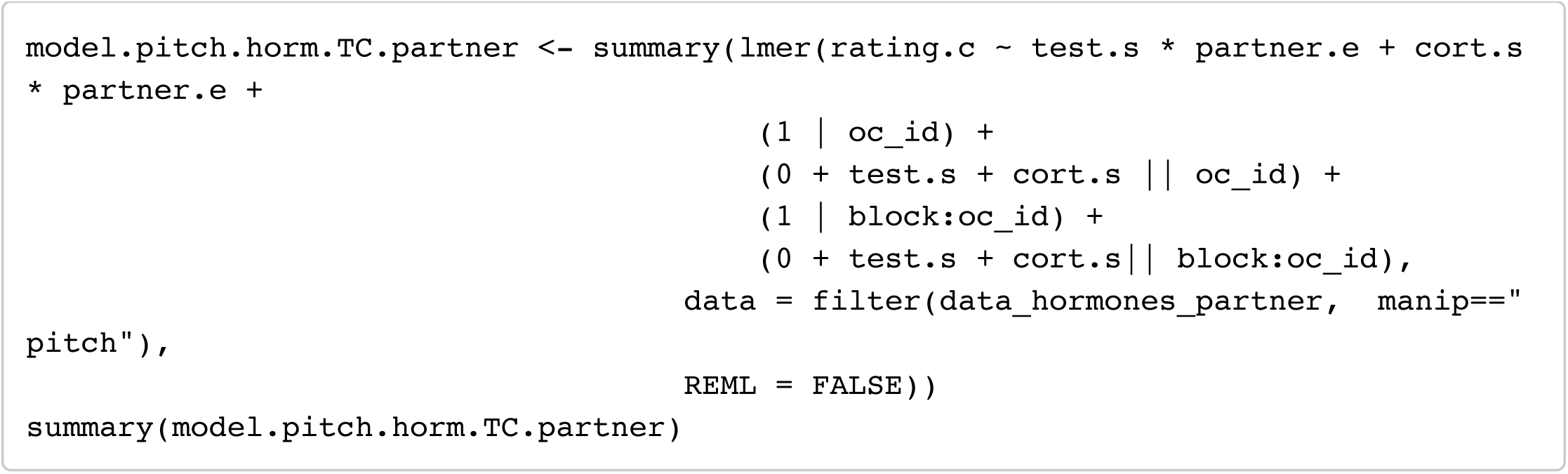

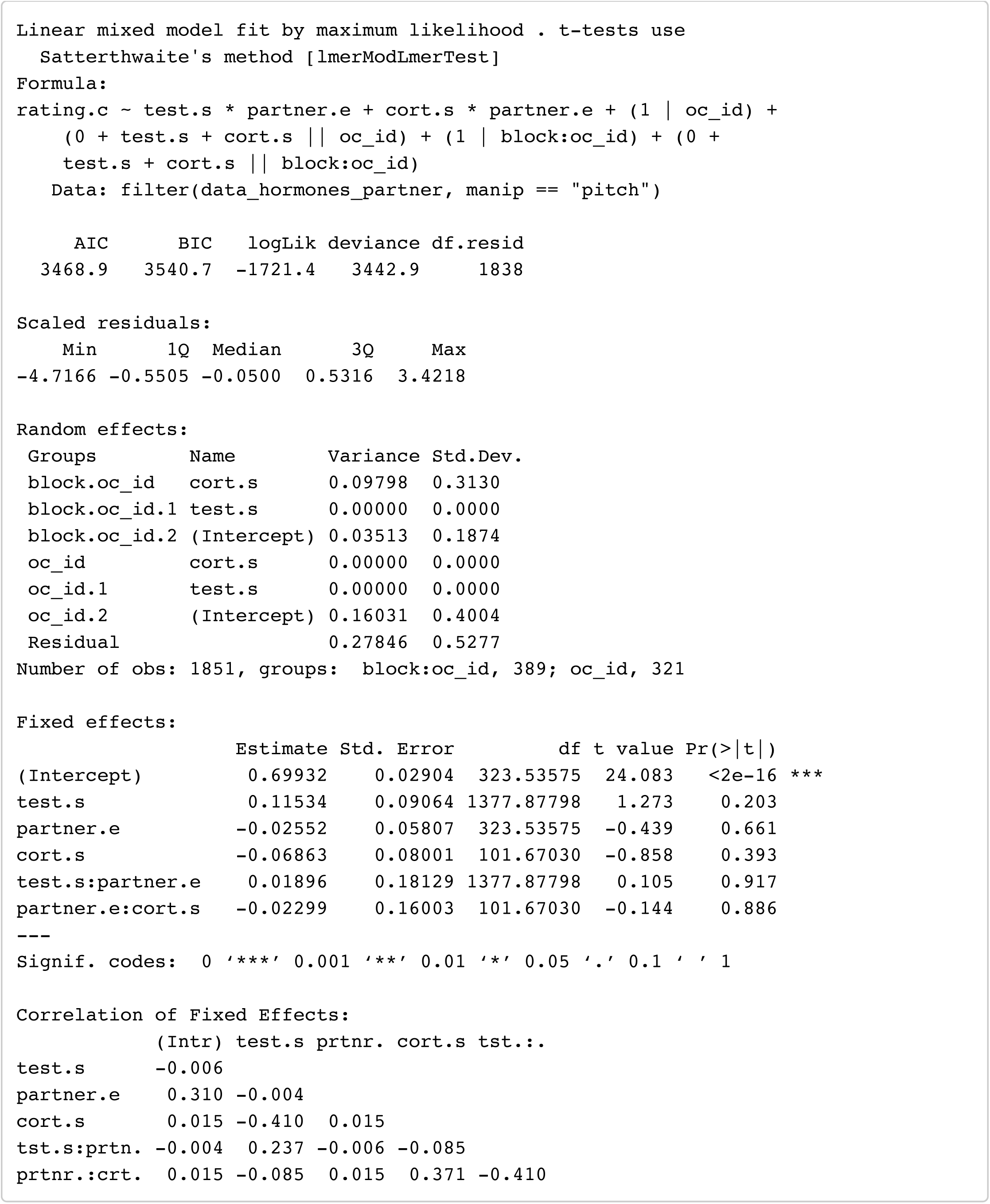

## Average Progesterone by Partnership Status

Testing whether the combined effects of partnership status and average progesterone predict masculinity preference.

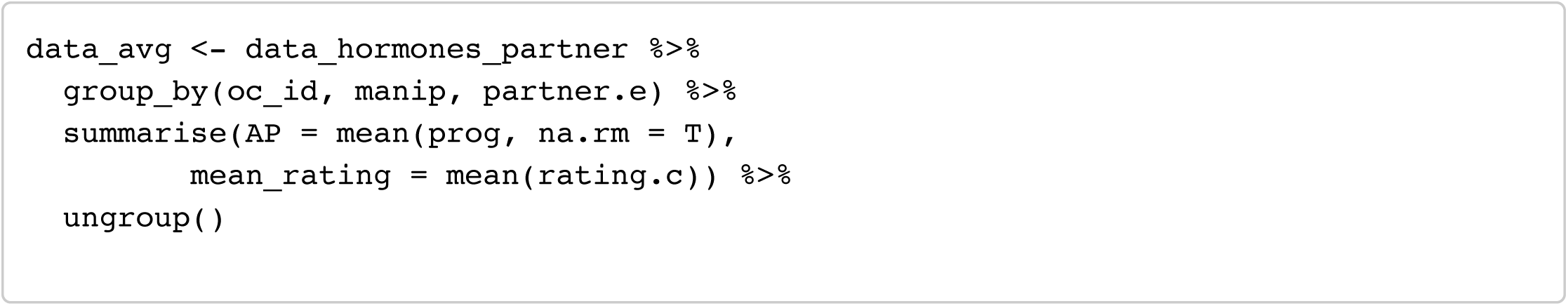

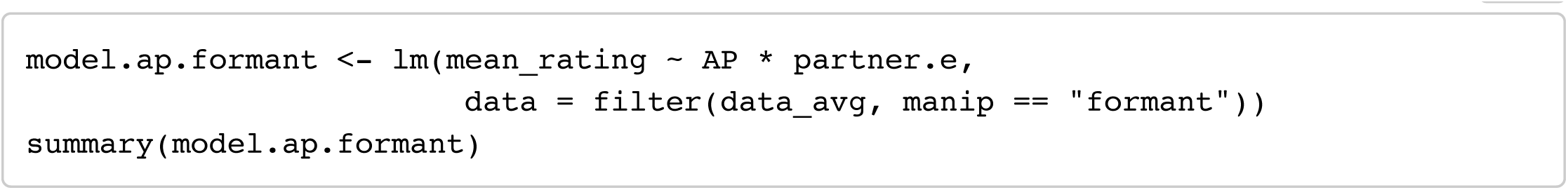

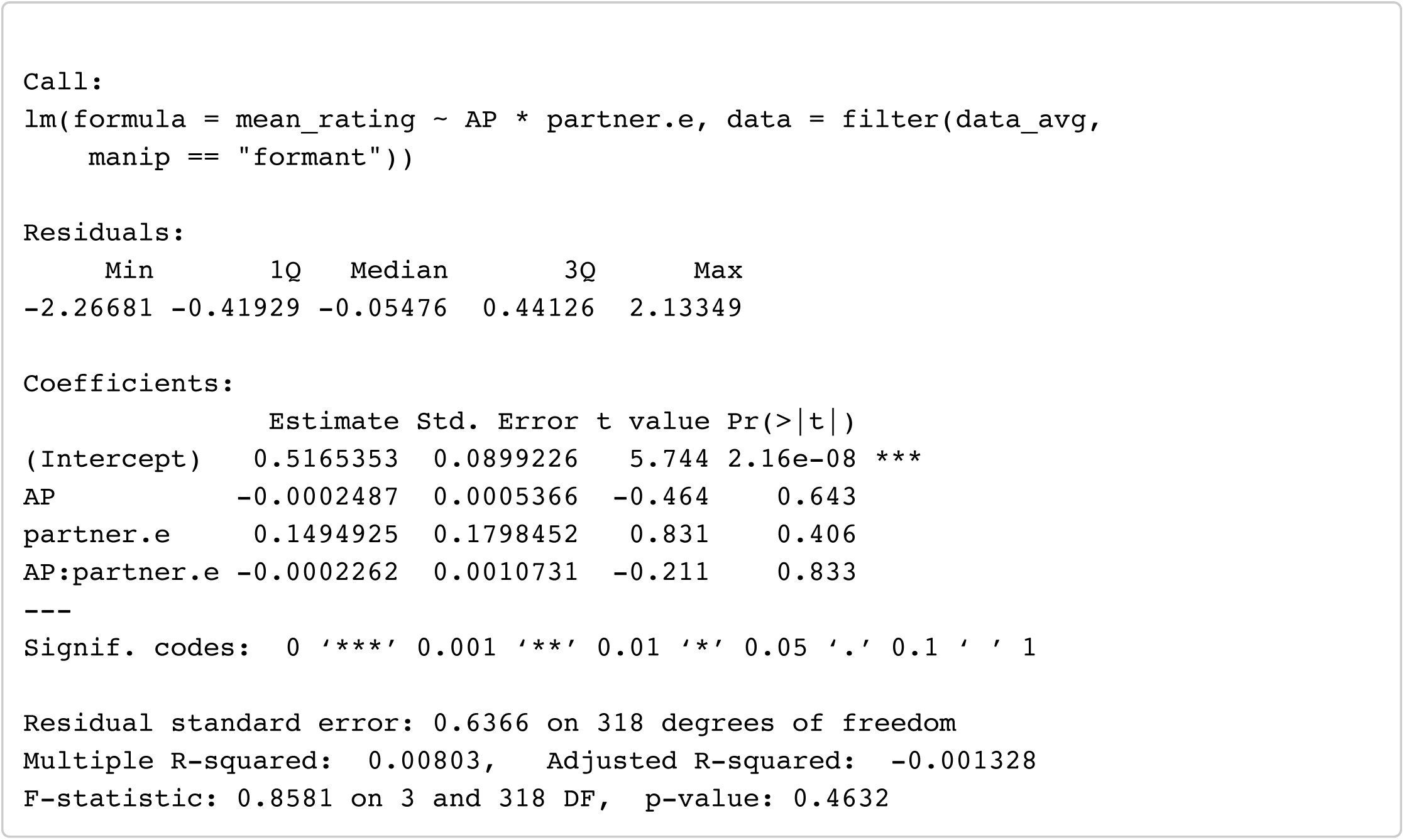

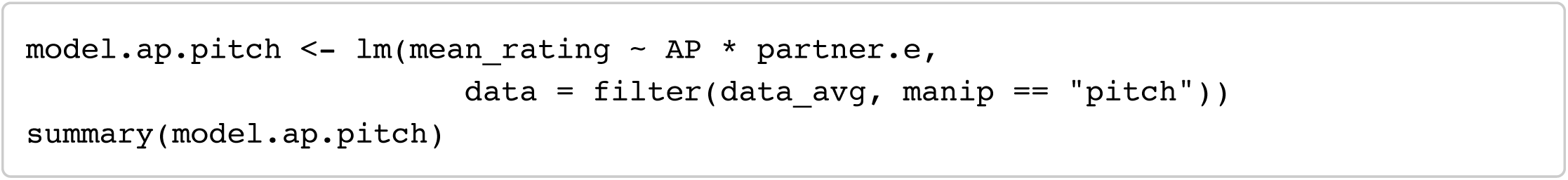

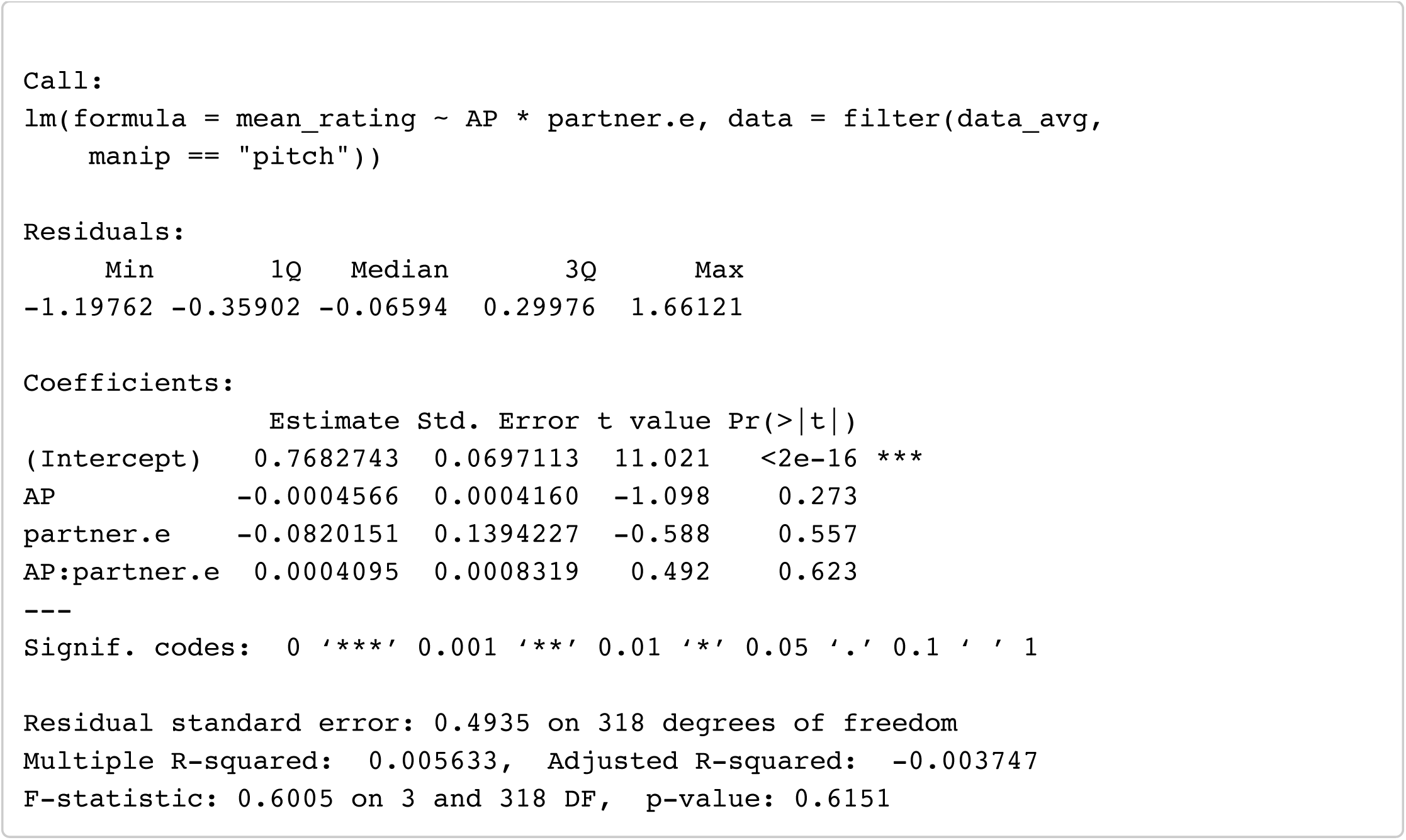

## Footnotes

1 p values for the effect of estradiol were .055 and .050, depending on the model.

